# Including tree spatial extension in the evaluation of neighbourhood competition effects in Bornean rain forest

**DOI:** 10.1101/2020.07.27.222513

**Authors:** David M. Newbery, Peter Stoll

## Abstract

Classical tree neighbourhood models use size variables acting at point distances. In a new approach here, trees were spatially extended as a function of their crown sizes, represented impressionistically as points within crown areas. Extension was accompanied by plasticity in the form of crown removal or relocation under the overlap of taller trees. Root systems were supposedly extended in a similar manner. For the 38 most abundant species in the focal size class (10 - <100 cm stem girth) in two 4-ha plots at Danum (Sabah), for periods P_1_ (1986-1996) and P_2_ (1996-2007), stem growth rate and tree survival were individually regressed against stem size, and neighbourhood conspecific (CON) and heterospecific (HET) basal areas within incremented steps in radius. Model parameters were critically assessed, and statistical robustness in the modelling set by randomization testing. Classical and extended models differed importantly in their outcomes. Crown extension weakened the relationship of CON effect on growth versus plot species’ abundance, showing that models without plasticity over-estimated negative density dependence. A significant negative trend of difference in CON effects on growth (P_2_ − P_1_) versus CON or HET effect on survival in P_1_ was strongest with crown extension. Model outcomes did not then support an explanation of CON and HET effects being due to (asymmetric) competition for light alone. An alternative hypothesis is that changes in CON effects on small trees, largely incurred by a drought phase (relaxing light limitation) in P_2_, and following the more shaded (suppressing) conditions in P_1_, were likely due to species-specific (symmetric) root competition and mycorrhizal processes. The very high variation in neighbourhood composition and abundances led to a strong ‘neighbourhood stochasticity’, and hence to largely idiosyncratic species’ responses. A need to much better understand the roles of rooting structure and processes at the individual tree level was highlighted.

## INTRODUCTION

One of most important advances in estimating and understanding dynamics of trees within forest communities was made when statistical analysis and population modelling moved away from the application of species- or guild-parameter averages and replaced them with spatially explicit estimates (DeAngelis and Mooij 2005, DeAngelis and Yurek 2017). Each and every individual was considered, its growth, survival and where possible its reproductive output, with reference to its neighbours. Neighbours are other trees close enough to the focal one to affect its resource acquisition and uptake (Pacala and Deutschman 1995, Pacala et al. 1996, Uriarte et al. 2004). Given that competition is a driving process of change in tree species abundance locally, differences in biomass, architecture and ecophysiological traits between focal trees and neighbours will in part be determining forest dynamics (Chen et al. 2018, 2019). Mean parameters often obscure differences between species, especially when variables are non-normally distributed and relationships are non-linear.

In addition to what can be measured and modelled deterministically, individuals from recruitment onwards are subject to demographic stochasticity affecting their survival (Engen et al. 1998, Lande et al. 2003). Environmental stochasticity in the form of climate variability (particularly rainfall and temperature) is thought to also play an essential role in forest dynamics (Vasseur and Yodzis 2004, Halley 2007). This form of stochasticity affects not only individual tree growth and survival directly, but also does so indirectly through its effects on neighbours and hence *their* competitive influence on the individual. Focal trees are simultaneously acting as neighbours to other ones nearby and reciprocal interactions operate. Due to these highly complicated and varying local-tree environments a form of what may be termed ‘neighbourhood stochasticity’ is realized. Any understanding of species-specific effects in neighbourhood modelling has, therefore, to cater for this inherently high system variability.

Within this conceptual framework of ongoing temporal and spatial variability, the role of neighbours on the growth and survival of small trees in tropical rainforests is analyzed more closely in the present paper. The data come from a long-term dynamics study at Danum in Sabah, NE Borneo. One motivation was to resolve better what constitutes conspecific (versus heterospecific) competition between trees; the other was to get closer to unravelling the role of below-ground processes, in the search for a mechanism. The new work builds on Stoll and Newbery (2005) and Newbery and Stoll (2013). To introduce the approach, it is first necessary to give the background of the previous Danum studies and modelling results to date, and then second to argue for the proposed extension, hypotheses and tests. As with all sites, data and model are context-dependent and contingent on site history. The principles behind the analysis should hopefully be relevant to other rainforest sites when making similar considerations.

### Current tree neighbourhood model

In the 10-year period of relatively little environmental climatic disturbance (P_1_:1986-1996) large trees of several species among the overstorey dipterocarps at Danum showed strong conspecific negative effects on the growth rates of juvenile trees in their immediate neighbourhood (Stoll and Newbery 2005). In the subsequent 11-year period (P_2_:1996-2007) which included an early moderately-strong El Nino Southern Oscillation (ENSO) event (April 1998), conspecific effects relaxed (Newbery and Stoll 2013). If the effects of the first period were a result of intra-specific competition, perhaps principally for light, then the dry conditions caused by the event in the second, which temporarily thinned the overstorey foliage and markedly increased small twig abscission (Walsh and Newbery 1999), would have allowed more illumination to the understorey, and hence ameliorated the earlier P_1_ conspecific effects. However, that conspecific effect could be for light presents a problem for two reasons. One is that heterospecific negative effects appeared to be much weaker or non-operational in P_1_ (Stoll and Newbery 2005), and the other is the difficulty of explaining a competitive effect for light (i.e. a mechanism of shading) that is species-specific. Large trees will presumably shade smaller ones regardless of their taxonomic identity, although responses to shading by affected trees might differ between species due to their physiologies. The two periods of measurement may also have differed in other respects besides intensity of drought stress and light changes, and these remain unrecorded or unknown. In terms of succession, the forest at Danum also advanced between P_1_ and P_2_ although still remaining within the late stage of its long-term recovery from a historically documented period of extensive dryness in Borneo in the late 19^th^ century, with tree basal area continuing to rise and overall tree density decreasing (Newbery et al. 1992, Newbery et al. 1999, Newbery and Stoll 2013).

The hypothesis advanced by Stoll and Newbery (2005) was that interactions below ground may primarily have been causing the conspecific effects for dipterocarps, in the form of competition for nutrients combined with, or enhanced by, host-specialist ectomycorrhizal (ECM) linkages between adult and juvenile trees within species. This would be particularly relevant for species of the Dipterocarpaceae, the dominant tree family in these forests, and which accordingly have the highest neighbourhood basal areas associated with the strongest conspecific effects. In stem size, focal juveniles were 10 − 100 cm girth at breast height, *gbh* (1.3 m above ground, equivalently ∼3 – 30 cm diameter, *dbh*), and were therefore well-established small-to-medium trees in the understorey and lower canopy (Newbery et al. 1992, Newbery et al. 1996). Compared to these small trees with their lower-positioned shaded crowns, the higher demands of the large well-lit and fast-growing adults above them may have been making relatively high demands on soil nutrients, and thereby drawing these resources away from the juveniles. As a result, the slowed juvenile stem growth may have been due to root competition, enhanced possibly by ECMs. Increased light levels in P_2_, even moderately and temporarily in 1998-99 (Walsh and Newbery 1999; Newbery and Lingenfelder 2004, 2009), likely allowed suppressed conspecific juveniles to attain higher growth rates than those in P_1_ (Newbery et al. 2011). It was postulated that that was in part or wholly caused by smaller trees reversing the nutrient flow back from the larger ones, i.e. from juvenile to adult (Stoll and Newbery 2005).

The extent and nature of any ECM linkages and the changing nutrient flows have not been experimentally demonstrated for this forest, so the nutrient hypothesis is tentative. It is difficult to conceive, though, of another mechanism that could explain the results, at least in physical and physiological terms. Against the hypothesis though is broader evidence that the degree of host specialism for ECM fungi in the dipterocarps may be weak because most dipterocarps appear to have many fungal species in common (Alexander and Lee 2005, Brearley 2012, Peay et al. 2015). Whilst generalist ECMs were recorded mainly for seedlings and some adults, small-to-medium sized trees might have been more strongly linked to adults via specialist ECMs, in a period of tree development when dependence on ectomycorrhizas for nutrient supply would be more important than in the earlier ontological stages. This differentiation would be particularly relevant for overstorey dipterocarps. Whilst it is quite possible that carbon moves though a general mycorrhizal network linking adults to seedlings when the latter are really very small and in deep shade (Simard et al. 2002, Simard and Durall 2004, Selosse et al. 2006) it does not mean necessarily that generalist ECMs would function in this same way after the sapling stage, as the small trees became gradually more illuminated. It is also feasible that the element most important for tree interactions changed over time from carbon to phosphorus as the nature of the ECM symbiosis switched from being generalist to specialist.

An alternative hypothesis is that conspecific effects as such were happening ‘by default’ (Newbery and Stoll 2013). Because, in some species, adults and juveniles tend to be spatially clustered due to the limited distances to which especially dipterocarp seeds are dispersed, conspecifics often made up most of the large-tree adult neighbour basal area around a focal juvenile. Conversely, some species lacked aggregations possibly because, where more-scattered juveniles now survive, the parents had recently died. Dipterocarps, and other overstorey species, show a wide range of aggregation at different scales (Newbery et al. 1996, Stoll and Newbery 2005, Newbery and Ridsdale 2016). Compared with a forest in which trees might theoretically be all distributed at complete randomness, one with aggregations would result in proportionally more trees of the same species (conspecifics) rather than different ones (heterospecifics), occurring at close distances. This fact would tend to an explanation of conspecific effects based on one common mechanism (such as shading); and the effect of the ENSO disturbance in P_2_ was to release understorey small trees of all species, to differing degrees depending on each species’ degree of responsiveness to light increases. The role of ECM linkages and nutrient flows would then become secondary, operating as a consequence of light effects (Newbery and Stoll 2013). Several overstorey species in the P_1_-P_2_ comparison were not dipterocarps however (presumably they had no ECMs) yet they still showed strong conspecific effects in P_1_, which were relaxed in P_2_ (Newbery and Stoll 2013). Possibly these other species with strong CON effects were endomycorrhizal and had similar degrees of specialism like those with ECMs. Strength of conspecific effect was furthermore not convincingly related to degree of spatial clustering within the dipterocarps (Stoll and Newbery 2005). The two resource-based hypotheses, ‘light’ versus ‘nutrients’, were not readily separable, and an extended approach was needed to better distinguish between them.

### Extending the neighbourhood model

Modelling attempts to date have mostly taken basal areas of neighbors around focal individuals defined by the radial distances between centres of tree stems, normally weighting each neighbour tree’s basal area by the inverse of distance (Canham et al. 2004, 2006; Canham and Uriarte 2006). Whether a tree was inside a circle of a given radius or within a 1-m annulus, or not, depended solely on the coordinates of its centre as a point distribution: focal and neighbour trees had no spatial extent. Competitive influences and ECM networking might therefore be more realistically represented by the allometric extension of crowns and root systems in the form of a zone of influence, or ZOI (Bella 1971, Ek and Monserud 1974, Gates and Westcott 1978, Pretzsch 2009). Zones would overlap in ways that simulated better resource allocation and in doing so conspecific effects in P_1_ would be expected to increase and differences in effects between P_1_ and P_2_ to generally strengthen.

The zone of influence concept must be recognized from the outset as a simplistic one in that it assumes that trees in their manner of influencing neighbours were above- and below-ground contiguous matching cylinders (Schwinning and Weiner 1998, Weiner et al. 2001, Stoll et al. 2002, Weiner and Damgaard 2006). The notion of similarity of light and nutrient competition strengths is likely not realistic, especially when there are differences between species in root-shoot allocation ratio and essentially very different mechanisms of competition are involved (Newbery et al. 2011, Newbery and Lingenfelder 2017).

Crown area has been found to be generally strongly positively correlated with stem diameter in studies of tropical tree architecture and allometry (e.g., Bohlman and O’Brien 2006, Antin et al. 2013, Blanchard et al. 2016, Cano et al. 2019). Zambrano et al. (2019) have recently explored using nearest neighbour models with crown overlap in relation to functional traits. Whilst above- and belowground effects will not be independent of one another for structural and physiological reasons, there is no direct evidence in the literature to suggest that lateral spread of root systems mirrors canopy shape and extent. As a start, a ZOI could be envisaged as being made up of many constituent points, symbolizing *plant modules* (branch ends with leaves, coarse and fine roots), so that points within focal trees’ zones, and those of their neighbours, would be at many various distances from one another (Sorrensen-Cothern et al. 1993, Pretzsch et al. 2015). Crowns would be expected to show some plasticity and to relocate themselves in space to achieve at least maximum light interception (Purves et al. 2007, Strigul et al. 2008). Roots can be also plastic, and maybe more so than crowns as they are without mechanical support constraints and are more exploratory in their search for nutrients.

Changing from a ‘classical’ non-spatial-to spatial-extension models might then be a way to distinguish between the two hypotheses. Spatial extension should lead to the detection of a stronger conspecific effect because any step that let tree size more closely represent the mechanical process of competition would presumably reinforce that effect. This is more immediately obvious when considering crown sizes and light interception: larger trees with larger crowns would shade larger areas of neighbours than smaller ones. But would this apply in the same way to roots below ground, where root systems of large trees, and their ECMs interlink more often with those of neighbours than do the root systems of smaller trees? Indeed, areas occupied by roots are usually quite heterogeneous in shape, and roots of difference sizes at different distances from trees have differing uptake capacities. A principal difference, therefore, between above-ground competition for light and below-ground competition for nutrients is that the former is probably almost entirely asymmetrical in nature and the latter in the main symmetrical (Weiner 1990, Schwinning and Weiner 1998).

Under symmetrical competition for resources, uptake and utilization by neighbours are linearly related to their biomass (proportionate), and under asymmetrical competition they are non-linearly, normally positively, related to biomass (disproportionate redistribution). These definitions do not exclude competition below ground between roots being slightly asymmetric too under some conditions, though the degree of asymmetry is likely to be far less than that for light above ground as the latter is one-directional and instantaneous in use and the latter three dimensional and gradual. The two forms of symmetry correspond to the removal of the smaller tree’s resources (exploitive non-redistribution) and to relocation of its resources (proportional or shared redistribution) as model modes. Models that fit better with removal form might suggest a predominance of light competition, ones that fit better with relocation, a predominance of nutrient competition. Higher competition above ground will partly translate to higher competition below ground, and vice versa due to root-shoot inter-dependencies. On the other hand, a root-shoot allocation strategy and plasticity could counteract that translation. If spatial models in either form failed to improve model fitting this might question whether competition for resources is at all a reason for the conspecific effects or invoke a search for why the model alternatives were not correctly representing envisaged neighborhood interactions.

If ECMs in general contribute to enhancing root competition, conspecific effects of neighbours under spatial extension models should furthermore be higher for dipterocarps than non-dipterocarps, especially under the relocation form — for neighbouring trees of similar sizes (basal areas). A mixed range of increases in effects might indicate specialist fungi operating more in favor of some host species than others. If an ECM network operates it can be postulated that the distance effect will not be mirroring resource depletion curves around trees, but be allowing exploration to much further away. Conspecific effects below ground would presumably operate most strongly in species that are strongly aggregated, not necessarily in that case requiring specialist ECMs; but for dipterocarps that are more spread out they would lack the immediate advantage of high local abundance and ex hypothesis the one way left for them to affect juveniles conspecifically would be through ECMs. Non-aggregated species would be expected to have greater releases in growth rates than aggregated ones, being much freer of adult influences at distance.

### Context and modelling aims

In the context of the nearest-neighbour modelling explored in this paper, terms ‘spatial’ and ‘non-spatial’ refer to the spatial *extension of crowns* around point stem locations. Using crown sizes constitutes a spatial model, using only stem centre locations constitutes a non-spatial one. This usage should not be confused with the one of statistical spatial point (pattern) analysis. ‘Spatial’ is a fundamental physical attribute that is applied in numerous contexts.

Unravelling the causal nexus of system interactions (direct and indirect effects, reciprocation and feedback, time-lagged) is very complicated if the aim is to reduce a phenomenon such as the *average* conspecific effect of a species at population and community levels to a set of understandable mechanisms operating between individuals in space and time (Clark 2007, Clark et al. 2010, Clark et al. 2011). Conspecific effects, if they are indeed real, and not ‘by default’, might play a role in determining species composition in forests, but they do not necessarily need to be competitive or facilitative if they come about from a combination of spatial clustering (caused by dispersal) and stochastic environmental (climatic) variability (Newbery and Stoll 2013).

This third concluding paper on the role of neighbourhood effects on tree growth and survival in the lowland rain forest at Danum in Sabah builds directly on Stoll and Newbery (2005) and Newbery and Stoll (2013) by incorporating spatial extension to trees. It attempts to (a) reject the ‘default hypothesis’ for conspecific effects in favor of a resource-based competition one, and where successful (b), reject the hypothesis that conspecific competition is largely for light in favor of the alternative that it is more for nutrients. This leads to a revision in how negative density dependence is seen to operate in tropical forests and its role in tree community dynamics, as well as a reconsideration of neighbourhood stochasticity.

## MATERIALS AND METHODS

### Study site

The two permanent 4-ha plots of primary lowland dipterocarp forest just inside of the Danum Valley Conservation Area (Sabah, Malaysia), close to the middle reaches of the Ulu Segama, are situated *c*. 65 km inland of the east coast of Borneo, at 4° 57’ 48” N and 117° 48’ 10” E. They are at c. 220 m a.s.l.; measure each 100 m x 400 m in extent, lie parallel c. 280 m apart: each samples the lower slope-to-ridge gradient characteristic of the local topography. Soils are relatively nutrient-rich for the region (Newbery et al. 1996, Newbery et al. 1999). Rainfall at the site is fairly equitable over the year, totaling c. 2800 mm on average, but the area is subject to occasional moderate ENSO drought events (Walsh and Newbery 1999, Newbery and Lingenfelder 2004).

The plots were established and first enumerated in 1986 (Newbery et al. 1992). Trees ≥ 10 cm girth at breast height (*gbh*) were measured for *gbh,* identified and mapped. The extent of taxonomic naming to the species level was, and has been since then, very high: vouchers are held at the Sandakan (Sabah) and Leiden (Netherlands) Herbaria. Plots were completely re-enumerated in 1996, 2001 and 2007. In the present paper we analyze data of the two longer periods, 1986 – 1996 (P_1_, 10.00 years) and 1996 – 2007 (P_2_, 11.07 years). For plot structural data refer to (Newbery et al. 1992, 1996, 2011). Measurement techniques and their limitations are detailed in Lingenfelder and Newbery (2009). An over-understory index (*OUI*, continuous scale of 0 – 100), for the 100 most abundant species in the plots, was adopted from Newbery et al. (2011). Three storeys are nominally designated as: overstorey (*OUI* > 55), intermediate (*OUI* 20 – 55) and understory (*OUI* < 20).

### Species selection

Of the 37 tree species which had ≥ 50 small-to-medium sized trees (10 − < 100-cm *gbh*), at 1986 and inside of 20-m borders to the two plots, plus another 11 overstorey ones with 20 or more such individuals — 48 in all (Newbery and Stoll 2013), 38 that had five or more dead trees in P_1_, were selected for the analyses here (Appendix1: Table S1). Among the species excluded was exceptionally *Scorodocarpus borneensis*, with five dead trees, but for which no model for survival could be satisfactorily fitted. Trees in P_2_ were also selected for the same 38 species and size class: numbers dying in this period were also ≥ 5. Table S1 of Appendix 1 has the species’ abbreviations which are used later in the Results. Precise locations (to 0.1-m accuracy) were known for every focal tree and its neighbours, the distance between them being the length of the radius (*r*) of a circle circumscribing the focal tree’s location (Fig. 1a).

**Fig. 1.**
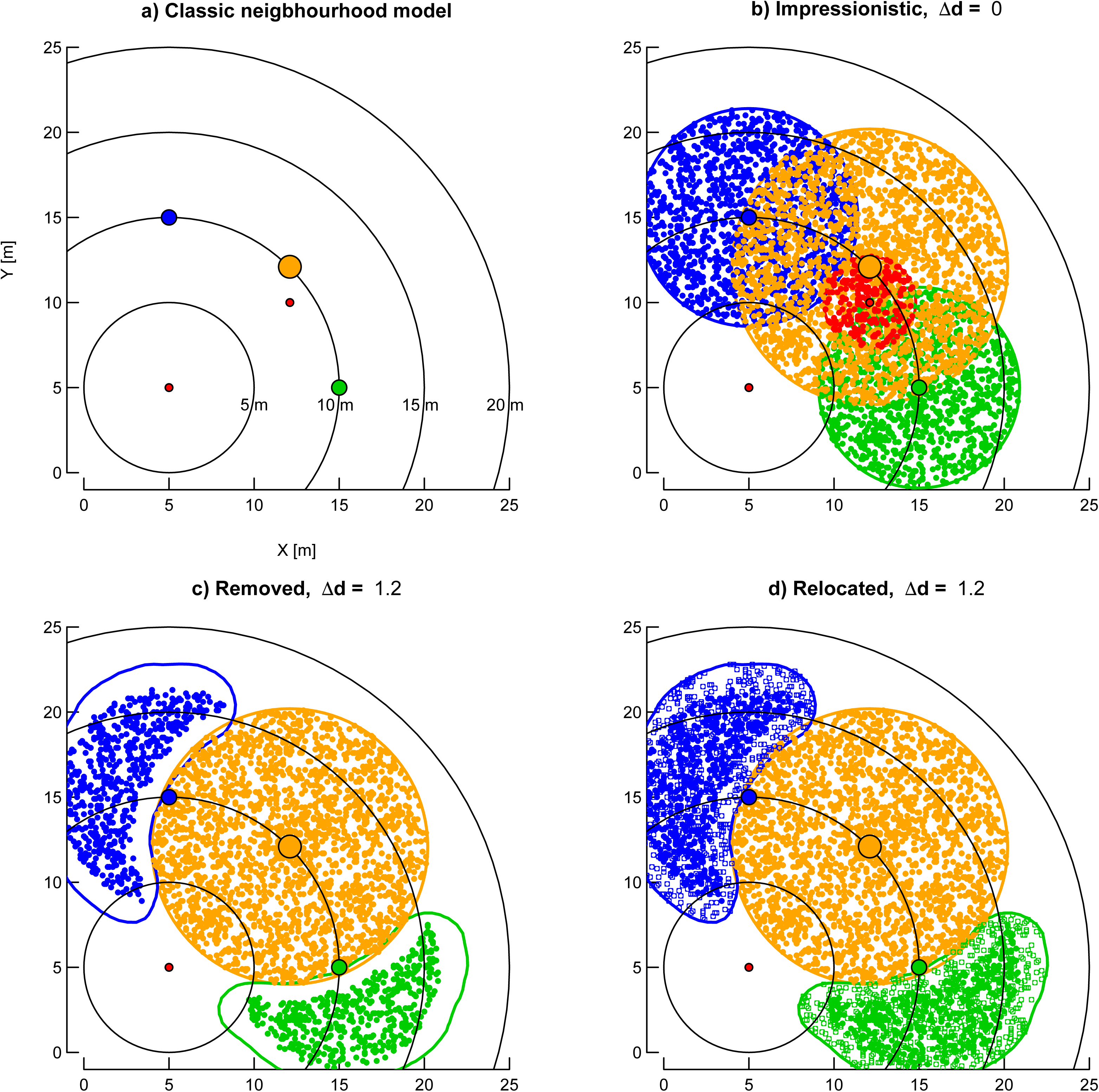
From abstract, mathematical to impressionistic representation of trees with crown plasticity in neighbourhood models. (a) Classical neighbourhood models represent trees as points without any spatial extension. Taking the red tree at X/Y = 5/5 as focal tree, it has no neighbours at 5 m neighbourhood radius (smallest circle centred at 5/5), four within 15 m and, depending on the exact definition of neighbours (i.e. < 10 or ≤ 10 m), one or four neighbours at 10 m neighbourhood radius. (b) Impressionistic representation of tree crowns as circles filled with as many points as the trees basal area at breast height (*ba,* in cm^2^) and crown radii (*cr*, in m) allometrically related to girth at breast height (*gbh*, in cm). Girth of the smallest trees (red at 5/5 and 12.1/10) is 50 cm, those of its neighbours in increasing girth order 175 cm (green at 15/5), 200 cm (blue at 5/15) and 300 cm (orange at 12.1/12.1). These girths correspond to crown radii of 2.7, 5.9, 6.4 and 8.1 m respectively (all-species regression, Table 1). All three bigger neighbours of the focal tree at 5/5 have at least parts of their crowns already within the 5 m neighbourhood of the smallest one. (c and d) Focal tree points may have points of bigger neighbours within their immediate neighbourhood as a function of some distance (Δd). Δd = 0 would allow complete overlap (as shown in b), whereas larger values of Δd flag individual points (open symbols) as being ‘shaded’ if they have points of bigger neighbours within Δd. Two possibilities are used to handle these shaded points. First, (in c) the points are completely removed (*pruned*). Second, (in d), in an attempt to mimic plasticity, shaded points are relocated to unshaded parts of the crown using two-dimensional contour functions (see text) to find the outline of these points. The number of points remains proportional to each tree’s *ba*. The tree at 5/5 is shown here in its role as a focal tree and the other red tree at 12.1/10 (in b) as a conspecific neighbour. This second red tree has no flagged points because Δd = 0. When this second one is taken as a focal tree the first one would have its crown extended, with possibly some points removed or relocated. The relevant focal tree’s position is taken as the stem coordinates (larger coloured points) or, alternatively, as the centroid of the unshaded part of its crown (not shown). For the largest tree (orange) these two positions coincide.

**Table 1.** Regressions between crown radii (*cr*, in m) and stem girth at breast height (*gbh*, in cm; square-root transformed) for 17 tree species at Danum, and the same for eight of these species each with *n* > 35 trees. The data are from of Sterck *et al*. (2001).

| Number of species | *n* | adj. $R^2$ | Term | Estimate<sup>a</sup> | SE | *t* | *P(t)* |
| --- | --- | --- | --- | --- | --- | --- | --- |
| 17 | 443 | 81.4 | intercept | -1.003 | 0.075 | -13.5 | < 0.001 |
|  |  |  | sqrt(*gbh*) | 0.523 | 0.012 | 44.0 | < 0.001 |
| 8 | 382 | 78.5 | intercept | -0.959 | 0.085 | -11.2 | < 0.001 |
|  |  |  | sqrt(*gbh*) | 0.516 | 0.014 | 36.3 | < 0.001 |

<sup>a</sup> major axis (model II) regressions had intercepts and slopes for the two equations as $-1.157/0.549$ and $-1.137/0.547$ respectively.

### Spatial extension of neighbourhood models

Crown radii, *cr* (in m), and their corresponding girths at breast height, *gbh* (in cm), were available for 17 species of the Danum plots (F. J. Sterck, *pers. comm*.). An allometric relationship was fitted with a linear regression by pooling all of these species’ trees (Table 1). For the most abundant eight species, with *n* > 35 individuals each (Sterck et al. 2001), regression estimates were very similar. These more abundant species were: *Aporusa falcifera, Baccaurea stipulata, Mallotus penangensis, M. wrayi, Parashorea melaanonan, Shorea fallax, S. johorensis, and S. parvifolia*, and all occurred in neighbourhood analyses reported in this paper. Relaxing the condition of independence for the X-axis (*gbh*), major-axis regression gave slopes very slightly larger than those for standard linear regression (Table 1). Applying the upper regression equation in Table 1 to all trees ≥ 10 cm *gbh* in 1986 and in 1996, the predicted total canopy covers were 18.516 and 19.069 ha respectively (both plots together), which for a land-surface area of 8 ha, represents leaf area indices (or fold-overlaps) of 2.315 and 2.384. Even trees ≥ 50 cm *gbh* gave corresponding covers of 9.623 and 10.296 ha. Since *cr* ∝ *gbh*^1/2^ and *ba* = *gbh*^2^/4π, *ca* ∝ *gbh or ca* ∝ *ba*^1/2^.

Once the *cr* of each individual tree in the plots had been determined as a function of its *gbh*, the (assumed) circular crown was ‘filled’ at random positions with 10 points per m^2^ crown area (*ppsqmca*: referred to later as ‘equal’) or, alternatively, as many randomly positioned points as the tree’s basal area, *ba* (in cm^2^). The alternative approach led to larger trees having more points per m^2^ crown area compared to smaller trees (*ppsqmca*: larger > smaller referred to later as ‘larsm’, Table 2). The size of each tree in terms of its *ba* was therefore reflected in the number of points per crown (Fig. 1b). Moreover, the alternative approach ensured that the total number of points per plot was identical to the total basal area, Σ *ba*, per plot (inside of borders), when all species were included, and if no points were removed (see below). Filling was realised by multiplying up the original data file with as many rows per individual tree as points within the crown and initially flagging each point as ‘uncovered’. Setting *cr* = 0 for every tree allowed a check of whether the algorithm was working correctly: such a parametrization corresponds to the non-spatial case, i.e. it must give exactly the same results as the non-spatial approach treating individual trees as mathematical points without spatial extension. How the number of points in crowns changes with increasing *gbh* under the ‘equal’ and ‘larsm’ approaches is illustrated in Table 2.

**Table 2.** Illustration of the allocation of points to canopy area (*ppca*) and per square meter of canopy area (*ppsqmca*) across a range of tree sizes (*gbh* – girth at breast height, *cr* – canopy radius, *ca* – canopy area, *ba* – stem basal area) under the two ‘filling’ options: ‘equal’, with a constant point density per m^2^, and ‘larsm’ where larger crowned taller trees have a disproportionally higher density than smaller lower ones.

| *gbh*<br>(cm) | *gbh*<sup>1/2</sup> | *cr* (m)<sup>a</sup> | *ca* (m<sup>2</sup>) | *ba*<br>(cm<sup>2</sup>) | *ppca*<br>equal<sup>b</sup> | *larsm*<sup>c</sup> | *ppsqmca* |  | ratio |
| --- | --- | --- | --- | --- | --- | --- | --- | --- | --- |
|  |  |  |  |  |  |  | equal | *larsm* |  |
| 10 | 3.163 | 0.645 | 1.307 | 7.958 | 6.5 | 7.96 | 10 | 12 | 1 |
| 30 | 5.477 | 1.848 | 10.73 | 71.62 | 18.5 | 71.6 | 10 | 39 | 4 |
| 100 | 10.000 | 4.200 | 55.4 | 795.8 | 42.0 | 796. | 10 | 190 | 19 |
| 300 | 17.321 | 8.007 | 201. | 7162. | 80.1 | 7162. | 10 | 895 | 90 |

<sup>a</sup>: From the equation in Table 1 of main paper: $cr = -1.0 + 0.52 \text{ } gbh^{1/2}$ .
<sup>b</sup>: Based on 10 points m<sup>-2</sup> of canopy area (3<sup>rd</sup> column from right).
<sup>c</sup>: Calculated directly as ‘*ba*’-points per canopy.

Different degrees of overlap were realized by visiting each point within every tree’s crown and evaluating the point’s local neighbourhood. If a point in a tree’s crown lay within a distance, Δd, of a point of a larger (overlapping) tree’s crown, the former was defined as being ‘shaded’ and was flagged. The distances, Δd in steps of 0.2 m, varied from 0.0 (points perfectly overlapping) to 1.2 m (no point of a larger tree’s zone of influence within 1.2 m of a smaller trees). Different values for Δd were allowed that corresponded to conspecific (CON) and heterospecific (HET) neighbours in the spatially extended models which involved two terms, i.e. Δd_CON_ applied to points of different trees of the same species, and Δd_HET_ to points for different trees of different species. This meant 49 different combinations of the Δd-levels on evaluating crown overlap at the start.

Flagged points were then either completely “removed” or they were “relocated” (Figs 1c, d). In the latter case, they were moved to lie within the unshaded part of the crown given by the contour of those points without points of bigger trees crown within Δd; contour function `kde2d` in R package MASS; (Venables and Ripley 2010, R_Core_Team 2017-2019). This procedure attempted to mimic crown plasticity, i.e. the tendency of a shaded crown to grow towards higher light availability and more away from being directly under larger shading neighbours. The contours were allowed to be larger than the original crowns by taking the lowest density contour lines as their outer edges. When all points of a smaller tree were flagged then removal and relocation would result in that tree disappearing from the neighbourhood.

Spatial extension models provide a test of the hypothesis that asymmetric competition for light, i.e. above ground, is the main determining process in tree-tree interactions at the population and community levels at Danum. If spatial models for growth response to neighbours, especially those that accentuate asymmetry (or non-linearity), result in stronger relationships with both species plot abundance and with survival response to neighbours than does the non-spatial one, this would confirm to light being the important factor; if not, the inference would be that nutrients below-ground using a symmetric competition mode are more important. The R-code for the calculations of points allocation to crowns, zone-of-influence overlap, and removal and relocation of points, is available on the GitHub Code Repository Platform (www.github.com) site indicated in Appendix 2, together with some technical details and explanation, and a small test data set (Stoll 2020).

The readjustment of crowns was performed once, across all trees ≥ 10 cm *gbh* in the two plots, for each of the Δd_CON_-x-Δd_HET_-level combinations (same seed for each randomization run). It was therefore set for all focal tree neighbourhood calculations to follow. When three (or more) crowns were overlapping in a common zone, the largest say A (i.e. the dominant) was considered with the first next largest B (below it) and an adjustment made to B. Then, the second next largest C was considered under the crowns of A and adjusted B. The procedure was therefore hierarchical, in the sense of [A −> B] −> C, and it left no overlap between the two adjusted crowns B and C. Non-sequential and other procedures would have been possible but they were not explored.

When a small tree was taken as a focal one (i.e. when its neighbourhood was evaluated) it was represented without any crown extension: only its stem coordinates were needed (Fig. 1). However, when that same tree was a neighbour to another focal one (of either the same or a different species) it would resume its canopy shape and points distribution, in the way they were set at the start by the universal overlap calculations.

When Δd was 0.0, there was no removal or relocation. This was because the probability of a larger crown’s overlapping point coinciding exactly in location with one of a smaller crown below was effectively null (within the limits of real number storage accuracy on the computer). The points might be viewed as being ‘symmetrical’: the tree is therefore ‘fully present’ in terms of its crown dimensions under Δd = 0.0 (Fig. 1b). As Δd increased, though, a rarely occurring distance of 0.2 m could happen by chance, more often so when point densities within the crowns increased (Table 2). This introduced a slight asymmetry. Points were allocated across the circular crowns, just once at random, and each time with the same seed set. (That stage might have been repeated but it would have led to an inordinate increase in computing time, even when say 100 realizations were averaged.) As Δd increased from 0.4 to 1.2 m, more and more flagged points were accumulated when crowns overlapped: the larger the Δd-value, the more ‘asymmetrical’ was the influence of the larger on the smaller crown because this resulted in more removals or more relocations, and hence points becoming sparser in the shaded crown parts under removal, or becoming denser in the unshaded crown parts under relocation. The total number of points is reduced in Fig. 1c, but remains unaltered in Fig. 1d: in both cases the smaller neighbours’ crowns became irregular in shape. If all points of a tree with a small crown became flagged that tree would disappear as a neighbour because its points were either completely removed or had no unflagged crown parts to which they could be relocated.

Once the points of all trees’ crowns had been either removed or repositioned, the neighbourhood of each focal tree was evaluated by summing the number of points within a focal tree’s neighbourhood, a circle with radius *r*, of all larger neighbours (Σ*ba*_ALL_), conspecific bigger neighbours (Σ*ba*_CON_) or heterospecific bigger (Σ*ba*_HET_), each point weighed by a linear distance decay factor (i.e. *ba* · 1/*r*). Analyses without distance decay (Stoll and Newbery 2005, Newbery and Stoll 2013) showed very similar results and these are not reported here. Summations were evaluated in 1-m steps for all neighborhood radii (*r*) between 1 and 20 m for focal trees, within the 20-m borders. To deal with *ln*-transformation of zero values, 1 cm^2^ was added to each Σ*ba* neighbourhood value.

The approach described so far offered, in addition, the possibility of a new way of defining a focal tree’s location. Besides, the original field-recorded stem co-ordinates, the centroid of the unshaded part of the tree’s crown (i.e. mean x- and y-values of focal tree points not having any points of bigger neighbours’ crowns within Δd) could be taken as an alternative, perhaps more relevant, location of that focal tree with respect to maximum light availability. In addition to the size (*ba*-only) models, and models with either one (Σ*ba*_ALL_) or two neighbour terms (Σ*ba_CON_* and Σ*ba_HET_*), crown area considerations provided eight combinations from the three spatial extension factors: ‘equal’ or ‘larsm’ for numbers of *ppsqmca*, times ‘removed’ or ‘relocated’ for point adjustment, times ‘stem’ or ‘crown’ location. The adjustment levels will usually be abbreviated hereon to ‘remov’ and ‘reloc’.

### Model fitting

Models for all possible combinations of radii, for CON and HET neighbors, and Δd (see above) were evaluated (i.e., 7 Δd_CON_ * 7 Δd_HET_ * 20 r_CON_ * 20 r_HET_ = 19600 cases). Least-squares fits for growth, and general linear models with binomial errors for survival, as dependent variables were then applied for all combinations of radii and Δd, excepting a few cases where fitting was not possible. The approach follows that of Stoll and Newbery (2005) and Newbery and Stoll (2013).

The absolute growth rate, *agr*, of focal trees between two times, *t*_1_ and *t*_2_ was modeled statistically as a function of size at the start of the period (*ba_t_*_1_), and one or two neighbour terms which were sums of *ba* of trees that survived the period and were larger than the focal one at *t*_1_, as either all (ALL), conspecific (CON) or heterospecific (HET) neighbours weighted by a linear distance decay. Regressing *agr* upon *ba* for each species per period, trees that had residuals < –3·SD were iteratively excluded. All variables were ln-transformed to normalize their errors. The neighbourhood models were:

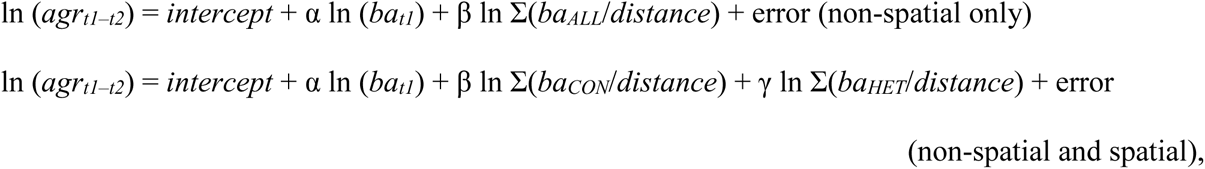

with *intercept*, α, β and γ as the regression parameters to be estimated by the least-squares approach and normally distributed errors. The summations *ba_CON_* and *ba_HET_* were evaluated in 1-m steps for all neighborhood radii between 1 and 20 m, with a border of 20 m. This second model was identical to the C_2_ one of Stoll and Newbery (2005). If less than five focal trees in the sample had CON neighbours, or less than five focal trees had not a single HET neighbour, these model fits were flagged and excluded from further consideration. Their estimates were usually based on respectively either very small or very large radii. The magnitude of effects on growth were quantified by calculating effect sizes as squared multiple partial correlation coefficients, or *t*^2^ / (*t*^2^ + df_resid_) (Cohen 1988, Rosenthal 1994, Nakagawa and Cuthill 2007). All models were fitted using alternatively no, linear and squared distance decay (Stoll et al. 2015).

For survival as dependent variable, the binary variable survival (0/1) was analysed using generalized linear model with binomial errors (logistic regression), and the same model structures as used for growth. When the proportion of dead trees is small (typically, < 0.1) the logit transformation becomes less effective (Collett 1991), and fitting is unreliable or even fails. For this reason, 10 species (see *Species selection* section) were not fully analysable for both survival and growth as dependent variables. No restrictions regarding numbers of CON and HET neighbours were put in place for the survival models. Some estimates (*est*) and their associated standard errors (se) were unrealistically very large, and to avoid these cases, estimates with se > 100 were excluded from the calculations of effects of neighbours on focal tree survival.

Survival effects were estimated by the raw regression coefficients, β, from logistic regression The fitted GLM is of the form ln(odds) = α + βX. Beta therefore expresses the difference in ln (odds) when X increases by 1 unit: exp(β) is the change in odds, or odds-ratio, and (exp(β) – 1)) × 100 is the corresponding increase or decrease in those odds (Fleiss 1994, Agresti 2007, Zuur et al. 2007, Fox 2008, Hosmer et al. 2013).

### Model comparisons

Models were tested and compared by taking a combined pluralistic statistical approach (Stephens et al. 2005, 2007). On the one hand, the classical frequentist approach is needed to assess the strength of model fitting and allow a hypothesis-testing framework (recently defended by Murtaugh 2014 and Spanos 2014), whilst on the other hand, the information-theoretic approach provides an efficient means of model comparison and inter-model summarization (e.g., Burnham et al. 2011, Richards et al. 2011), with the final outcome being purely relative yet avoiding ‘data-dredging’ and undue heightening of confidence through multiple testing that contravenes the rules of independence (Symonds and Moussalli 2011). The formulation of good alternatives to the null hypothesis may lead to more informed model fitting and testing than when none are posed beforehand (Burnham and Anderson 2001, Anderson and Burnham 2002). Several cautionary points have been raised in the literature concerning the general use of the information-theoretic approach (see Richards 2005, Arnold 2010). Especially, it does not sit well on its own within a critical rationalist approach to science. It provides for a valuable heuristic complement, however.

Accordingly, the analysis here was a mixture of approaches, structured as follows. First, the central reference model is just tree size (*gbh*), plus the basal area (*ba*) of ‘ALL’ neighbours’ basal area within radius *r*. The question was whether model fits were improved by having CON and HET terms in place of ‘ALL’, and then having fixed spatial terms for them (eight alternatives). The differentiation between CON and HET constitutes one quantum-level change in information and the addition of spatial form a second. These eight spatial forms were not fully independent of one another in their information because CON and HET *ba*-values will be highly correlated. The modes of decay (the inverse-distance weighting applied to neighbourhood BA) offered three different ways to improving model fitting. Accordingly, the reference model, *ex hypothesis,* for within periods 1 and 2 and for growth and for survival, was the non-spatial ‘ba + ALL’ one with linear decay. It may not have necessarily been the best fitting model compared with the other non-spatial and spatial ones. Individual species’ best-fitting models were said to differ strongly from the reference model when the ΔAICc was > |7|, and to be not different when ΔAICc was ≤ 7 (Burnham and Anderson 2010). A ΔAICc-value of 7 or one more negative meant a ‘much better’ model, one of 7 or more positive, a ‘much worse’ one. ΔAICc > |7| is equivalent to a Pearson-Neyman significance level of *P* ≤ 0.003 − 0.005 (with k = 1 to 4 independent variables; Murtaugh 2014).

The dependence of CON-effects for growth or survival at the community level (one point for each of the 38 species) on total plot *BA* per species, was estimated again by standard linear regression. Because models with ΔAIC_c_ in the 2-7 range have some support and should perhaps not be too readily dismissed (Burnham et al. 2011, Moll et al. 2016), results and estimates from all different non-spatial and spatially extended models are reported. Regression statistics from specific models but different neighbourhood radii or Δd were often very similar and had very small ΔAIC_c_ among them. The correlations between differences in the CON or HET effect sizes on growth between periods (P_2_ – P_1_) and CON or HET effect (expressed as raw coefficients) on survival in P_1_ or P_2_ were tested at the community level with the expectations stated in the Introduction.

The final effect sizes, for a non-spatial or spatial model, per species and period, were found by averaging raw coefficients (equally weighted) across all radii and Δd-values with fits ≤ 2 ΔAICc of the best one, i.e. the one with the smallest AICc (Ripley 2004, Claeskens and Hjort 2008). Averaging was considered valid here because all of the models involved had exactly the same structure (same terms), and so within species and period they would be differing in the exact combination of r_CON_, r_HET_, Δd_CON_ and Δd_HET_ values used (see (Cade 2015, Banner and Higgs 2017), for general discussion). Averaging was unweighted, i.e. no Akaike weights, w_i_, were applied since there was no a priori reason to do so within such a small AICc-band (Burnham and Anderson 2001, 2010). There were often very many models in this 2-ΔAICc range, and in some cases there was a change in sign for a minority of them; r and Δd values were often very close to one another. Alternative ways of summarizing these coefficients, namely averaging only those values with sign the same as that of the overall mean, or taking the medians, resulted in very small differences in the overall outcomes, and hence the simple arithmetic mean was used.

Calculations were performed largely in R (version 3.4.2; R_Core_Team 2017-2019), using package `AICcmodavg` (Mazerolle 2019) to find AICc, the small-sample-size correction of AIC, Akaike’s Information Criterion (Hurvich and Tsai 1989, Burnham and Anderson 2010). Predicted *R*^2^-values were found using the predicted residual error sum of squares (PRESS) statistic (Allen 1974, Fox and Weisberg 2011). The calculation of (pseudo-) *R*^2^ for logistic regression followed (Mittlböck and Schemper 1996), where *R_L_* ^2^ = [(L_0_−L_p_)/L_0_] · 100, L_0_ and L_p_ being the log-likelihoods of the model with only the intercept and with the nearest-neighbour (spatial) terms respectively (Hosmer et al. 2013, p. 184). The *R* ^2^-values were adjusted as in linear least-squares regression, although they are not directly comparable.

### Randomizations

To more rigorously test the significance of the CON and HET coefficients, the model fitting was re-run for *n’* =100 randomizations of locations of trees within the plots. The randomization outcomes of Newbery and Stoll (2013) were re-used. The method that produced them is described in detail in Appendix B (*ibid*.): it involved simple rules allowing different minimum distances between nearest-neighbour trees within the same and different size classes (six defined), and it ensured that the same overall frequencies of size distribution for each species were maintained. On each run focal trees were those, of each species (in the size class used for the observed trees), which were now located within the 20-m plot boundaries: CON and HET neighbourhoods were accordingly realistically randomized; any spatial clustering in observed tree distributions will have been removed as well. The procedure also tests whether the relationships in the community-level graphs might have arisen by chance.

## RESULTS

### Finding the best fit models

Frequency distributions of growth and survival CON (raw) coefficients within 2ΔAICc, for each of the 48 species first analysed in P_1_ and P_2_ (survival in P_1_ gave 46 histograms) using the non-spatial and the spatial “larsm/reloc/crown” and “larsm/remov/crown” modes, were inspected visually for evidence of obvious bimodality, or multimodality, which would indicate inconsistency in the final averages estimated (Appendix 3). Bimodality was judged to be present when there were two clear modes separated by being to either side of zero or otherwise by a peak difference at least approximately twice the mean coefficient. Of the 570 cases, 16 (2.8%) showed evidence of bimodality (seven ‘±-zero’, nine ‘≥ 2-fold difference’). There were no cases of multimodality. Repeating this analysis with growth and survival HET coefficients, but for just the spatial “larsm/reloc/crown” mode, just five of 190 cases (2.6%) were correspondingly bimodal (two ‘±- zero’, three ‘≥ 2-fold difference’). For both CON and HET coefficients bimodal cases were occurring across many different species and not the same for different modes or periods. Different peaks were arising because models were fitting at two clusters of similar radii (and delta-values) suggesting that occasionally two neighbourhood relationships may have been operating. Overall, these cases are too infrequent to have affected the main results to any major degree.

A particularly interesting feature is that for non-spatial models many species had over 100 (out of the maximum of 400 possible), and for spatial models thousands or tens of thousands (out of 19’600 maximally), CON-effect estimates within the 2ΔAICc-band. This latter maximum was actually reached for *Pentace laxiflora* (‘larsm/reloc’ and ‘larsm/remov’) growth in P_2_, and for *Dehassia gigantifolia* (‘larsm/reloc’) survival in P_1_. It means that many models were indistinguishable in their estimates of CON-effects, and that radius or Δd interacting with *ba* had little role, and presumably the main information lay in the presence or absence of any neighbour within 20 m of the focal tree.

### Individual species’ model fits

With linear distance decay and any of the eight spatial model forms, having ‘ba + CON + HET’ as the terms for growth responses in P_1_ led to 8-14 out 38 species (on average 29%) showing better fits than when using just ‘ba + ALL’ terms. No particular model excelled by being better fitting for appreciably more species, although for four of them (11%) the fits were better by just using ‘CON + HET’ without spatial extension (Appendix 1: Table S2a). For P_2_ the outcome was similar but slightly weaker in that 7-14 species (28%) had correspondingly better fits. No-decay models were similarly frequent to linear distance ones, although squared distance models were fewer in both periods. The number of species which had worse fits when including spatial extension, compared with ‘ba + ALL’, were very few in P_1_ (0-2), and slightly more for P_2_ (1-3).

Spatial models for survival responses led to very few species with improved fits in P_1_ and P_2,_ for linear distance decay 2-5 (8%) and 2-7 (12%) out of 38 species respectively (Appendix 1: Table S2b). Replacing the model terms ba + ALL by ba + CON + HET, resulted in 0-1 species with improvements in P_1_ and P_2_: the corresponding number of species with worsening fits when comparing spatial models with ‘ba + CON + HET’ was 0-1 in both P_1_ and P_2_ (Appendix 1: Table S2b). Tables of non-spatial and the two spatial, ‘larsm/reloc/crown’ and ‘larsm/remov/crown’, model parameter fits for all species, for growth and survival, in P_1_ and P_2_ are given in Appendix 4.

Resetting the reference model to ‘ba + CON + HET’ instead of ‘ba + ALL’, for spatial linear decay modes, the number of species with improved fits decreased to 3-10 (17%) for P_1_ and 4-11 (20%) for P_2_, especially ‘larsm’ models the reduction was down to 3-4 for P_1_ and 4-6 for P_2_ better fitting (Appendix 1: Table S3) whilst just 0-1 and 1-3 species with ‘larsm’ models were respectively worse than the reference one. These comparisons imply that part of the improved model fitting under spatial extension compared with ‘ba + ALL’ was because CON and HET were being used as separate terms.

Considering the individual species’ fits in P_1_ and P_2_, over the non-spatial and two spatial ‘larsm/reloc/crown’ and ‘larsm/remov/crown’ models, for growth, 9-13 (29%) of species had adjusted *R*^2^-values ≥ 50% and only 8-12 (26%) < 20% (Appendix 1: Table S3); and for survival — recalling here that *R*^2^ is a ‘pseudo’-estimate — far fewer at 0-2 (3%) had *R*^2^-values ≥ 50% and as many as 33-35 (89%) of species with just < 20%. *P*-values of CON coefficients were ≤ 0.05 for 9-22 (41%) for growth and 6-11 (22%) for survival; 7-16 (30%) and 13-21 (45%) with *P* ≥ 0.25. HET coefficients showed similar distributions, for growth 15-22 (49%) at *P* ≤ 0.05 and for survival 4-8 (16%) of species, with correspondingly 8-13 (28%) and 14-21 (46%) at *P* ≥ 0.25 (Appendix 1: Table S4). In general, the significance of fits and their coefficients were weaker for survival than growth regressions, but the CON and HET coefficients’ *P*-values were rather similar. Adjusted *R*^2^- values were similar for the non-spatial and two spatial models of more interest, though CON coefficients were more often significant (*P* ≤ 0.05) for spatial than non-spatial, and HET coefficients showed a slight trend in the opposite direction (Appendix 1: Table S4).

### Effect sizes dependence on species’ plot basal area abundance and density

Regressing the 38 species’ CON effect sizes on growth in P_1_, whether at the individual species’ level they were significant or not, against plot BA (log_10_-transformed), showed that the non-spatial model with ‘ba + CON + HET’ led to a substantially better fit (*P* ≤ 0.001) than with ‘ba + ALL’ (*P* = 0.18) (Table 3a), and accounted for the maximum adjusted and predicted *R*^2^ of all non-spatial and spatial models. However, including spatial extensions with the eight different forms led to reduced fits, rather surprisingly, with adjusted and predicted *R*^2^ decreasing by about a third (and *P* ≤ 0.01). The eight spatial forms differed little from one another in fit although ‘equal’ was slightly better than ‘larsm’. In P_2_, the relationships were similar but less strong and less significant, the non-spatial ba + CON + HET model achieving significance only at *P* ≤ 0.05. Predicted *R*^2^- values were very low, much lower than the adjusted values for the eight spatial forms (Table 3a). In P_1_ and in P_2_ the slopes of the relationships changed little between non-spatial and spatial modes, and if at all were slightly less negative for the spatial ones (Table 3a). To recall, the stronger the CON or HET effect the more negative it was, so if the species’ values decreased with increasing plot BA the expected slope of the relationship would be negative.

**Table 3.**
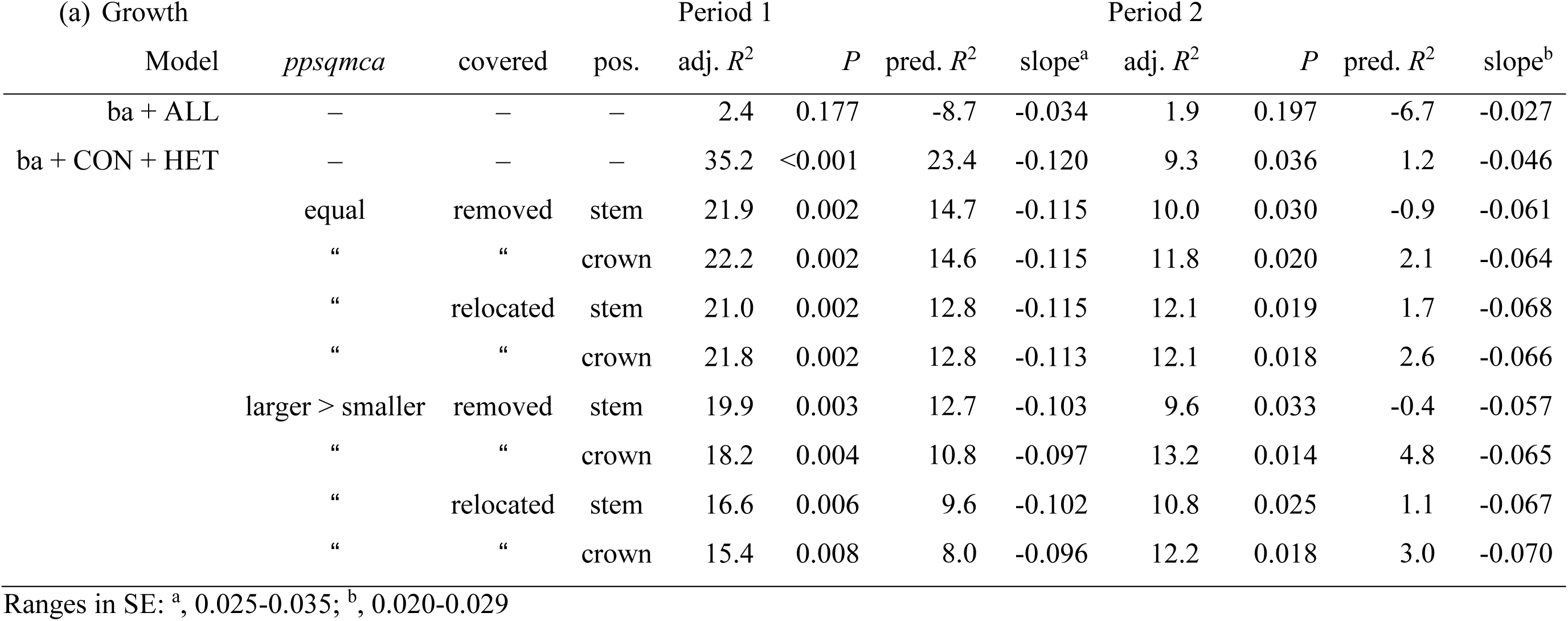
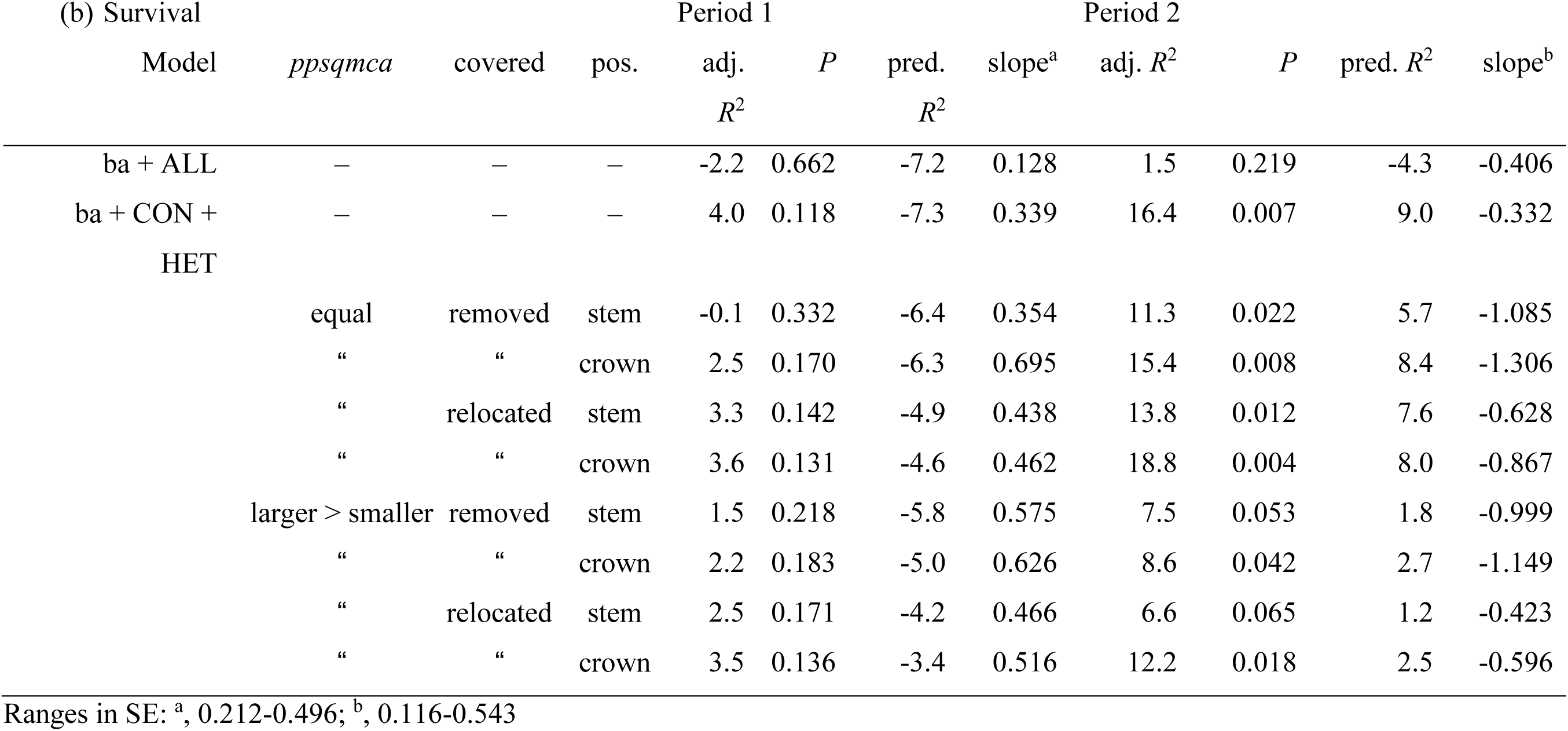
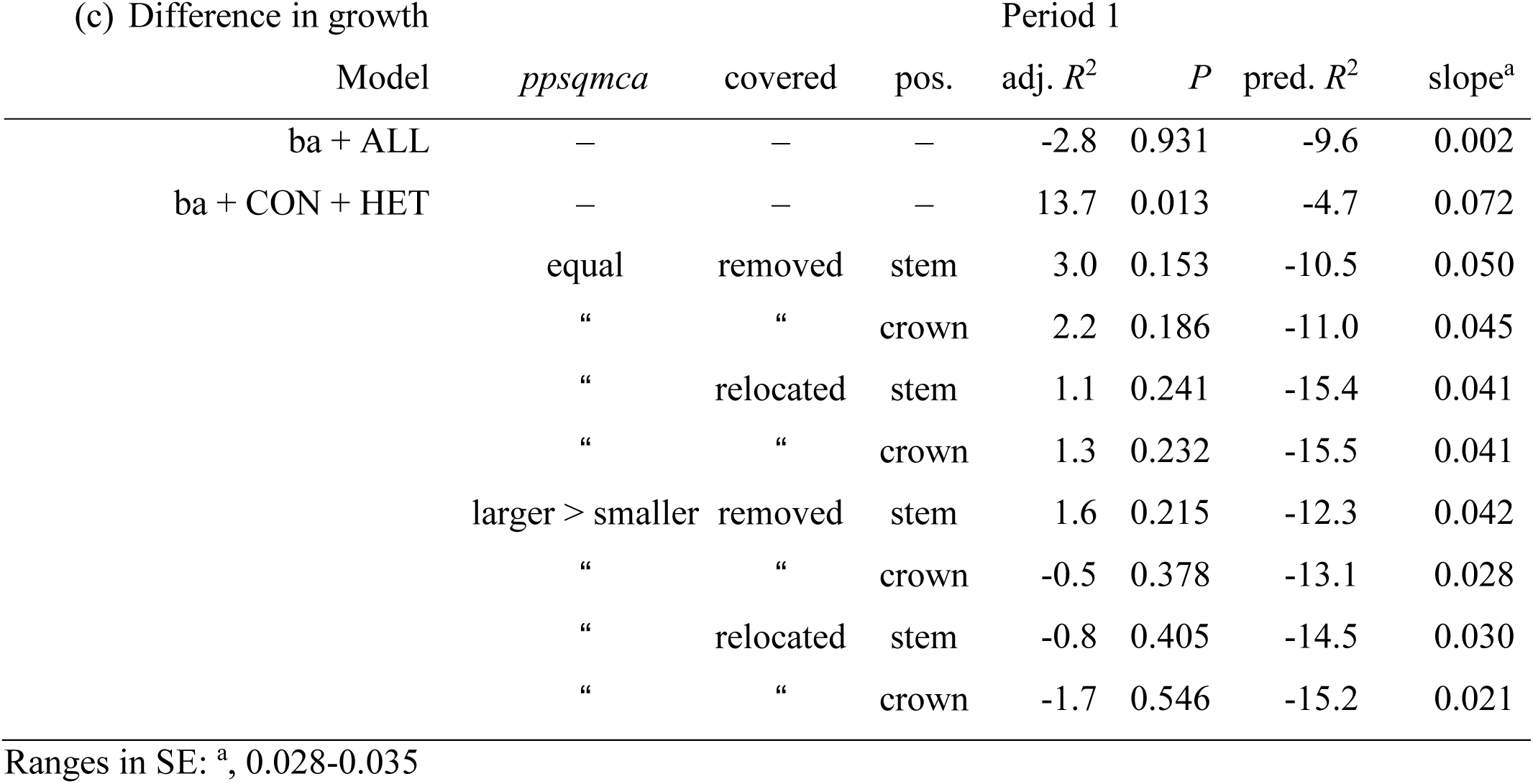
Dependence of conspecific (CON) effects in terms of (a) absolute growth rate, and (b) survival in periods 1 and 2 (P_1_, P_2_), and (c) difference in CON effect sizes in growth rates between periods (P_2_ − P_1_), of the 38 species, for the different non-spatial and spatial models using linear distance decay, on plot-level basal area (as log_10_[*BA*]). The models involved basal area (*ba*) and one or two neighbourhood terms and, in the spatial case, one of the eight different forms of crown extension (as structured in Table 2). The final effects sizes came from model averaging, i.e. finding mean CON coefficients across all model fits ≤ 2 ΔAIC_c_ of the best fitting one.

Considering the 38 species’ CON effects on survival in P_1_, regressions for both non-spatial and spatial forms were very weakly dependent on plot BA (*P* = 0.13 to 0.33 for the spatial ones). However, the relationships here were stronger in P_2_ than P_1_, and showed improved fits for spatial forms (*P* < 0.05 in all but two cases) over non-spatial ones, *R*^2^-values reaching almost as high as those found for growth in P_1_ (Table 3b). The slopes of the relationships were positive in P_1_ and negative in P_2_, becoming steeper for spatial compared with non-spatial modes, despite the lack of significance (Table 3b). Difference in CON effect size on growth P_2_−P_1_ (i.e. effect size and P_2_ minus that at P_1_) regressed on plot BA had much lower adjusted and predicted *R*^2^-values than for CON effects on growth in P_1_ and P_2_ separately, and most notably the non-spatial model with ‘ba + CON + HET ‘was far poorer fitting (Table 3c). None of the eight spatial modes had significant fits (i.e. *P* ≥ 0.15). Slopes for these differences in CON effect size were all positive but less so for spatial than non-spatial modes.

Of the nine spatial and non-spatial model forms times eight ‘CON-HET vs growth-survival vs P_1_-P_2_’ combinations (72 in all), correlations between effect sizes (for growth), or raw coefficients (for survival), and log_e_ (population size), all were weak and insignificant except for HET effect on growth in P_1_ across all eight spatial model forms was consistently positive (*r* = 0.433 to 0.500, *P* ≤ 0.005. Variables were all approximately normally distributed except HET effect on growth in P_2_ with one distinct outlier.

### Cross-correlations between the eight spatial models

For each of the eight CON-HET x growth-survival x P_1_-P_2_ combinations there were, among the 28 pair-wise correlations of the eight different spatial models (38 species selected), several-to-many showing very high agreement (*r* = 0.96 to > 0.99; Appendix 1: Table S6). Differences arose as the correlations between models became weaker. For CON-growth-P_1_ and -P_2_ which spatial model form was used had little influence as the correlations were always very high. For the corresponding HET-growth-P_1_ and -P_2_, the minimum *r*-values (and corresponding *t*-values) decreased moderately, especially for ‘larsm/reloc’ vs ‘equal/remov’. In comparison to growth based variables, correlations between spatial models based on survival variables dropped considerably, especially for ‘larsm/reloc’ vs ‘equal/remov’. Crown or stem location accounted very little for differences between spatial models. Growth models, therefore, depended very little on the spatial form, but survival models so did much more. Spatial extension was apparently influencing survival more than growth across species, for CON and HET cases.

### Radii of neighbourhood effects

For the “larsm/reloc/crown” model, across the 38 species, mean of mean model-fitted radii were similar for growth and survival CON and HET coefficients at P_1_ (9-10 m). By P_2_ CON mean radii exceeded HET ones for both growth and survival (Table 4), CON and HET means being on average closer to 10 m – midway for the radii modelled (viz. ≤ 20 m). The weaker CON effects in P_2_ than P_1_ is commensurate with increasing mean radius. Mean radii were based in some cases on very many model fit estimates where radii ranged greatly, and radii were not normally distributed for every species. Nevertheless, mean range of radius, over which effects were averaged, was clearly smaller for CON than HET, for both growth and survival at P_1_ (10 vs 13-14 m); but by P_2_ these were much more similar at 11 m for growth and 13 m for survival (Table 4). The lessening of CON coefficients from P_1_ to P_2_, and their moving towards the HET ones, occurred as radii became more similar.

**Table 4.** Means of means (± SE) and of ranges of radii within the models fitted within 2ΔAICc, across the 38 species, and the means of the corresponding regression slopes of CON (conspecific) or HET (heterospecific) coefficients on radii, for the growth and survival response variables in periods P_1_ and P_2_, using spatial model ‘larsm/crown/reloc’.

|  |  | Growth |  | Survival |  |
| --- | --- | --- | --- | --- | --- |
|  |  | CON | HET | CON | HET |
| Mean of radius mean (m) |  |  |  |  |  |
| | $P_1$ | $9.85 \pm 0.83$ | $9.55 \pm 0.86$ | $9.99 \pm 0.78$ | $8.99 \pm 0.74$ |
| | $P_2$ | $13.30 \pm 0.72$ | $7.86 \pm 0.91$ | $11.11 \pm 0.85$ | $7.98 \pm 0.72$ |
| Mean of radius range (m) |  |  |  |  |  |
| | $P_1$ | $9.87 \pm 1.15$ | $13.16 \pm 1.17$ | $10.03 \pm 1.11$ | $14.47 \pm 1.04$ |
| | $P_2$ | $10.74 \pm 0.99$ | $10.89 \pm 1.34$ | $13.16 \pm 1.08$ | $13.03 \pm 1.11$ |
| Mean slope of estimate on<br>radius ($\text{m}^{-1} \cdot 10^3$) | | | | | |
| | $P_1$ | $10.31 \pm 4.13$ | $-20.09 \pm 4.79$ | $0.00 \pm 52.7$ | $6.59 \pm 8.61$ |
| | $P_2$ | $6.14 \pm 2.27$ | $-9.04 \pm 3.15$ | $24.8 \pm 12.1$ | $0.30 \pm 16.70$ |

Across the 38 species, again for the same spatial model, and using the raw coefficients (i.e. not effect sizes for growth), mean CON coefficient was positively correlated with mean CON radius at which the effect operated for growth in P_1_ (*r* = 0.545, *P* ≤ 0.001) and P_2_ (*r* = 0.370, *P* ≤ 0.05), and survival in P_1_ (*r* = 0.208, *P* > 0.05) and P_2_ (*r* = 0.436, *P* ≤ 0.01), although mean difference in CON coefficients for growth in P_2_−P_1_ were not significantly correlated with mean CON radii for growth in P_1_ and P_2_ (*r* = −0.137, *P* > 0.05). In contrast, mean HET coefficient was weakly (*P* > 0.05) negatively correlated with mean HET radius for growth in P_1_ (*r* = −0.096) and P_2_ (*r* = −0.248), and survival in P_2_ (*r* = −0.169), although survival in P_2_ showed a correspondingly much stronger positive correlation (*r* = 0.509, *P* ≤ 0.001), and mean difference in HET coefficients for growth in P_2_−P_1_ were also not significantly correlated with mean HET radii for growth in P_1_ and P_2_ (*r* = −0.030, *P* > 0.05). Thus, species with strong negative CON coefficients for growth tended to be operating at short distances, and the strong positive ones at much larger distances (≤ 20 m). Even so, differences in CON growth coefficients P_2_−P_1_ across species were less related to neighbour distance than those for P_1_ and P separately.

Correlations between CON and HET regression coefficients, growth and survival, P_1_ and P_2_, with best fitting radii within the 2ΔAICc range were also found. Histograms of the 38 species correlation coefficients revealed a clear difference between CON and HET: the former were always bimodal, with strong negative and positive correlations, and the latter were normally distributed around zero, i.e. most species were weakly or not correlated with radius (Appendix 1: Fig. S1a). This would reflect the more species-specific nature of CON effects (by definition) versus the highly mixed and diverse ones bundled into HET effects. The difference values will have been very highly spatially autocorrelated as they were using neighbouring *ppsqmca* locations. A change in radius increments across the defined neighbourhood crowns (see Fig. 1), so different combinations of points would be achieved as different neighbour’s crowns are encompassed.

Regressions for all of the 38 species studied, again for growth and survival, P_1_ and P_2_, estimated the changes in CON or HET coefficient per meter of radius (Table 4). For ca. 90% of the species the slope-values are very small. Graphs of coefficient versus radius (Appendix 5) indicated mostly continuous set of lightly curved lines, increasing or decreasing with radius; occasionally there was a mixture of lines resulting in little overall trend, rarely disjunctions (five cases of 1-2 m). Histograms of the slopes for coefficient versus radius did highlight, however, a few strongly positive or negative outliers, especially for CON survival in P_1_ (Appendix 1: Fig. S1b). The five important species’ cases that might have biased the study’s conclusion are highlighted in the Appendix. Strong bimodality explained some of them because when the tail values formed a minority of points, basic assumptions of regression were not being met. Only very small shifts in mean CON effects on survival in P_1_ occurred on the exclusion of tail values.

### Crown overlap and readjustment

Correlations between mean Δd_CON_ and mean Δd_HET_ (across the model fits within 2ΔAICc, as for coefficients and radii) for growth and survival in P_1_ and in P_2_, for ‘larsm/crown/reloc’ were all very weak and insignificant (*r* = −0.077 to 0.034, *P* ≥ 0.65). Likewise, for either Δd_CON_ or Δd_HET_ between growth and survival, in P_1_ and in P_2_, correlations were weak (*r* = −0.240 to 0.057, *P* ≥ 0.15). Using ‘larsm/crown/remov’ as the model gave very similar outcomes.

Mean Δd_CON_ and Δd_HET_-values across the 38 species (with ‘reloc’) were nevertheless very similar and all sitting near the centre of the level range of 0 to 1.2 m preset (Table 5): the overall average was 0.524 (range 0 – 1.2). Means for survival were slightly higher than those for growth. Means using ‘remov’ in the model were also very close to those with ‘reloc’. Histograms of the 38 species’ mean Δd-values, for the different combinations of CON/HET, P_1_ and P_2_ and growth/survival, were all either roughly even or slightly normally distributed, in the full range 0 to 1.2, but none were skewed (Appendix 1: Fig. S2). The expected possible separation then of models fitting best with Δd = 0 versus those with Δd > 0 – recalling the important qualitative difference and its consequence – was not obvious.

**Table 5.** Means of means (± SE) of Δd-values, with ranges in parenthesis, within the models fitted within 2ΔAICc, across the 38 species, for the growth and survival response variables in periods P_1_ and P_2_, and using spatial model ‘larsm/crown/reloc’. CON – conspecific (Δd_C_); HET – heterospecific (Δd_H_).

|  | Growth |  | Survival |  |
| --- | --- | --- | --- | --- |
|  | CON | HET | CON | HET |
| $P_1$ | $0.494 \pm 0.057$<br>(0 – 1.20) | $0.485 \pm 0.044$<br>(0 – 1.01) | $0.573 \pm 0.047$<br>(0 – 1.16) | $0.602 \pm 0.034$<br>(0 – 1.00) |
| $P_2$ | $0.527 \pm 0.044$<br>(0.09 – 1.20) | $0.446 \pm 0.047$<br>(0 – 0.99) | $0.519 \pm 0.045$<br>(0 – 1.20) | $0.549 \pm 0.040$<br>(0 – 0.98) |

Despite these overall community-level means being so similar and central, species differed individually from one another markedly in the distributions of their Δd_CON_ and Δd_HET_-values for the model fits, across the full 0-to-1.2 range. Some had best fits with only Δd_CON_ or Δd_HET_ = 0, or alternatively 1.2, others had low- or high-valued skewed peaks, and several showed clear declines from, or inclines towards, the scale extremes. The ‘boxes’ defined by Δd 0 – 1.2 for CON and HET were largely, and approximately evenly, filled with points of the species means; and hence the very poor correlations noted. Visually matching species’ histograms of Δd_CON_–values for models using growth versus those using survival, both in P_1_ (‘reloc’) – and the same for Δd_HET_ (38 x 2 = 76 combinations), 30 showed a strong tendency for Δd to be low or 0.0 for growth yet high or at 1.2 for survival, and 14 the converse, i.e. 58% of cases were radically opposite in most frequently fitted Δd-values for the two responses. Once more, the models with ‘remov’ barely differed from ‘reloc’, having very similar patterns.

Mean Δd_CON_ and Δd_HET_-values for the 38 species were, furthermore, not strongly or consistently related to the CON and HET effect sizes in the best fitting models either, when again placing attention on growth and survival responses in P_1_ and in P_2_ (the ‘reloc’ model). Seven of eight correlations were insignificant (*P* > 0.05), and the one for CON/survival/P_2_ only marginally so (*r* = −0.331, *P* = 0.043). With ‘remov’ in place of ‘reloc’ in the model, a different period was significantly highlighted, as CON/survival/P_1_ (*r* = −0.396, *P* = 0.014). Hence, CON and HET effect sizes were seemingly unrelated to degree of overlap of crowns (ZOIs). Correlations of species’ mean Δd_CON_ and Δd_HET_-values and plot-level BA were poor too (*r* = −0.160 to 0.097, *P* ≥ 0.34).

### Effect sizes on growth, and effects on survival

The absolute values of the effect sizes on growth, as squares of the partial correlation coefficients, are proportions of the total model variance (*R*^2^) accounted for in each species’ fitting. Squared partial correlations (e.g. CON) are proportions of the variance in Y that is unaccounted for by the other variables (ba + HET) in multiple regression, i.e. when these other variables are set constant (at their means) and have no variance. By contrast semi-partial or part correlations squared would express the proportion of variance in Y accounting for all variables in the model (overall model *R*^2^; Cohen 1988, Warner 2013). For the two periods, CON and HET, and for the two spatial models “larsm/reloc/crown” and “larsm/remov/crown”, half of the species defined by the eight medians had variances of just 2.7 to 4.4% or less, the upper quartiles reaching 7.2 to 13.0%, and just a very few species attaining > 20%. It is these few that give the most leverage to the relationships at the community level. Fits with relocated crowns were slightly better overall than with removed crowns. These variances, in an approximately similar order of magnitude, are realized by the spread of differences in CON effects on growth in Figs 2 and 3 (signs reassigned).

**Fig. 2.**
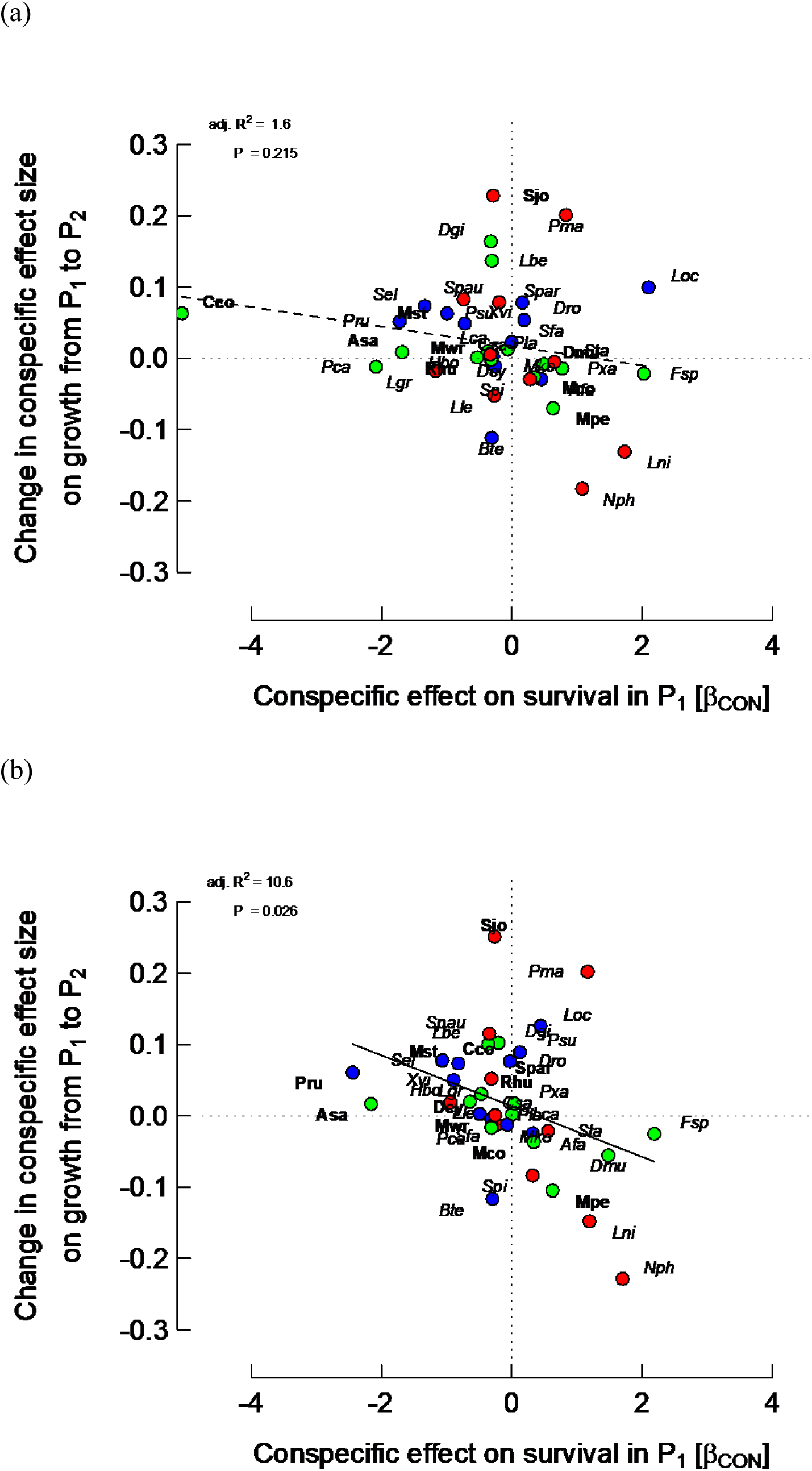
Relationships between *differences* in conspecific (CON) effect sizes on growth rates between periods (P_2_ − P_1_) and CON effects on survival in period 1 (P_1_; β_CON_ = change in log[ODDS] per unit change in log(1 + sum(CON*_ba_*))) for the 38 species, from spatially-extended neighbourhood models with size (*ba*) and two neighbour terms (HET and CON). Points of focal trees having points of bigger neighbours within their zone of influence were either (a) removed, or (b) relocated. Crown position was used as focal tree position in order to evaluate its neighbourhood (lines 8 and 10 in Table 3a refer). Colour codes for points. OUI, over-understorey index: < 20 (green), ≥ 20 – 55 (blue), > 55 (red). Species’ labels: italicized, CON effect on survival in P_1_ with *P* ≥ 0.05, in bold *P* < 0.05. Points (each representing a single species) above or below the horizontal dotted line (Y = 0) indicate respectively decrease or increase of CON effects on growth, going from P_1_ to P_2_. Points to the left or right of the vertical dotted line (X = 0) indicate respectively negative or positive CON effects on survival (CON neighbours increasing or decreasing mortality.

**Fig. 3.**
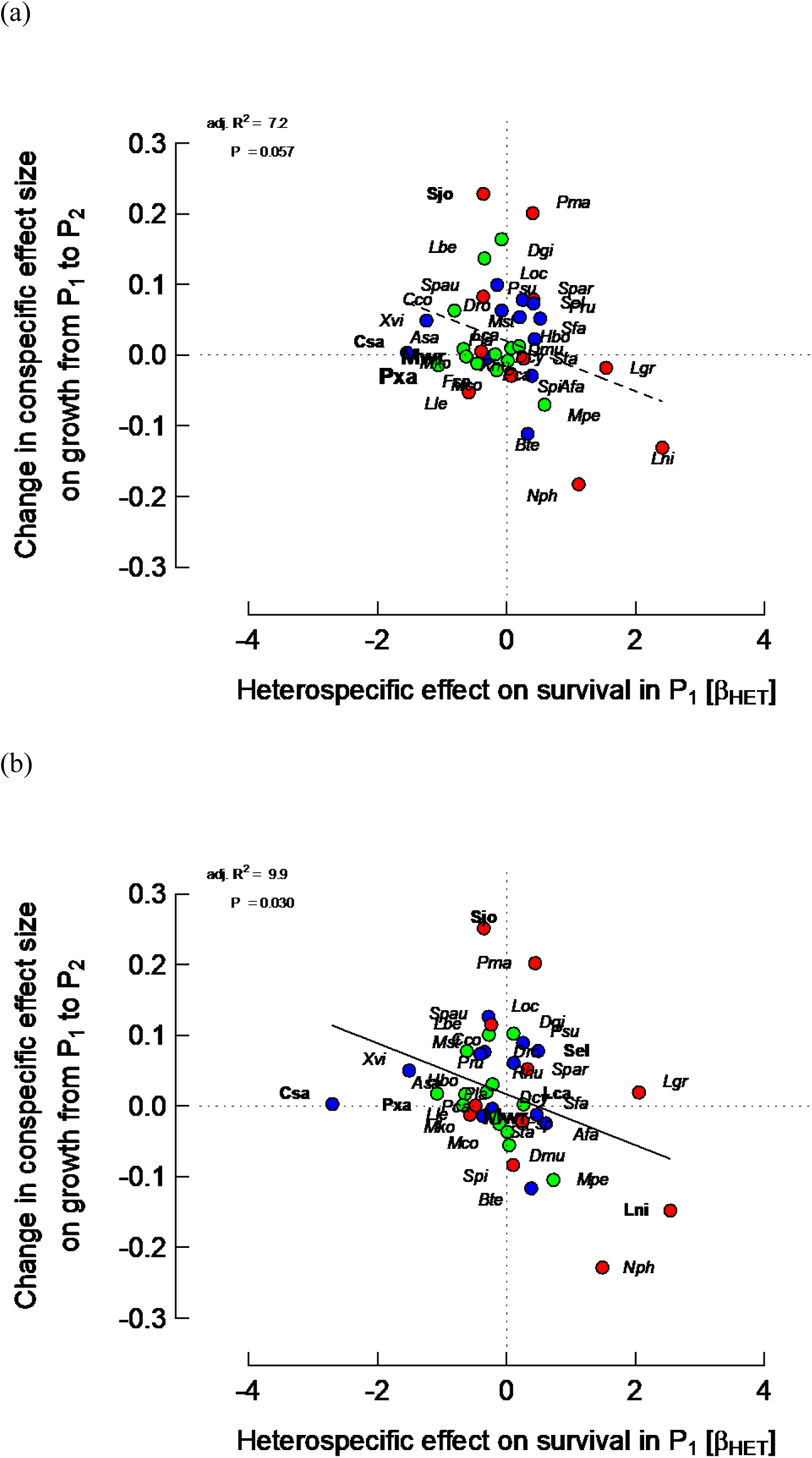
Relationships between *differences* in conspecific (CON) effect sizes on growth rates between periods (P_2_ − P_1_) and HET effects on survival in period 1 (P_1_; β_HET_ = change in log[ODDS] per unit change in log(1 + sum(HET*_ba_*))) for the 38 species, from spatially-extended neighbourhood models with size (*ba*) and two neighbour terms (HET and CON). Points of focal trees having points of bigger neighbours within their zone of influence were either (a) removed, or (b) relocated. Details are the same as for Fig. 2, except for species’ labels: larger font, HET effect on survival in P_1_ with *P* < 0.01.

### Community-level graphs

The strongest correlations were between CON difference effect on growth P_2_−P_1_ and CON or HET effect on survival in P_1_ for “larsm/reloc/crown” with *r* = −0.361 and −0.352 respectively (*P* = 0.026 and *P* = 0.030; Figs. 2b, 3b). Those corresponding for “larsm/remov/crown” were weaker, with *r* = −0.206 and −0.311 (*P* = 0.215 and 0.057; Figs. 2a, 3a). CON difference effect on growth on the sum of CON and HET effects on survival in P_1_, however, showed an even stronger correlation for “larsm/reloc/crown” with *r* = −0.437 (*P* = 0.006), although rather less strongly for “larsm/remov/crown” with *r* = −0.296 (*P* = 0.071). Correlations between CON difference effect on growth P_2_−P_1_ and CON, or HET, effect on survival in P_2_, however, were very poor with a range in *r* = −0.023 to 0.055 (*P* = 0.74 to 0.89); and likewise the sum of CON and HET effects versus survival in P_2_, for “larsm/reloc/crown” and “larsm/remov/crown” were very weak (*r* = −0.014 and 0.034; *P* = 0.93 and 0.84).

By contrast to the difference in CON growth effects, the HET difference effect on growth P_2_−P_1_ and CON, or HET, effect on survival in P_1_ for “larsm/reloc/crown” or “larsm/remov/crown” were all insignificantly correlated with the range in *r* = −0.136 to −0.003 (*P* = 0.42 to 0.98) (Figs 4. and 5). However, this HET difference effect on growth P_2_−P_1_ was much better – yet in opposite ways – correlated with CON or HET effect on survival in P_2_ for “larsm/reloc/crown” with *r* = 0.367 and −0.516 respectively (*P* = 0.023 and 0.001), and for “larsm/remov/crown” with *r* = −0.061 and −0.602 respectively (*P* = 0.72 and < 0.001). Differences in HET growth effects P_2_−P_1_ on survival as the sums of CON and HET effects on survival in P_2_, were also significantly negatively correlated for P_2_ (*r* = −0.541 and −0.331, *P* < 0.001 and 0.043, respectively for “larsm/remov/crown” and “larsm/reloc/crown”), but not P_1_ (*r* = −0.039 and −0.160, *P* = 0.82 and 0.34).

**Fig. 4.**
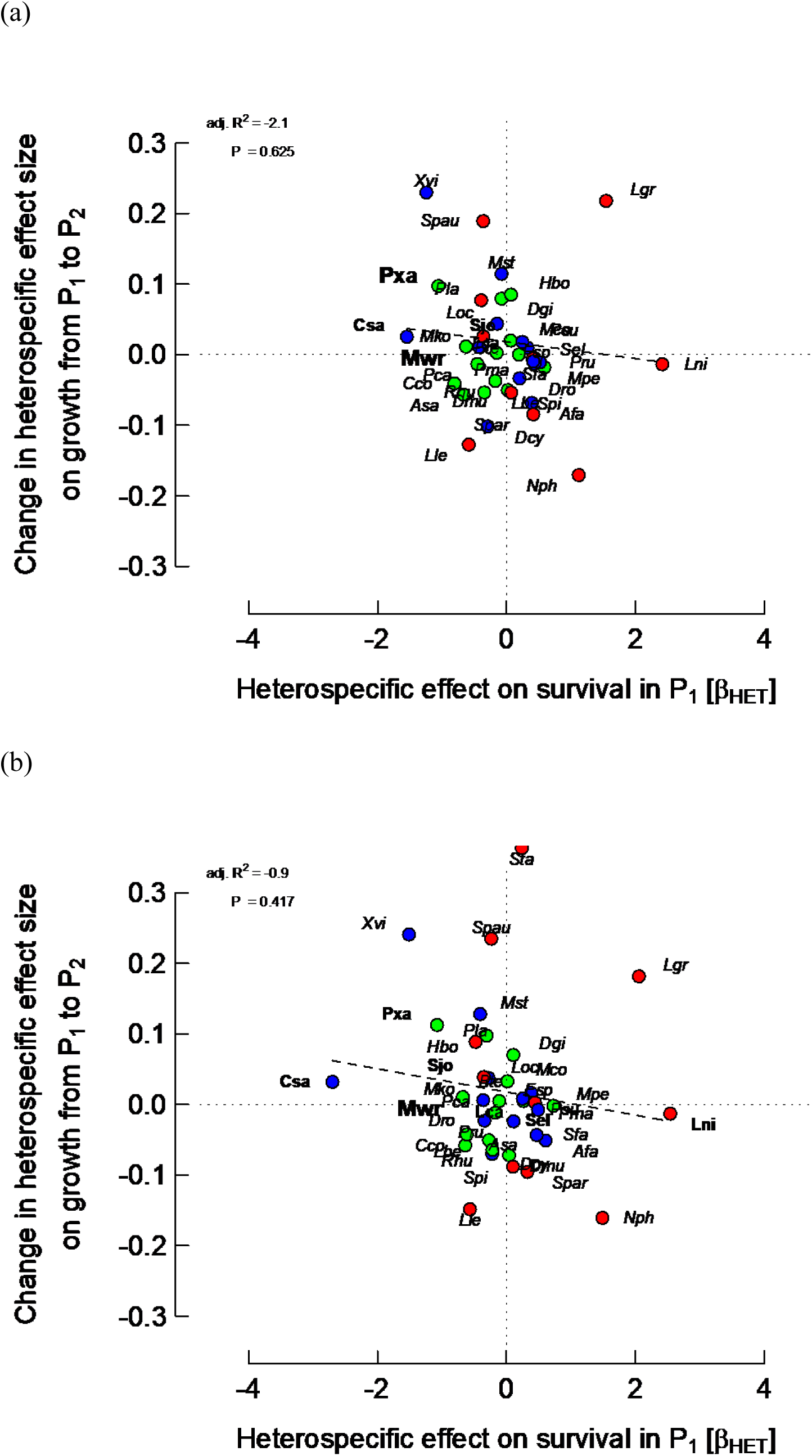
Relationships between *differences* in heterospecific (HET) effect sizes on growth rates between periods (P_2_ − P_1_) and HET effects on survival in period 1 (P_1_; β_HET_ = change in log[ODDS] per unit change in log(1 + sum(HET*_ba_*))) for the 38 species, from spatially-extended neighbourhood models with size (*ba*) and two neighbour terms (HET and CON). Points of focal trees having points of bigger neighbours within their zone of influence were either (a) removed, or (b) relocated. Details are the same as for Fig. 2, except for species’ labels: larger font, HET effect on survival in P_1_ with *P* < 0.01.

**Fig. 5.**
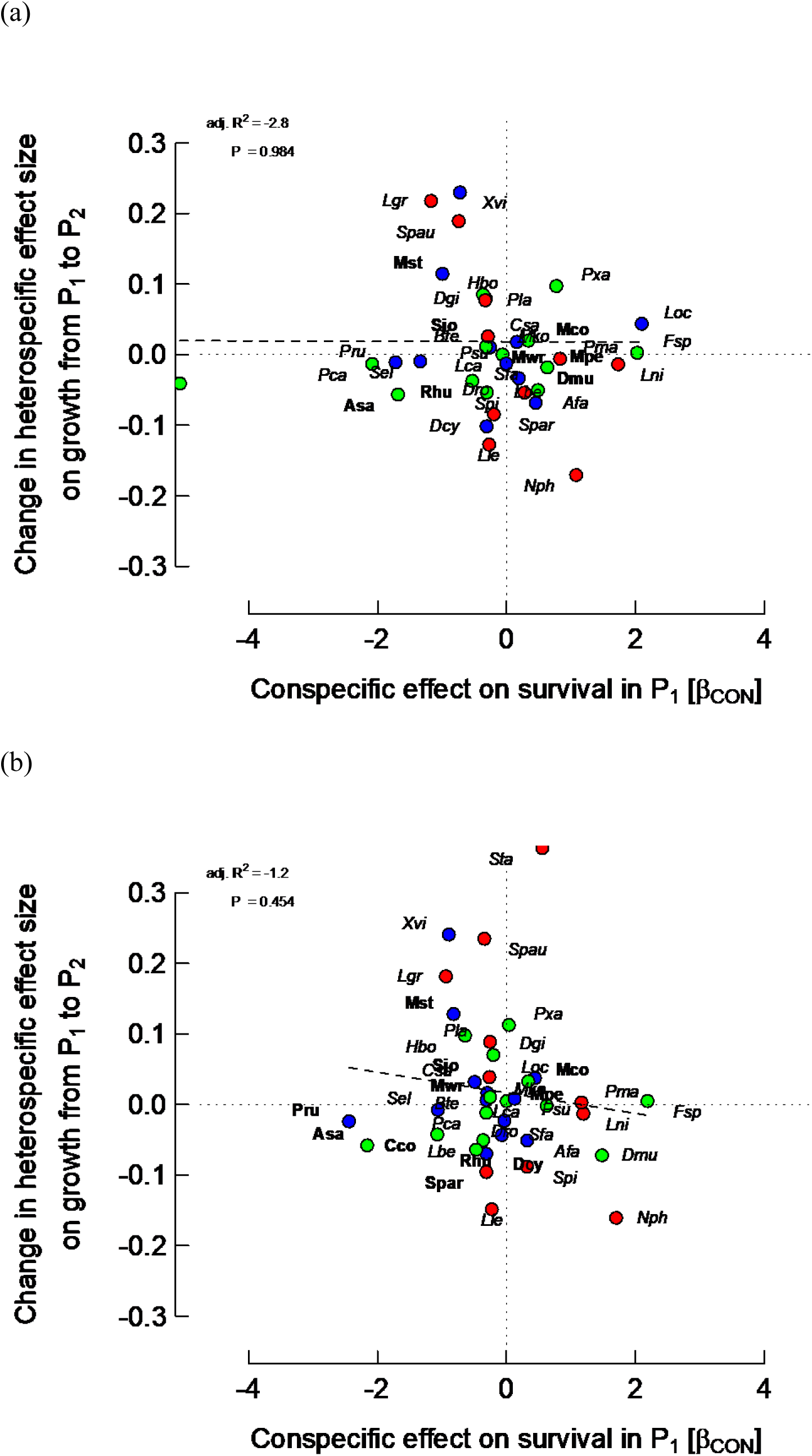
Relationships between *differences* in heterospecific (HET) effect sizes on growth rates between periods (P_2_ − P_1_) and CON effects on survival in period 1 (P_1_; β_CON_ = change in log[ODDS] per unit change in log(1 + sum(CON*_ba_*))) for the 38 species, from spatially-extended neighbourhood models with size (*ba*) and two neighbour terms (HET and CON). Points of focal trees having points of bigger neighbours within their zone of influence were either (a) removed, or (b) relocated. Details are the same as for Fig. 2.

Community level graphs using non-spatial models that complemented the two spatial ones in Figs 2 to 5 showed no trends or significance for a dependence on effect on survival (CON or HET) in P_1_ (Appendix 1: Figs. S3 and S4). Further, community-level graphs that used simply CON effect on growth versus CON effect on survival within one period, P_1_ or P_2_, also showed no significant relationships (Appendix 1: Fig. S5). The interesting trends happen, therefore, when *difference* in growth effect between P_1_ and P_2_ is related to effect on survival in P_1_ (CON and HET) using a *spatial* model.

Within the best-fitting spatial model, “larsm/reloc/crown”, CON effect on growth in P_1_ was significantly and positively correlated with that in P_2_, and also positively with HET effect on growth in P_1_ (*P* < 0.001) yet less strongly with HET effect on growth in P2 (*P* <0.10) (Appendix 1: Table S5). Likewise, CON effect on growth in P_2_ correlated positively with HET effect on growth in P_1_ and P_2_ (*P* < 0.05), but HET effects on growth in P_1_ and P_2_ were much less strongly correlated (Appendix 1: Table S6). CON and HET effects on survival in P_1_ and P_2_ were all generally poorly and insignificantly correlated with one another, except for CON effects on survival in P_1_ with HET effects on survival in P_1_ (positive). Stronger was the correlation between HET effect on survival with the same on growth in P_2_ (*P* < 0.01). Between survival and growth variables few were significantly correlated apart from CON survival in P_1_ with HET effect on growth in P_2_ (positive) and the same with HET effect on survival in P_1_ (negative). The model “larsm/remov/crown” had a similar pattern of correlations with an even stronger negative correlation for HET survival and growth in P_2_ again (Appendix 1: Table S6).

The relationship between CON difference effect on growth P_2_−P_1_ and CON effect on survival in P_1_ taken as a linear regression, i.e. assuming now a dependence of growth effects on survival effect, indicated that the best-fitting community-level plots were for “larsm/reloc/stem” and “larsm/reloc/crown” (*P* < 0.05), and that removing overlapping canopy instead of relocating it led to much lower fits (*P* > 0.15) (Table 6a). Having “equal” instead of “larsm” canopy allocations led to also weak, marginally significant fits. Restricting the regressions though to those 24 species which had CON effects on survival significant at *P* < 0.1 (no-decay mode), led to much stronger fits than for 38 species, with adjusted and predicted *R*^2^-values up to almost 30 and 16% respectively, maximal for the ‘larsm/reloc/stem’ and ‘…/crown’ spatial forms (Table 6b).

**Table 6.** Dependence of the difference in conspecific (CON) effect sizes in growth rates between periods (P_2_ − P_1_), for the different non-spatial and spatial models using linear distance decay, on the CON effects on survival in period 1(P_1_), for (a) all 38 species, and (b) the 24 species for which CON effects on survival in P_1_ were significant at *P* < 0.1 (see text for details). The models involved basal area (*ba*) and one or both neighbourhood terms (CON + HET) and, in the spatial case, one of the eight different forms of crown extension (as structured in Table 2). The final effects sizes came from model averaging (as in Appendix 1: Table S2).

| (a) 38 species |  |  |  | Periods 1 and 2 |  |  |
| --- | --- | --- | --- | --- | --- | --- |
| Model | *ppsqmca* | covered | pos. | adj. $R^2$ | $P$ | pred. $R^2$ |
| ba + ALL | – | – | – | 7.7 | 0.051 | -0.2 |
| ba + HET + CON | – | – | – | -0.5 | 0.369 | -21.4 |
|  | equal | removed | stem | 2.1 | 0.190 | -5.5 |
|  | “ | “ | crown | 2.2 | 0.182 | -5.9 |
|  | “ | relocated | stem | 6.4 | 0.068 | -3.4 |
|  | “ | “ | crown | 6.1 | 0.073 | -3.6 |
|  | larger > smaller | removed | stem | 0.2 | 0.305 | -7.1 |
|  | “ | “ | crown | 1.6 | 0.215 | -5.7 |
|  | “ | relocated | stem | 11.8 | 0.020 | 2.0 |
|  | “ | “ | crown | 10.6 | 0.026 | 0.7 |

| (b) 24 species |  |  |  | Periods 1 and 2 |  |  |
| --- | --- | --- | --- | --- | --- | --- |
| Model | *ppsqmca* | covered | pos. | adj. $R^2$ | $P$ | pred. $R^2$ |
| ba + ALL | – | – | – | 1.5 | 0.260 | -22.3 |
| ba + CON + HET | – | – | – | -1.5 | 0.422 | -31.3 |
|  | equal | removed | stem | 10.5 | 0.067 | 1.3 |
|  | “ | “ | crown | 7.1 | 0.111 | -30.1 |
|  | “ | relocated | stem | 15.7 | 0.031 | 3.1 |
|  | “ | “ | crown | 15.7 | 0.032 | 2.7 |
|  | larger > smaller | removed | stem | 3.9 | 0.178 | -9.3 |
|  | “ | “ | crown | 6.8 | 0.116 | -8.0 |
|  | “ | relocated | stem | 26.0 | 0.006 | 12.9 |
|  | “ | “ | crown | 25.9 | 0.006 | 12.8 |

In the community-level graphs, neither over-understorey status nor spatial patterning of trees (following could explain differences and trends in the CON and HET effect (Appendix 6 for detailed results: Table S1 and Fig. S1). However, in P_1_ though not P_2_, both CON and HET effects were significantly negatively correlated with stem relative growth, recruitment and mortality rates from early plot census analyses, i.e. species with strong negative effect values had also fast growth and population dynamics (Appendix 6 for detailed results: Table S2.).

A case could be made for excluding *N. philippinensis* and *A. sanguinolenta* (two of the five species whose changes in effects with radius were unusual) from Fig. 2b but that would have moved the fitted line only very slightly upwards with a similar slope (the points become a little more positive for CON effect on survival in P_1_). In conclusion, the final set of 38 species’ values appear quite robust for the community-level analysis. At this community level, correlation or regression of species’ slopes versus CON or HET coefficient was not feasible due to high skew and strong leptokurtis respectively.

### Spatial and non-spatial models compared

The community-level graphs of difference in CON effect on growth P_2_−P_1_ versus either CON or HET effect on survival, particularly for the ‘larsm/crown/reloc’ model (Figs 2b and 3b), showed stronger and more significant relationships than those with non-spatial models (Appendix 1: Fig. S3a, b). Correlation between CON effects on growth in P_1_ and in P_2_ between the two models were both strong (*r* = 0.802 and 0.848 respectively, *P* ≤ 0.001), but for CON and HET effects on survival in P_1_ they were weaker (*r* = 0.570 and 0.567, *P* ≤ 0.001). This differences between the two community-level plots is more the likely due to these CON and HET effects on survival.

To come closer to understanding the reasons for these differences, graphs of CON effects on survival in non-spatial and spatial models, and the same for HET effects, showed that the non-spatial model was more prone to serious outliers away from the general linear trends than the spatial one (Appendix 1: Fig. S6). Indeed, for CON effects one species, *Polyalthia rumphii*, had an extremely large negative value, and for the HET ones three species, *P. rumphii*, *P. sumatrana* and *Syzygium tawaense*, had unusually high positive values. Clearly these points created considerable leverage and were the main causes for the lack of agreement in the spatial and non-spatial community-level graphs. The probabilities associated with the *t-*values for these effect sizes coefficients in the individual species’ regressions were all large (0.41; 0.42, 0.88 and 0.98 respectively), i.e. insignificant, indicating large uncertainties about the estimates. Otherwise, high *P*-values were mostly attached to the coefficients close or at zero in Figs 2b and 3b, that is not distinguishing them much from null effects. *Cleistanthus contractus* and *Ardisia sanguinolenta*, both moderately separated from the main cluster of points in Fig. S4a in Appendix 1 had effect sizes that were significant or marginally so (*P* = 0.02 and 0.09). These subtle differences in reliability are not quite so apparent from fonts applied to species codes in Fig. S4. In passing, the remarkable species *S. johorensis* had outlying positive values for its difference in CON effect on growth P_2_−P_1_, for both the non-spatial and spatial models, but that was accounted for by the highly significant large effects in P_1_ (*P* ≤ 0.001) despite the corresponding effects in P_2_ being very close to zero (*P* > 0.05).

Omitting the above three outlying species resulted in stronger correlations between difference in CON effects on growth P_2_−P_1_ and CON effect on survival in P_1_ for the non-spatial (*r* = -0.313, *P* = 0.067) and spatial (*r* = -0.366, *P* = 0.031) models. For HET effect on survival in P_1_ the improvement was correspondingly even better (*r* = −0.430 and −0.364, *P* = 0.010 and 0.031). The main reason, however, that the spatial models were slightly superior overall than the non-spatial ones despite the more rigorous selection of species’ estimates, was that a group of five species, *D. muricatus*, *F. splendidissima*, *Lithocarpus niewenhuisii*, *Mallotus penangensis* and *Neoscortechinia philippinensis*, became more spread in their increasingly positive CON effects on survival in P_1_. A similar case for positive HET effects on survival in P_1_ is less strong though, involving just *L. niewenhuisii, N. philippinensis*, and *Lithocarpus gracilis*. Compared with non-spatial models, spatial models seemed to emphasize more positive rather than negative effects on survival. The two models, non-spatial and spatial, became more similar with the 35-species analyses.

Taking the axes coordinates for the spatial and non-spatial graphs of difference in CON effect on growth P_2_−P_1_ versus CON effect on survival in P_1_ as two 35 x 2 matrices, Procrustes rotation (package `vegan` in R; Mardia et al. 1979, Oksanen et al. 2019) highlighted strong agreement between the models, rescaling the spatial matrix to the target non-spatial one gave a correlation coefficient of 0.423 (*P* = 0.011; tested with 999 randomizations). Repeating this procedure for HET effects on survival in P_1_ led to a Procrustes correlation of 0.863 (*P* = 0.001). Thus, comparison of HET-based community graphs led to better model matching than did CON-based ones.

### Randomization of neighbourhoods

Across the 48 species, randomized mean CON effects on growth in P_1_ and P_2_, and their differences P_2_−P_1_, as well as CON effects on survival in P_1_ and P_2_, were mostly not significantly different from zero, judged by their confidence limits calculated as ± 3 SE (Appendix 7: Fig. S1). Cases of significance occurred more often among the 10 excluded species, and particularly for survival effects: in these cases, the limits were usually much larger than for the 38 retained species (see Appendix 1: Table S1). Retrospectively, the species selection was therefore well supported.

Considering only the 38 selected species, the mean observed effects were significantly different from the randomizations (i.e. they lay outside of the ± 3 SE limits) either positively, not or negatively for 5, 7 and 26; 8, 11 and 19; and 17, 11 and 10, of them for CON effects on growth in P_1_, P_2_ and P_2_−P_1_ respectively. The corresponding numbers for CON effects on survival in P_1_ and P_2_ were 8, 10 and 20; and 8, 19 and 11 (Appendix 7: Fig. S1). These frequencies show clearly that CON effects on growth in P_1_ and P_2_ were negative for a majority of species but differences moved to being mostly positive. Likewise, CON survival effects in P_1_ were in the majority negative too, but in P_2_ more species’ effects were insignificant, and positive and negative effects were more similar in frequency. Numbers of differences (positive, negative or null) among the other 10 species are uninformative given the statistical grounds for these species’ exclusion.

Defining moderate limits as being 0.06 to < 0.1 and 0.6 to < 1.0 for growth and survival effects respectively and corresponding large as being ≥ 0.1 and ≥ 1.0, among the 38 selected in P_1_ and P_2_ species such moderate and large limits for growth effects were moderately frequent (28 of 2 x 38 combinations, 37%). For the difference in CON effects P_2_−P_1_, 18/38 species (47%) had medium and large differences, whilst for survival ones they were similar (24/76, 32%). For the excluded 10 species, a majority of limits, for both growth and survival, were moderate or large (27 of 4 x 10, 67.5%).

HET effects on growth in P_1_ and P_2_, and their difference P_2_−P_1,_ plus HET effects on survival in P_1_ and P_2_, were mostly not significantly different from zero, judged by their confidence limits calculated as ± 3 SE (Appendix 7: Fig. S2). As with the CON effects, limits (± 3 SE) were much larger for the 10 excluded than the 38 selected species.

Of the 38 species, mean observed HET effects differed significantly from the randomization means positively, not or negatively for 4, 8 and 26; 2, 8 and 28; and 14, 9 and 15 of them for growth in P_1_, P_2_ and P_2_−P_1_ respectively. Thus, HET effects on growth were again predominantly negative in both periods. The numbers for survival in P_1_ and P_2_ were correspondingly 13, 10 and 15, and 18, 9 and 11. Hence for growth ± or 0 cases were rather similarly distributed over the 38 species for HET as for CON.

The strengths of the HET differences between empirical and randomized means, calculated in the same way as for CON effects, were very similar to the latter: 28/76 (37%) of species with medium and large differences in P_1_ and P_2_, 14/38 (37%) for P_2_−P_1._ For survival, medium and large HET effects differences in P_1_ and P_2_ formed 22/76 cases (29%). Among the excluded species these differences were also frequent with 20/40 (50%).

Putting CON and HET effect differences in comparison, significant negative CON and HET effects were in the majority for growth in P_1_ and P_2_; and with CON a majority positive, but HET more evenly distributed, for P_2_−P_1_. For survival in P_1_ and P_2_, CON effect differences were also predominantly negative in P_1_, yet fairly evenly positive or negative in P_2_ with most non-significant. HET effect differences showed the converse however, being evenly negative, not and positive in P_1_ yet predominantly positive in P_2_. For many species, there was evidently a shift in sign of survival differences between P_1_ and P_2_. Due in part to their sampling unreliability, differences for the excluded 10 species were often more pronounced for CON than HET (Appendix7: Fig. S2 cf. S1).

Simulating the regression of difference in CON effects on growth P_2_−P_1_ versus CON effects on survival in P_1_ 100 times using the randomizations, for all 38 selected species, and ‘larsm/reloc/crown’ spatial model, resulted in 62 slopes that were positive and 38 that were negative. Seven lines were individually significant at *P* < 0.05, two with negative and five with positive slopes (Appendix 7: Fig. S3a). Just one *t*-value (of −3.2) for the slopes was more negative in the randomizations than in the observed relationship (*t* = −2.3). This supports the inference (on the basis of a two-tailed null hypothesis) that the relationship in Fig. 2b is statistically robust at *P* < 0.05 level (arguably at *P* ≤ 0.02) since it lies outside of the 95% (or 98%) confidence envelopes of *t*-values under the null hypothesis of randomly positioned neighbourhoods. These randomizations will have removed any influences of spatial aggregation and therefore local dominance of species.

Repeating the community level simulation with just the 24 species with significant CON specific effects on survival in P_1_, resulted in 59 slopes that were positive and 41 that were negative. Seven lines were individually significant at *P* < 0.05, two with negative and five with positive slopes (Appendix 7: Fig. S3b). Neither of the negative *t*-values of the slopes (both −2.2) was more negative in the randomizations than in the observed relationship (*t* = −3.0), supporting the inference that the relationship was statistically robust here at *P* < 0.01.

The proportion of slopes of the difference in CON effects on growth P_2_−P_1_ versus HET effects on survival in P_1_ from the 100 randomizations were 46 negative and 54% positive, with 12 simulation lines significant (*P* ≤ 0.05), five negative and seven positive (Appendix 7: Fig. S3c). Four *t*-values for slopes (−2.8 to −3.3) were more negative in the randomization than in the observed relationship (*t* = −2.3), indicating only significance at *P* < 0.1 on a two-tailed basis. Why there was an imbalance of positive to negative slopes for CON (∼60:40) compared HET (∼50:50) under randomization remains to be explored.

Again, considering the 24-species community-level HET relationship, 50 and 50 of the slopes were respectively negative and positive and 14 lines were individually significant (*P* ≤ 0.05), seven negative and seven positive (Appendix 7: Fig. S3d). The empirical regression, accounting for very similar variance as for the relationship with 38 species, had (the) seven randomized slopes all more negative than that for the observed relationship (*t* = −1.8), suggesting that H_0_ might be rejected only at *P* ≤ 0.2. Narrowing the species considered from 38 down to 24, had an opposite effect for HET than it did for CON relationships in terms of improved statistical fitting (worsening versus improving respectively). Overall, the strength of the HET relationship was therefore much less significant (indeed non-significant at *P* > 0.1) than that for CON (significant at 0.02 ≤ *P* < 0.05) based on the randomization testing.

Using the mean effects from the randomizations, difference in CON effects on growth P_2_−P_1_ vs CON effect on survival in P_1_, for the 38 species — plotted in a similar way as for empirical data in Fig. 2b, had a positive correlation (*r* = 0.360, *P* = 0.026). However, excluding one clear outlier (*Hydnocarpus borneensis*), the correlation was then closer to zero (*r* = 0.080, *P* = 0.640), as should be expected from an effective randomization procedure. Had the number of randomizations been higher than 100, this and maybe other outliers would have been less important.

## DISCUSSION

### Spatial and non-spatial models

Difference in CON effect on growth P_2_−P_1_ was significantly *negatively* correlated with both CON and HET effect on survival in P_1_, but not in P_2_. Conversely, the difference in HET effect on growth P_2_−P_1_ was correlated *positively* with CON, yet negatively with HET, survival in P_2_ — but not in P_1_. There was therefore a part reversal of CON and HET effect associations over time. CON and HET effects on growth were positively correlated with one another in both P_1_ and P_2:_ CON and HET effects on survival, however, were positively correlated only in P_1_. Hence, CON and HET effects on growth, and on survival, appear to have been coupled in P_1,_ and then decoupled in P_2_. The randomization tests showed a high statistical confidence in the relationship between difference in CON effects on growth and CON effect on survival (*P* ∼ 0.02), but a similar relationship for HET effects was not nearly so robust (*P* > 0.1). Whilst HET effects evidently had a role in the tree neighbour interactions in P_1_ and P_2_, the main results concern the CON effects: conspecific interactions might be seen as being embedded in a diffuse matrix of heterospecific ones. Whilst the ‘larsm/reloc/crown’ spatial model was marginally the best supported realization of zone-of-influence competition, the concept and alternative models may have interpretational difficulties, highlighting the almost intractable complicatedness of diverse forest tree-tree interactions.

The models used in this paper defined neighbours as trees lying with a radius of 20 m and with *gbh* greater or equal to that of the focal one. For the majority of understorey species this meant that their focal trees often had few conspecific neighbours, especially if the focal trees themselves were ∼50-100 cm *gbh,* because these species rarely ever attained sizes of > 50 (or even 100) cm *gbh*. The opposite was the case though for overstorey species: for them conspecific neighbours were often far larger, and sometimes included the largest trees in the plots. On a simple biomass basis, then, CON effects on understorey species were expected to be rare and not as strong as the commoner overstorey ones, although both would be subject to similar levels of HET neighbour basal area. The effects of large-canopy conspecific trees would be even higher when the adults were aggregated (Newbery and Stoll 2013). In addition, removal and relocation of parts of crowns in spatial models was affecting mostly sub-canopy overstorey trees, those that on the one hand were being overlapped by upper canopy and emergent trees’ crowns, and on the other hand were remaining still large enough to make major contributions to CON and HET basal areas. Zone-of influence adjustments, according to storey position and size-class (*gbh*) frequency distribution, determined differences in how spatial and non-spatial models were operating for each species. The influence of adjustments on regression fits was weaker for under-than overstorey species. But then most understorey species would not be expected to be plastic in their crown adjustment to move towards light, because they are shade-tolerant trees, and drought-tolerant ones would only be temporarily exposed to higher light levels to have had insufficient time to change crown position before the canopy closed again.

For a neighbourhood model involving above-ground architectural traits, allometric regressions of crown area versus *gbh* would ideally have been better constructed for each of the 38 tree species, had sufficient data been available. Combining all species led to an averaging of crown areas across different species in each *gbh* class, e.g. a small tree of a dipterocarp (overstorey) and one of a euphorb (understorey) species would have been equivalent. Height was not involved in the crown overlap (removal/relocation) calculations: it was tacitly assumed that a tree with a large *gbh* was always higher than, and overlapping, a small adjacent one. This had important consequences for the crown adjustment algorithm. Crown depth and volume were not involved either. The simple non-species-specific allometric approach may, therefore, have distorted the true variation within and between species. The basic model, whilst being in some ways more realistic in the incorporation of crown area at all, may have introduced complicated biases when these crowns did not match well with each species’ ecological-defined height-diameter-crown area relationships. Spatial models adjusted crowns by removal and relocation of parts of them when overlap occurred. If this really was happening in the forest, (Sterck et al. 2001) would have incorporated them when making their crown measurements. So to some extent natural crown adjustment was already in the allometric equation. In this connection, the influence of coordinates of focal tree stem versus those of crown centroid was barely detectable in the outcomes of the spatial models.

The algorithms used for adjusting crown (and root system) overlap came nevertheless at a cost to some realism of the spatial models. Non-spatial models, with basal area at a distance from the focal tree placed at a neighbour tree’s centre, and the spatial models with Δd = 0 where crown and root system extents were determined (without any adjustments) by the common allometric equation, present two well-defined ends of a scale in crown extension, none to full. However, once Δd_CON_ or Δd_HET_ were allowed to increment > 0.0, adjustment meant potential removal or relocation. When overlap of smaller trees however was complete, either by one larger crown, several together, or one causing relocation of a less large one in a domino-manner, they could disappear as neighbours. This was likely to happen often because firstly the predicted LAI was close to 2.5 when Δd was 0, and secondly, understorey species’ trees from their ecologies are almost always shaded by others, especially in P_1_. The analysis of Δd-values selected by the best fitting models showed that in the main Δd was not 0.0. Even though the C_2_ two-term model was used, many understorey species had reduced CON basal areas because when small and fully shaded from above, they were still larger in *gbh* than the majority of the focal trees. Shading hierarchy paralleled storey structure, and may have given an undue bias to CON effects of overstorey species (as was the selection in Stoll and Newbery 2005).

In building the nearest-neighbour models, the focal tree was a point, with no crown extension or adjustment. Another, but computationally far more demanding approach, would have been to allow focal trees to have irregular crowns, and then find CON and HET basal area for each of the allocated points, and integrate basal areas per tree. But then focal trees themselves would be susceptible to disappearance if they were completely overlapped, presenting a dilemma. The random positioning of points in crowns at the start, before any adjustments for overlap, was also done just once. Had this step used multiple randomizations, then points allocation, adjustments, and crown shapes would have been allowed to vary, and thereby provided more robust mean effect sizes. However, this second potential extension involved prohibitively long computation times. Catering for these two fine-scale (within-crown) sources of variability may not have affected qualitative outcome of the analysis of the empirical data too much but it would have allowed for some modelling uncertainties to be taken into account. The many radial increments times Δd levels led to large numbers of very similarly-fitting models, particularly as points in crown area (*ppsqmca*) were from single crowns as neighbours and very small changes in the fitted coefficients came from the radial points moving across at 1-m increments.

Using spatial extension posited that the statistical modelling would move a step closer to forest realism in that above- and below-ground allocation of (neighbours) biomass under the symmetry/asymmetry of competition within the zone of influence would capture the CON and HET influences better than a non-spatial model. Neighbourhood models for the individual species highlighted, though, that whilst involving spatial models did more often explain focal tree growth (not survival) better than non-spatial ones, those models with a *ppsqmca* allocation proportional to tree basal area (‘larsm’), either with relocation or removal of overlapping parts of crowns, were not more often better than those with an even allocation (‘equal’). The former introduced an asymmetry, or non-linearity, in neighbour interactions, the latter not: however, increasing Δd also introduced degrees of asymmetry so that as this parameter was increased, removal or relocation led to ‘equal’ points distributions becoming less equal. Conversely, with the ‘larsm’ model form a larger shading crown could cause a lower crown to be adjusted, which when through relocation would increase the lower one’s *ppsqmca*.

Conspecific effects of neighbours on growth in P_1_ were more weakly (though still significantly) related to plot-level BA for spatial than for non-spatial models, and likewise for differences in CON effect on growth P_2_−P_1_, but there was no influence of model form for CON effects versus BA in P_2_. Conspecific effects of neighbours on survival in P_1_ and P_2_ were also independent of BA, whether the model form was non-spatial or spatial. The relative strengths and directions of the relationships for CON effect on growth versus plot BA in P_1_ and P_2_, for non-spatial models, is the same as reported before (Stoll and Newbery 2005, Newbery and Stoll 2013). If spatial extension (in the form of plasticity of crown size and overlap, and location) were supposed to simulate competition for light between neighbours better, and be a basis for the proposed negative density-dependence of CON effect size on plot-level species abundance (i.e. more abundant species in the plots with higher BA had associated with them stronger more-negative CON effects on their small trees), the reduced fitting would imply either that any driving causal influence of abundance *per se* (plot BA) was not happening through asymmetric competition for light, or that non-spatial models without plasticity overestimated the ‘true’ effect of negative density-dependence. Alternatively expressed, plasticity allowed crowns to be distributed closer to how they are thought to compete for light yet removed the conspecific negative density dependence (Stoll et al. 2002).

The reduced fits of the spatial compared with the non-spatial models can be explained best by the application of Jensen’s Inequality (Jensen 1906, Ruel and Ayres 1999) (see also Ross 2014), and most simply for Δd = 0, because the 1/d weighting of basal area is a concave function of d. The mean of the inverses of distances from the focal tree location to points in the neighbours (circular) crown will always be greater than the inverse of the distance from the focal tree to the centre of the neighbour one (being the mean coordinate of all points in that crown). Consider a neighbour of crown radius 1 m whose centre is 3 m from a focal tree. The closest point on the circumference of the crown is 2 m and the furthest point 4 m from the focus. The mean of their 1/d values is (0.5 + 0.25)/2 = 0.375, but 1/d for the centre of the crown is 1/3 = 0.333. A more extreme example: a crown of 5 m radius has its centre 7 m from the focal tree. The corresponding means of inverse distance and distance from centre to focus are (0.5 + 0.083)/2 = 0.292 and 0.143. Thus if *ppsqmca* are allocated under ‘larsm’ at random within crowns equal in number to the neighbour tree’s basal area in cm^2^, a spatial model will always give a higher weighting to that neighbour’s basal area compared with the non-spatial model with all basal area at the tree’s centre. It follows that the spatial models will fit less well than the non-spatial ones because the larger CON and HET basal areas are moved more positively, away from zero, on the X-axis which leads to the dependence of growth or survival (the slope in the regression) to be less steep. Nevertheless, the model fits for ‘no’, ‘lin’ and ‘squ’ distance weightings differed rather little (Appendix 1: Table S2) suggesting that the rescaling caused by the different weightings was, though important, small in its influence. The logarithmic transformation of CON and HET basal areas in the models would have dampened the differences between types of distance weighting.

The same arguments apply to the ‘equal’ allocation of crown points, but the influence of the inequality will be generally less because *ca* is proportional to *ba*^1/2^ and not *ba*. Adjustments for crown overlap by removal or relocation emphasized the influence of Jensen’s Inequality. Adjustment of points by removal would leave them on average both closer to or further away from a focal tree than before (i.e. to either side of a shading larger crown), or by relocation increase the average the inverse distance weighting further. Since logistic regressions are more sensitive to changes in range and skew in predictor continuous variable (CON and HET basal area weighted by 1/d) than gaussian normal ones, the loss in fit moving from non-spatial to spatial models will be greater for CON effects in survival than CON effects on growth especially. For HET effects the differences are ameliorated by the heterospecifics making up that large matrix of neighbour trees that shift about in their canopy positions between one another under adjustment much less than is experienced for conspecific trees.

By the same token, that the relationship between CON effects on growth in P_2_ versus BA fits were barely altered under spatial extension, and it was shown that CON effects were overall relaxed in this period compared with P_1_ (they became less negative, Newbery and Stoll 2013), then another factor such as competition for, or utilization of, nutrients might account better for the negative density-dependence — but only if patterns of nutrient acquisition are not following light ones in the same way, i.e. the root systems are crown-size and -shape unrelated. That the slopes of relationships changed little between non-spatial and spatial modes, even a little less steep for the latter compared with the former, suggests that the lowered variance accounted for was mainly due to added variability coming from the common crown allometric equation being unsuitable for all species, the random allocation of crown points, and way overlap led to crown or root system removal or relocation. This together raises then the possibility that below-ground interactions were as or more important than above-ground ones in this forest.

These aspects all likely contributed to the poorer fits of spatial models than the non-spatial one for CON and HET effects against plot BA. It is further notable, that the regressions against plot BA and the community-level graphs showed no clear trends between overstorey and understorey species other than larger-stemmed overstorey species tending to be out on the extremes of the negative relationship and the understorey ones clustered at the centre. The spreading of a group of species to the other side (CON effects on survival in P_1_ being positive) is of considerable interest. Had light been the predominant factor a clearer storey-related pattern should have been more evident (see Newbery et al. 2011), especially in view that because of the disappearance of smaller crowns overstorey species were emphasized over understorey ones.

### Changes between periods and conspecific mechanisms

The evidence and arguments so far suggest that competition for light was not the main or sole driving factor behind CON and HET effects. Asymmetry of competition can be explained physically in terms of light interception and shading, and yet the very strong asymmetry that the models invoked (Table 2) did not lead to significantly better model fits than with symmetry. By putting aside the outlying and statistically unreliable estimates of four of the species, the ‘larsm/crown/reloc’ spatial model scaled well on to the non-spatial one. Competition among root systems is normally expected to be much more symmetric than among leaves and crowns (neighbouring trees assessing and taking up resources in the soil in proportion to their respective biomasses), and with the involvement of ectomycorrhizas the interaction can be even more facilitative than competitive (Newman 1983). However, in competition modelling, resources below ground (principally for nutrients outside of dry periods), have often been assumed to have either a negligible role or to be operating similarly, or in proportion, to the light factor. This may be more likely for fast growing, colonizing or secondary forest growth, but is difficult to understand for a late end-succession or mature primary forest like Danum (Newbery et al. 1992).

There exists a problematic implicit assumption with the zone of influence competition modelling concept when defined by simple spatial extension of tree form and mass, its icon being crown area. Corresponding to the crowns, overlap and plasticity is directly and similarly implied for the root systems. The same effects of removal and relocation are *sensu lato* also implied for processes below-ground, vertically matching crown with root system adjustments, so that presumably smaller trees roots are respectively thinned or concentrated away from those of larger ones. But is this spatial model, even in its main components, realistic? Based on the differing physiologies of the tree parts, only a broad correlation between root system and stem/crown biomasses across tree sizes would be expected. The idea of ‘overlap’ remains importantly problematic. For a light competition it is readily interpretable but for root systems a tendency towards a symmetry or linearity of full intermixing in acquiring water and nutrients would be expected. The overlap Δd-value (up to 1.2 m) was to allow for an oblique shading zone; but for roots the equivalent is unclear – a depletion zone for water and nutrients that did not have daily or annual variation? The parameters, Δd_CON_ and Δd_HET_ were set to apply in the same way for all species, irrespective of their leaf size and density, branch structure, root distribution and corresponding ecophysiological differences.

The growth rate calculations in this paper did not specifically exclude all trees with ‘invalid’ measurements (see Lingenfelder and Newbery 2009), that is those where the point-of-measurement had moved slightly, stem (bark) condition changed or deteriorated, recording was inaccurate due to liana growth, etc. At the 1996 and 2001 censuses (end of P_1_ and P_2a_) close to 10% of growth estimates were invalid, and trees with invalid estimates had rates on average 47% lower than valid ones (Newbery and Lingenfelder 2009). However, excluding, in this paper, trees with > −3 SD deviations on the species’ *agr*-vs-log(*ba*) regressions will have caught most of the more extreme low-growth values. As a rough estimate then reliable growth rates were likely up to 4.7% underestimated, and the assumption has to be made for the present modelling that this small bias applied fairly evenly across periods and species and had little influence on the conclusions. The dependence of growth rates on the field recorded *gbh*s at two times is sensitive and prone to error and bias: and, this latter is more acute the more trees in the population are dying or close to death (with prior declining *rgr*’s). However, mortality rates (trees ≥ 10 cm *gbh*) were 27% higher in P_2_ than in P_1_ — 1.99 vs 1.57 %/yr, so ‘true’ rates would have been underestimated a little more so in P_2_ than P_1_, and thus the difference in CON effect on growth P_2_−P_1_ for species with the relatively higher within-period mortalities in P_1_ and P_2_ underestimated. The slopes of the lines in the community-level graphs of Fig. 2 are therefore slight underestimates. This issue of growth rate validity is crucial to evaluating how mortality depends on prior growth rate because assessing growth rate is very difficult when the stem itself is deteriorating just before death (Lingenfelder and Newbery 2009).

Compared with P_1_, the temporary decrease in soil water availability followed by increases in light levels to the understorey caused by the 1998 ENSO event in P_2_, differentially affected changes in species’ growth and survival rates between periods (Newbery and Lingenfelder 2004, 2009; Newbery et al. 2011). However, the extent to which differences in response between individuals of any one species were directly caused by the drought/light-change environment or were indirectly caused by their also-affected neighbours’ growth and survival rates, or both, depends on a highly complex set of spatial-temporal tree-tree interactions. Integrating across each species’ tree population results in simply their average growth and survival rates for each period. A species responding positively to the P_1_-P_2_ external change might be expected to become more competitive for both above- and below-ground resources and to thereby have stronger CON-HET effects on its neighbours. Conversely, a negative response could be because the neighbours are responding more negatively to the change and their CON-HET effects become weakened.

The community-level diagrams indicate that those species which suffered the largest CON and HET effects on survival from neighbours in P_1_ (i.e. their relative increase in mortality from this cause was highest), had the largest releases from CON — but not HET — effects on growth P_2_−P_1_. Conversely, those species that were little, or even positively, affected in their survival by CON and HET neighbours had either small releases or decreased negative effects on growth between P_1_ and P_2_. Expressed otherwise, with growth rates intermediate or even slightly higher in P_2_ than P_1_, yet more equally spread across species in P_2_, the more the suppressed species in P_1_ appeared to be released and the less suppressed ones hampered. The neighbourhood regression models only test though for CON and HET effects on growth rates *within* periods (separately), but they do not test for an interaction *between* CON and HET effects on growth rates and periods, which for the inferences drawn for this paper was assumed to be zero.

One potential explanation for the patterns in the community level graphs is that local thinning of focal trees in P_1_, due to the CON effects on growth and survival, might have relaxed competition between them in P_2_ and thus created the release in CON effect on growth. If that were to have operated the relatively small focal trees would have had to be very close to one another indeed to allow for intraspecific interactions to operate (mechanistically). This was generally not the case. Just one species, *D. muricatus*, possibly reached sufficient local densities in clusters on ridges (Newbery et al. 1999, Newbery and Ridsdale 2016). This species was most unremarkable on account of its position on the community-level graphs. A HET effect on survival in P_1_ however seems more plausible on spacing grounds, and so together it can be supposed that CON and HET basal area together contributed to any general density or thinning effect (Figs 2 and 3).

The physiological process by which prior slowed growth leads to the death of a tree, and how growth and mortality are actually recorded in populations over time, is fundamental to an understanding and interpreting the community-level relationship in Figs 2 and 3. For the Danum forest this was demonstrated by Lingenfelder and Newbery (2009) and Newbery and Lingenfelder (2009). If probability of tree mortality is generally continuously related to stem growth rate, that is a population is not divided into say two discrete classes where one is dying very fast independently of growth rate and the other very slowly (as in perhaps an age-related disease susceptibility situation), species with higher mortality rates will be expected to have on average (prior to death, and for those remaining and not yet dead) lower *rgr* than do species with lower mortality rates. It is important to note that the CON effects on survival in P_1_ are not alone determining mean survival rate of that species, but only how CON basal area increases or decreases it. The consequences of the CON effect for species with a very low compared with a moderate or high average rate of survival may be different.

If the main effect of declining growth on reduced survival can be translated to negative CON effect reducing growth and negative CON effect reducing survival — since growth rate and survival overall for a tree are in part determined by the neighbour effects, then a CON effect that reduces growth rate further should also have the consequence of reducing survivorship further. Hence, a ‘release’ in CON effect on growth P_2_−P_1_ will be larger (positive) for species with larger (negative) CON and HET effects on survival in P_1_ than those with smaller (positive and negative) effects on survival, because the more suppressed a tree is in its growth the greater the potential for release when conditions that caused the suppression are removed. That CON and HET effects on survival appear to be operating more in P_1_ than P_2_ is compatible with the thesis that small focus-sized trees in the undisturbed, closed and shady understorey will have lowered survival rates due to these relatively low light conditions, and their large competitors in the overstorey will exacerbate the situation by exerting increasingly larger negative effects on their growth.

The processes that resulted in the *release* of CON effects on growth in P_2_ may not have been the same one either that was linking CON effects on growth to CON (and HET) effects on survival in P_1_. The one in P_2_ was being largely driven by an external change in the environment leading to increased and variable light conditions within the understorey, whereas the one before in P_1_ was determined more by a steady environment and more closed, shaded internal-forest conditions. Thus under a trade-off in responses, shade-intolerant species would be dying most in P_1_ and yet responding (i.e. the survivors and new recruits) most to light in P_2_, whilst shade-tolerant ones would be less affected in P_1_ but, being weaker competitors under more lighted conditions they would continue to have lowered growth rates, CON effects on growth became minimal. That differences in CON but not HET effects on growth were strong, indicates that species-specific root processes might have been interacting positively with a growth-survival trade-off along the light gradient, and this thereby offers a route to explaining species’ idiosyncrasies at the community level. There would be the freedom of various individualistic conspecific processes to be operating. The idea is that it is not simply the difference in response to light that likely differentiated species (as shown in Newbery et al. 2011), but the interactive effect of CON-neighbours roots *on that* overall tree growth response to light measured.

Could the results be the outcome of random patterns and processes operating? The lines fitted on the community graphs go through zero with a negative slope. A completely random set of responses would settle around zero according to the central limit theorem; and this was supported by the randomization runs: on the other hand, a net outcome of interactions between neighbours would also result if species were balancing out their negative and positive effects, particularly when most of the neighbourhood of any one species is largely HET. A form of zero-sum game in the whole forest might be implied. It may be axiomatic that all interactions even out in a dynamic equilibrium: the CON and HET effects that some species on average experience as negative, others experience as being positive. What characterizes the differences in CON and HET effects on growth and CON and HET effects on survival in the two periods might be largely a question of chance where the individuals happen to be located with respect to their neighbourhoods. In mixed forest, areas will differ in local BA density either at random or with some degree of local clustering. High BA neighbourhoods create greater overlap of zones of influence and competition than low BA ones. If a shade-intolerant species should by chance happen on average to have more of its small trees in high BA patches it would be expected to show a strong CON and HET effect on growth and survival; and if in low patches, weaker effects.

Three aspects suggest that species responses were highly idiosyncratic because of the lack of any interpretative trends. Firstly, the arrangement of species in the community-level graphs in Figs 2 and 3 seem to have no clear explanation in terms of the under-versus overstorey classification, or population variables for P_1_ and P_2_ (indicating a growth-survival trade-off across species), or species’ reactivities to the drying event in P_2_, which were all useful in explaining changes in tree growth rates between P_1_ and P_2_ (Newbery et al. 1999, 2011; Newbery and Lingenfelder 2004). Further, there are no apparently excepting reasons for the outlying species’ points in Figs. 2 and 3, even for *Shorea johorensis* with its very pronounced relaxation of the negative CON effect on growth between P_1_ and P_2_. No common consistent explanation can be offered either for the group of five species whose CON effect of survival was positive (lower right portion of Fig. 2b).

Secondly, individuals of shade-intolerant tree species are expected to respond more negatively, in terms of their growth and survival, than those of shade-tolerant species as CON and HET basal areas increasing in their neighbourhood impose more shaded conditions for small trees. When the limiting (light) condition is removed, shade-intolerant species will respond faster than shade-tolerant ones – the link between mortality on growth being relaxed. However, the simple categorization into shade tolerant and light-demanding species (e.g. Whitmore 1984, Turner 2001) scarcely applies here with so few forest gaps and lack of secondary species. Several main understorey species respond to increased light and are apparently more drought-than shade-tolerant, and understorey trees of overstorey species show a wide variation in responses to changing light conditions (Newbery et al. 1999, 2011; Newbery and Lingenfelder 2004, 2009, 2017). Therefore, to refer to any ‘trade-off’ connected with life-history strategy is irrelevant and misleading for this forest. Although species may differ on average in their responses, a wide variation within species exists because of many other factors which interact with light, particularly nutrient and water availability.

Thirdly, the new evidence, using spatially extended models, does not dispel that CON effects were acting ‘by default’ (Newbery and Stoll 2013), i.e. they arose when conspecific species were clustered and large trees formed most of the neighbours of focal ones locally? The spatial pattern analyses in the present study concluded that degree of aggregation as a general factor for all species was unrelated to the trends in the community-level graphs. This does not mean though that it was not of *one* of a set of staggered factors leading to CON effects on growth and applied mainly to the large strongly clustered dipterocarps (Stoll and Newbery 2005). Randomization tests will not only have removed clustering but all possible linkages between, and complementation, of neighbouring root systems.

Since strong CON effects on growth contribute to lowered tree *rgr*, which when very low or at zero will usually result in tree death, the community level graphs in Figs 2 and 3 are representing a form of difference in CON effect on growth P_2_–P_1_ versus CON effect of growth in P_1_. And if CON (and HET) effects on growth (not growth rates *per se*) were randomly occurring in tree populations labelled as species, then negative slopes could be explained by the ‘regression-to-the mean’ phenomenon (Tversky and Kahneman 1974, Kelly and Price 2005). Random changes in CON effects between P_1_ and P_2_ would mean that groups with high CON effects in P_1_ would have lower values on average in P_2_, and their difference becomes closer to zero, and those with low CON effects in P_1_ work in the opposite way but also moving towards zero (the overall ‘mean’). The correlation between CON effect on growth in P_1_ with that in P_2_, for the non-spatial and spatial (‘larsm/crown/reloc’) models were 0.466 (*P* = 0.003) and 0.521 (*P* ≤ 0.001) respectively. So whilst the correlation was higher for the spatial model there was sufficient variation to cause a regression to the mean. That might *in part* explain the passing through the origin and the negative slope. Randomizations — which simply relocated tree positions —would be expected to show a similar trend, and they do, although the empirical line being just significant, suggests that a set of some real CON effects – perhaps just for the main large dipterocarps (Stoll and Newbery 2005) – were operating in addition to a resampling-over-time artefact.

We hypothesize that the negative slope in the community-level graphs of Figs 2 and 3 is in part determined by a gradient of strong-to-weak below-ground rooting processes allowed by different degrees of response to light-level change. The negative density dependence relationship of differentiating CON effects on growth depending on plot BA might then be better explained by below-ground rather than above-ground processes? Conspecificity, if it is real and not by ‘default’, must have mechanisms that are species-specific in order to operate: adults must be affecting juveniles of their own more than other species, and juveniles are only, or very largely, affected by their own, and not all, adults. Although not applying to all large-treed species, juveniles with ECMs may have been relieved in P_1_ of their dependence on close-by adults for carbon and nutrient transfer under the P_1_ light-limited conditions. With more light in P_2_ the focal trees became more autonomous as their own C input increased and this meant they could, though increase fine root and hyphal growth, acquire phosphorus and other elements by uptake and transfer more independently.

The analyses of the neighbourhood effects of large trees on small focal ones at Danum, over the two periods, presented a paradox. The negative density-dependence relationship based on the non-spatial model was weakened in spatial models due to averaging of inverse-distance weightings. If the latter with crown extension and plasticity is a truer representation of forest structure and tree-tree interactions than the former without it, then the strength of negative density dependence was over-estimated by Stoll and Newbery (2005). Yet the spatial models came with several provisos and limitations that questioned their full reality, in particular in the ways the conspecific tree interactions for understorey species were de-emphasized and those for overstorey species were over-emphasized.

Nevertheless, the community-level graphs relating differences in CON effects on growth P_2_−P_1_ to those on survival in P_1_ were slightly better for spatial than non-spatial models which suggested that the link between growth and survival responses to neighbours, and the change in the former under differing forest conditions (P_1_ to P_2_), were in part due to a dependence of survival on growth but also partly due to differences in relative importance of asymmetric and symmetric competition (potentially with facilitation via ectomycorrhizas). However, species of the Dipterocarpaceae and Fagaceae (likely also ECM) show no grouping on the community-level diagrams in Figs 2 and 3. To achieve some progress on understanding the mechanisms behind the tree-tree interactions it is essential to have detailed data on root distribution, growth and activity of each of the different species, and how allocation and plasticity in stems and crowns is related to those of root systems. However, for practical reasons this is going to be extremely difficult to achieve with the limited sampling techniques currently available. For now, the only rare available data for Danum are overall fine root biomass and dynamics estimates, undifferentiated at the species level (Green et al. 2005). Appealing to similar studies in other forests for support (and extremely few exist) is unlikely to be useful as the interpretation of forest dynamics at Danum is closely dependent on precise information about that forest and the plots recorded.

### Conclusion

The forest at Danum, and a large part of the surrounding region in Sabah, is subject to the influences of climatic variability (Newbery et al. 1999, Walsh and Newbery 1999), a main source of environmental stochasticity that drives forest dynamics (Newbery and Lingenfelder 2004). Events such as dry ENSO periods introduce marked changes in growth and structure, albeit over relatively short time periods, but ones that have much longer term effects (Newbery et al. 2011). Different species’ trees, depending on their size, local environment, and ecophysiology, respond individually and differently in their growth rates. Tree population structures move along trajectories until the next event disturbs them again (Huston 1994) and throw the interactions again into disarray (Richards 1996). The responses of the trees, particularly smaller ones, to their larger neighbours, in this changing environment, entails both the focal tree’s response to the variation in external conditions and the collective changes of the neighbours to it. Existence of conspecific negative density dependence operating on small-tree growth and survival was called into question by the present study because it was lost when moving from non-spatial to spatial models. Recent re-evaluation of data from three tropical tree recruitment studies have raised doubts too that this type of dependence is as important in forest dynamics as was previous contended (Detto et al. 2019).

The tropical forest ecosystem cannot necessarily be assumed to be in an equilibrium: there is neither reason nor evidence to show that the species populations measured over two decades (1986-2007) would have coexisted in similar proportions in the past, or that they will continue likewise to coexist in the future (Tokeshi 1999). Present dynamics, in terms of tree growth, recruitment and mortality, are determined by historical contingencies and site conditions. No stabilizing trade-offs among species need occur either beyond the simple physical constraint of maximum total forest biomass, and possibly a feedback determined within the system by under-overstorey guild structures (Newbery et al. 1992). In the absence then of empirical results it might be unwise to assume evolutionary strategies operating for root systems (see Dybzinski et al. 2011, McNickle and Dybzinski 2013). Environmental stochasticity, as pink/red noise in the Danum ENSO signal for instance (Newbery et al. 2011) adds in theory a potentially highly complicated mixing effect on tree-tree interactions. Furthermore, the continually varying species composition of neighbourhoods around individual (focal) trees, temporally and spatially creates a *neighbourhood stochasticity*, which is highly problematic to define, record, model, and use predictively. One clear realization of this was that the change in CON effects on tree growth moving from P_1_ to P_2_ appeared to be partly related to (correlated with) CON and HET effects on survival in P_1_, and yet it is difficult to explain this relationship satisfactorily using the recorded parameters of the species’ mean dynamics and population structures from the same periods.

## Acknowledgements

This work was part of the Royal Society of London’s SE Asian Rain Forest Research Programme 1985-2015. We thank the Sabah Biodiversity Council and the Danum Valley Management Committee for permission to continue our research at Danum, R. C. Ong (Sabah Forest Department) for host support, the late C. E: Ridsdale for field identifications and over-seeing taxonomic identifications with the Sandakan Herbarium, M. Lingenfelder for maintaining the plots database, and A. F. Karolus (DVFC) for field work assistance. For field inputs to the previous censuses, thanks go also to E. J. F. Campbell and M. J. Still (1986); D. N. Kennedy and H. Petol (1996), M. Lingenfelder (2001) and K. F. Poltz (2007). We appreciate the support of the Scientific Computing Centre at the University of Basel for facilitating the main statistical calculations. Two anonymous reviewers are also thanked for their valuable comments on the first submission.

## Author contributions

The idea of extending the models by crown extension was initially conceived by PS and then discussed through all the later stages between him and DMN. PS wrote the programs in R and ran the regression models providing the core results. DMN very largely wrote the paper, with inputs from PS. DMN made the ancillary statistical analyses, and developed the theory. Both authors read, discussed and corrected the various versions. The author contributions were overall roughly equal, and the order is therefore alphabetical.

## Data Accessibility

The raw input data input are available at the Dryad Digital Repository (DOI: 10.5061/dryad.q573n5tfx).

## List of Supplementary Materials

App. 1. Table S1. List tree species analyzed and their population sizes.

Table S2. Model fits: spatial versus non-spatial based on AIC.

Table S3. Complement to Table S2, with alternative reference model.

Table S4. Model variances and P-levels reached by individual species fits.

Table S5. Pairwise correlations across species’ model estimates.

Table S6. Cross-correlations of CON and HET effect sizes.

Fig. S1. Frequency histograms of species’ CON and HET Δd-values.

Fig. S2. Community-level graphs for non-spatial model complementing Figs. 2 and 3.

Fig. S3. Community-level graphs for non-spatial model complementing Figs. 4 and 5.

Fig. S4. Graphs for spatial model of CON effects on growth vs survival.

Fig. S5. Graphical comparison of non-spatial and spatial models for effects on survival.

Fig. S6. Frequency histograms of species’ CON and HET coefficients.

App. 2. Computer code (in R) for the calculations of crown distribution, points allocation, overlap and adjustment for the spatial models.

App. 3. Fig. S1. Frequency histograms of CON effects on growth for each of species analyzed.

Fig. S2. Frequency histograms of CON effects on survival for each of species analyzed.

App. 4. Tables of the effect sizes, statistical fits and significances, and best fitting radii, of the regression outcomes for all species, non-spatial and two spatial models, for growth and survival responses in Periods 1 and 2 (*four large Excel files*).

App. 5. Fig. S1. Relationships of CON effects on growth with best-fitting neighbourhood radius.

Fig. S2. Relationships of CON effects on survival with best-fitting neighbourhood radius.

App. 6. Analysis of the role of structure (Table S1), dynamics (Table S2) and patterning (Fig. S1) of the species studied in relation to CON and HET effect sizes.

App. 7. Fig. S1. Graphical summaries of all species’ randomization runs, for CON effect sizes on growth and survival in Periods 1 and 2.

Fig. S2. Graphical summaries of all species’ randomization runs, for HET effect sizes on growth and survival in Periods 1 and 2.

Fig. S3. Community-level graphs showing the lines from the randomizations which test-match Fig. 2b.

