## Supplementary material for "Including tree spatial extension in the evaluation of neighbourhood competition effects in Bornean rain forest": nncrexm appendices: App1_supportsn.pdf (from another version of this preprint: version 2, 2020-08-04)

Appendix 1: Table S1.

Full list of the 48 species in the two 4-ha plots at Danum combined, the first 38 of which (above the middle solid line) were selected for the more detail analyses and statistical modelling. Grounds for inclusion of the last of the 38, and exclusion of the other 10 species are explained in the main text. Abbreviations ('Abbr.') are used in figures for point labelling species, the matching eight-character codes from the 1986-2007 database are given for reference, with family codes. 'OUI' is the over/understorey index (scale 0-100; see Newbery *et al*. 2011: > 55, overstorey [red], 20-55 intermediate [blue], < 20 understorey [green]). 'BA $P_1$ and $P_2$', are the basal area abundances of each species at the plot level ($m^2$/ha; trees ≥ 10 cm *gbh*), in periods 1 and 2. 'N. trees' shows the numbers of focal trees (10 - < 100 cm *gbh*) that were potentially usable ('pot.'), those excluded ('excl.') where residuals in the *agr* vs. *ba* [individual focus tree basal area] were < −3 SD), and finally those accepted ('used'), for periods 1 and 2. 'N. dead' are numbers of focus trees that died by the ends of periods 1 and 2. The taxonomy follows the stand of the updated database of 2010. On tabulation, the archived data file will give the total counts of focal trees at the starts of $P_1$ and $P_2$ which equal the sums of the counts of potential and dead trees (e.g. APORFALC/$P_1$, *n*_total = 118 + 20 = 138).

| | | | | | | BA | | N. trees P1 | | | N. trees P2 | | | N. dead | |
|---|---|---|---|---|---|---|---|---|---|---|---|---|---|---|---|
| Genus | Species | Abbr. | Code | Family | OUI | $P_1$ | $P_2$ | pot. | excl. | used | pot. | excl. | used | $P_1$ | $P_2$ |
| *Aporosa* | *falcifera* | Afa | APORFALC | EUPH | 54.1 | 0.43 | 0.41 | 118 | 0 | 118 | 114 | -7 | 107 | 20 | 19 |
| *Ardisia* | *sanguinolenta* | Asa | ARDISANG | MYRS | 6.8 | 0.19 | 0.20 | 274 | -10 | 264 | 266 | -13 | 253 | 51 | 71 |
| *Baccaurea* | *tetrandra* | Bte | BACCTETR | EUPH | 40.8 | 0.24 | 0.24 | 106 | -3 | 103 | 92 | -4 | 88 | 17 | 19 |
| *Chisocheton* | *sarawakanus* | Csa | CHISSARA | MELI | 49.0 | 0.25 | 0.26 | 78 | -1 | 77 | 72 | -2 | 70 | 9 | 12 |
| *Cleistanthus* | *contractus* | Cco | CLEICONT | EUPH | 3.0 | 0.08 | 0.09 | 146 | -2 | 144 | 142 | -5 | 137 | 11 | 16 |
| *Dacryodes* | *rostrata* | Dro | DACRROST | BURS | 26.1 | 0.12 | 0.13 | 83 | 0 | 83 | 74 | -3 | 71 | 10 | 11 |
| *Dehaasia* | *gigantocarpa* | Dgi | DEHAGIGA | LAUR | 15.6 | 0.07 | 0.07 | 68 | -3 | 65 | 63 | 0 | 63 | 7 | 9 |
| *Dimorphocalyx* | *muricatus* | Dmu | DIMOMURI | EUPH | 9.8 | 0.34 | 0.33 | 407 | -11 | 396 | 370 | -6 | 364 | 55 | 63 |
| *Dysoxylum* | *cyrtobotryum* | Dcy | DYSOCYRT | MELI | 50.2 | 0.25 | 0.26 | 74 | -1 | 73 | 65 | -1 | 64 | 15 | 12 |
| *Fordia* | *splendidissima* | Fsp | FORDSPLE | LEGU | 4.0 | 0.16 | 0.18 | 251 | -7 | 244 | 250 | -5 | 245 | 32 | 65 |
| *Hydnocarpus* | *borneensis* | Hbo | HYDNBORN | FLAC | 10.6 | 0.04 | 0.04 | 58 | 0 | 58 | 59 | -4 | 55 | 6 | 5 |
| *Lithocarpus* | *gracilis* | Lgr | LITHGRAC | FAGA | 64.3 | 0.18 | 0.28 | 20 | 0 | 20 | 13 | 0 | 13 | 14 | 10 |
| *Lithocarpus* | *leptogyne* | Lle | LITHLEPT | FAGA | 56.5 | 0.17 | 0.23 | 27 | -1 | 26 | 31 | 0 | 31 | 9 | 9 |
| *Lithocarpus* | *nieuwenhuisii* | Lni | LITHNIEW | FAGA | 57.0 | 0.23 | 0.22 | 61 | -4 | 57 | 55 | -5 | 50 | 7 | 11 |
| *Litsea* | *caulocarpa* | Lca | LITSCAUL | LAUR | 16.4 | 0.15 | 0.15 | 140 | -4 | 136 | 110 | -1 | 109 | 53 | 77 |
| *Litsea* | *ochracea* | Loc | LITSOCHR | LAUR | 27.7 | 0.17 | 0.17 | 68 | -3 | 65 | 53 | -1 | 52 | 19 | 24 |
| *Lophopetalum* | *beccarianum* | Lbe | LOPHBECC | CELA | 18.3 | 0.13 | 0.15 | 149 | -1 | 148 | 167 | -7 | 160 | 15 | 18 |
| *Madhuca* | *korthalsii* | Mko | MADHKORT | SAPO | 32.1 | 0.70 | 0.69 | 266 | -8 | 258 | 265 | -5 | 260 | 20 | 38 |
| *Mallotus* | *penangensis* | Mpe | MALLPENA | EUPH | 19.4 | 0.12 | 0.13 | 109 | -1 | 108 | 122 | -4 | 118 | 14 | 21 |
| *Mallotus* | *stipularis* | Mst | MALLSTIP | EUPH | 27.9 | 0.10 | 0.09 | 65 | 0 | 65 | 53 | 0 | 53 | 8 | 17 |
| *Mallotus* | *wrayi* | Mwr | MALLWRAY | EUPH | 10.4 | 0.95 | 0.94 | 1031 | -11 | 1020 | 972 | -15 | 957 | 189 | 217 |
| *Maschalocorymbus* | *corymbosus* | Mco | MASCCORY | RUBI | 0.8 | 0.11 | 0.09 | 144 | -2 | 142 | 128 | -1 | 127 | 71 | 49 |
| *Neoscortechinia* | *philippinensis* | Nph | NEOSPHIL | EUPH | 70.2 | 0.34 | 0.33 | 44 | 0 | 44 | 36 | 0 | 36 | 7 | 9 |
| *Parashorea* | *malaanonan* | Pma | PARAMALA | DIPT | 65.1 | 0.71 | 0.90 | 51 | 0 | 51 | 37 | 0 | 37 | 14 | 19 |
| *Pentace* | *laxiflora* | Pla | PENTLAXI | TILI | 63.0 | 0.42 | 0.56 | 103 | -4 | 99 | 75 | -2 | 73 | 23 | 36 |
| *Polyalthia* | *cauliflora* | Pca | POLYCAUL | ANNO | 11.9 | 0.15 | 0.14 | 160 | -1 | 159 | 147 | -2 | 145 | 28 | 26 |

| | | | | | | | | | | | | | | | |
|---|---|---|---|---|---|---|---|---|---|---|---|---|---|---|---|
| *Polyalthia* | *rumphii* | Pru | POLYRUMP | ANNO | 20.0 | 0.07 | 0.08 | 78 | -1 | 77 | 74 | -1 | 73 | 6 | 13 |
| *Polyalthia* | *sumatrana* | Psu | POLYSUMA | ANNO | 44.0 | 0.34 | 0.34 | 113 | -1 | 112 | 114 | -3 | 111 | 12 | 13 |
| *Polyalthia* | *xanthopetala* | Pxa | POLYXANT | ANNO | 14.8 | 0.14 | 0.11 | 93 | -2 | 91 | 81 | -4 | 77 | 31 | 45 |
| *Reinwardtiodendron* | *humile* | Rhu | REINHUMI | MELI | 18.1 | 0.14 | 0.13 | 112 | -3 | 109 | 72 | -3 | 69 | 34 | 48 |
| *Shorea* | *fallax* | Sfa | SHORFALL | DIPT | 39.1 | 0.93 | 0.94 | 182 | -1 | 181 | 151 | -4 | 147 | 58 | 79 |
| *Shorea* | *johorensis* | Sjo | SHORJOHO | DIPT | 87.0 | 3.17 | 3.83 | 51 | 0 | 51 | 32 | -1 | 31 | 37 | 31 |
| *Shorea* | *parvifolia* | Spar | SHORPARF | DIPT | 86.6 | 2.32 | 2.49 | 51 | 0 | 51 | 40 | 0 | 40 | 32 | 22 |
| *Shorea* | *pauciflora* | Spau | SHORPAUC | DIPT | 65.8 | 1.17 | 1.09 | 42 | 0 | 42 | 42 | 0 | 42 | 18 | 17 |
| *Shorea* | *pilosa* | Spi | SHORPILO | DIPT | 69.8 | 0.97 | 1.00 | 59 | 0 | 59 | 52 | 0 | 52 | 13 | 21 |
| *Syzygium* | *elopurae* | Sel | SYZYELOP | MYRT | 23.2 | 0.10 | 0.09 | 66 | -1 | 65 | 53 | -2 | 51 | 15 | 18 |
| *Syzygium* | *tawaense* | Sta | SYZYTAWA | MYRT | 57.7 | 0.44 | 0.50 | 35 | 0 | 35 | 32 | -3 | 29 | 8 | 6 |
| *Xanthophyllum* | *vitellinum* | Xvi | XANTVITE | POLY | 28.6 | 0.10 | 0.10 | 72 | -2 | 70 | 65 | -1 | 64 | 5 | 7 |
| *Barringtonia* | *lanceolata* | Bla | BARRLANC | LECY | 60.4 | 0.35 | 0.35 | 81 | -1 | 80 | 74 | -2 | 72 | 1 | 10 |
| *Dipterocarpus* | *kerrii* | Dke | DIPTKERR | DIPT | 56.0 | 0.24 | 0.28 | 24 | 0 | 24 | 22 | 0 | 22 | 1 | 5 |
| *Drypetes* | *longifolia* | Dlo | DRYPLONG | EUPH | 55.3 | 0.14 | 0.12 | 28 | 0 | 28 | 25 | 0 | 25 | 3 | 7 |
| *Gonystylus* | *keithii* | Gke | GONYKEIT | THYM | 42.2 | 0.24 | 0.25 | 61 | -1 | 60 | 60 | -1 | 59 | 0 | 9 |
| *Knema* | *latericia* | Kla | KNEMLATE | MYRI | 5.1 | 0.04 | 0.05 | 83 | 0 | 83 | 95 | -3 | 92 | 2 | 7 |
| *Microcos* | *reticulata* | Mre | MICRRETI | TILI | 60.6 | 0.12 | 0.11 | 23 | -1 | 22 | 19 | 0 | 19 | 4 | 7 |
| *Ochanostachys* | *amentacea* | Oam | OCHAAMEN | OLAC | 59.5 | 0.24 | 0.22 | 25 | 0 | 25 | 19 | 0 | 19 | 3 | 7 |
| *Scorodocarpus* | *borneensis* | Sbo | SCORBORN | OLAC | 84.8 | 0.92 | 0.89 | 27 | -1 | 26 | 23 | -1 | 22 | 5 | 7 |
| *Syzygium* | *lineata* | Sli | SYZYLINE | MYRT | 63.0 | 0.29 | 0.33 | 22 | 0 | 22 | 24 | -1 | 23 | 3 | 3 |
| *Vatica* | *dulitensis* | Vcu | VATIDULI | DIPT | 45.1 | 0.37 | 0.36 | 53 | -2 | 51 | 52 | -2 | 50 | 4 | 6 |

**Appendix 1: Table S2**. Number of species (out of 38) for which regression models are > 7 $\Delta AIC_c$ worse/better than, or ≤ 7 $\Delta AIC_c$ indifferent from, the *ba* + ALL model with linear distance decay as a reference (0/0/38). The dependent variable was (a) absolute growth rate, or (b) survival (0/1), in period 1 ($P_1$, 1986 – 1996) or period 2 ($P_2$, 1996 – 2007). Numbers of species are given for models with size only (*ba*), then with one neighbour term added (*ba* + ALL), or two neighbourhood terms added (*ba* + CON + HET), as non-spatial models; and for the two neighbour terms with one of each of the eight different forms of crown extension (with [no], linear [lin] or squared [squ] distance decay), as spatial models. Models with two neighbour terms and spatial extension were allowed to have different Δd's for their CON and HET terms. Zones of influence of individual trees were filled either with equal numbers of points per unit crown area (*ppsqmca*: 'equal') or such that large trees had more points per unit crown area than smaller trees (*ppsqmca*: 'larger > smaller'), as a function of stem *gbh* (see Methods). Covered modules (i.e. modules with modules of bigger trees within a given distance) were either removed (pruned) or relocated, and the trees position (pos) taken as being at either the original stem coordinates or the centre of gravity of the uncovered points.

| **(a) Growth** | | | | Period 1 | | | Period 2 | | |
|---|---|---|---|---|---|---|---|---|---|
| | | | | Decay | | | Decay | | |
| Model | *ppsqmca* | covered | pos. | no | lin | squ | no | lin | squ |
| ba | – | – | – | 10/ 0/28 | – | – | 5/ 0/33 | – | – |
| ba + ALL | – | – | – | 0/ 0/38 | 0/ 0/38 | 0/ 0/38 | 0/ 0/38 | 0/ 0/38 | 0/ 0/38 |
| ba + CON + HET | – | – | – | 0/ 6/32 | 0/ 4/34 | 1/ 4/33 | 0/ 3/35 | 0/ 4/34 | 0/ 1/37 |
| | equal | removed | stem | 0/12/26 | 0/13/25 | 2/ 7/29 | 0/11/27 | 2/ 9/27 | 4/ 6/28 |
| | " | " | crown | 0/14/24 | 0/14/24 | 2/11/25 | 0/11/27 | 0/11/27 | 1/ 6/31 |
| | " | relocated | stem | 0/14/24 | 0/11/27 | 1/ 9/28 | 1/13/24 | 2/14/22 | 3/ 5/30 |
| | " | " | crown | 0/14/24 | 0/11/27 | 1/10/27 | 1/14/23 | 1/14/23 | 2/ 7/29 |
| | larger > smaller | removed | stem | 0/10/28 | 2/ 9/27 | 4/ 8/26 | 3/ 7/28 | 3/ 7/28 | 4/ 6/28 |
| | " | " | crown | 0/ 9/29 | 2/ 8/28 | 4/ 8/26 | 2/ 7/29 | 2/ 8/28 | 3/ 7/28 |
| | " | relocated | stem | 0/11/27 | 1/ 9/28 | 2/ 8/28 | 2/ 8/28 | 2/ 8/28 | 4/ 7/27 |
| | " | " | crown | 0/12/26 | 0/10/28 | 0/ 9/29 | 2/ 7/29 | 2/ 7/29 | 3/ 6/29 |

| **(b) Survival** | | | | Period 1 | | | Period 2 | | |
|---|---|---|---|---|---|---|---|---|---|
| | | | | Decay | | | Decay | | |
| Model | *ppsqmca* | covered | pos. | no | lin | squ | no | lin | squ |
| ba | – | – | – | 2/ 0/36 | – | – | 2/ 0/36 | – | – |
| ba + ALL | – | – | – | 0/ 0/38 | 0/ 0/38 | 0/ 0/38 | 0/ 0/38 | 0/ 0/38 | 0/ 0/38 |
| ba + CON + HET | – | – | – | 0/ 2/36 | 0/ 0/38 | 0/ 0/38 | 0/ 1/37 | 0/ 1/37 | 1/ 1/36 |
| | equal | removed | stem | 0/ 5/33 | 0/ 2/36 | 4/ 2/32 | 0/ 9/29 | 0/ 6/32 | 2/ 5/31 |
| | " | " | crown | 0/ 5/33 | 0/ 4/34 | 3/ 3/32 | 0/ 9/29 | 0/ 5/33 | 2/ 4/32 |
| | " | relocated | stem | 1/ 4/33 | 1/ 2/35 | 2/ 2/34 | 0/11/27 | 0/ 7/31 | 1/ 4/33 |
| | " | " | crown | 1/ 4/33 | 1/ 2/35 | 2/ 1/35 | 0/ 9/29 | 0/ 7/31 | 1/ 2/35 |
| | larger > smaller | removed | stem | 0/ 6/32 | 1/ 2/35 | 2/ 1/35 | 0/ 5/33 | 1/ 4/33 | 1/ 3/34 |
| | " | " | crown | 0/ 6/32 | 0/ 5/33 | 1/ 3/34 | 0/ 4/34 | 0/ 3/35 | 1/ 3/34 |
| | " | relocated | stem | 0/ 4/34 | 0/ 2/36 | 1/ 1/36 | 0/ 5/33 | 0/ 3/35 | 1/ 3/34 |
| | " | " | crown | 0/ 5/33 | 0/ 3/35 | 1/ 2/35 | 0/ 5/33 | 0/ 2/36 | 1/ 2/35 |

Table S3. As Table S2 in the main text but with ‘ba + CON + HET’ as the reference model instead of ‘ba + ALL’. Otherwise the table parameters are the same.

| **(a) Growth** | | | | Period 1 | | | Period 2 | | |
|---|---|---|---|---|---|---|---|---|---|
| | | | | Decay | | | Decay | | |
| Model | *ppsqmca* | covered | pos. | no | lin | squ | no | lin | squ |
| ba | – | – | – | 14/ 0/24 | – | – | 9/ 0/29 | – | – |
| ba + ALL | – | – | – | 5/ 0/33 | 4/ 0/34 | 7/ 0/31 | 1/ 0/37 | 4/ 0/34 | 4/ 0/34 |
| ba + CON + HET | – | – | – | 0/ 0/38 | 0/ 0/38 | 0/ 0/38 | 0/ 0/38 | 0/ 0/38 | 0/ 0/38 |
| | equal | removed | stem | 0/10/28 | 0/10/28 | 5/ 4/29 | 1/10/27 | 0/10/28 | 4/ 4/30 |
| | “ | “ | crown | 0/11/27 | 0/ 9/29 | 5/ 5/28 | 0/12/26 | 0/10/28 | 3/ 5/30 |
| | “ | relocated | stem | 1/12/25 | 0/ 9/29 | 5/ 6/27 | 1/13/24 | 1/11/26 | 3/ 4/31 |
| | “ | “ | crown | 1/12/25 | 0/10/28 | 5/ 5/28 | 1/13/24 | 1/ 9/28 | 3/ 5/30 |
| | larger > smaller | removed | stem | 0/ 3/35 | 1/ 3/34 | 7/ 3/28 | 2/ 5/31 | 3/ 5/30 | 6/ 4/28 |
| | “ | “ | crown | 0/ 4/34 | 2/ 4/32 | 6/ 3/29 | 2/ 5/31 | 3/ 6/29 | 5/ 6/27 |
| | “ | relocated | stem | 0/ 5/33 | 0/ 3/35 | 4/ 2/32 | 2/ 5/31 | 3/ 5/30 | 5/ 5/28 |
| | “ | “ | crown | 0/ 4/34 | 1/ 4/33 | 4/ 1/33 | 2/ 6/30 | 3/ 4/31 | 4/ 5/29 |

| **(b) Survival** | | | | Period 1 | | | Period 2 | | |
|---|---|---|---|---|---|---|---|---|---|
| | | | | Decay | | | Decay | | |
| Model | *ppsqmca* | covered | pos. | no | lin | squ | no | lin | squ |
| ba | – | – | – | 4/ 0/34 | – | – | 5/ 0/33 | – | – |
| ba + ALL | – | – | – | 0/ 0/38 | 0/ 0/38 | 1/ 0/37 | 2/ 0/36 | 1/ 0/37 | 2/ 0/36 |
| ba + CON + HET | – | – | – | 0/ 0/38 | 0/ 0/38 | 0/ 0/38 | 0/ 0/38 | 0/ 0/38 | 0/ 0/38 |
| | equal | removed | stem | 0/ 1/37 | 1/ 0/37 | 4/ 1/33 | 0/ 6/32 | 0/ 1/37 | 2/ 1/35 |
| | " | " | crown | 0/ 1/37 | 1/ 0/37 | 2/ 2/34 | 0/ 7/31 | 0/ 3/35 | 1/ 3/34 |
| | " | relocated | stem | 1/ 1/36 | 2/ 0/36 | 3/ 1/34 | 0/ 7/31 | 0/ 4/34 | 0/ 1/37 |
| | " | " | crown | 1/ 1/36 | 2/ 0/36 | 3/ 1/34 | 0/ 7/31 | 0/ 3/35 | 0/ 1/37 |
| | larger > smaller | removed | stem | 1/ 2/35 | 1/ 1/36 | 1/ 0/37 | 0/ 0/38 | 0/ 0/38 | 2/ 1/35 |
| | " | " | crown | 1/ 2/35 | 1/ 1/36 | 1/ 1/36 | 0/ 1/37 | 1/ 1/36 | 1/ 1/36 |
| | " | relocated | stem | 0/ 2/36 | 1/ 2/35 | 1/ 1/36 | 0/ 0/38 | 0/ 0/38 | 1/ 1/36 |
| | " | " | crown | 0/ 2/36 | 1/ 2/35 | 1/ 1/36 | 0/ 1/37 | 0/ 1/37 | 1/ 1/36 |

Table S4. Numbers of species reaching increasing levels of (a) variance accounted for, and significance of (b) CON and (c) HET coefficients, averaged across all fitted models within $2\Delta AIC_c$ of the best fitting one; for growth and survival response models in $P_1$ and $P_2$. 'Non-spat' is the non-spatial model, and 'relocate' and 'remove' indicate the 'larsm/reloc/crown' and 'larsm/remov/crown' spatial models (linear decay).

| [a] mean adj.$R^2$ (%) | | | Number of species | | | | | |
|---|---|---|---|---|---|---|---|---|
| | | | <2 | < 5 | < 10 | < 20 | < 50 | ≥ 50 |
| Growth | P1 | non-spat | 0 | 3 | 5 | 8 | 29 | 9 |
| | | relocate | 1 | 3 | 5 | 9 | 27 | 11 |
| | | remove | 1 | 3 | 5 | 9 | 27 | 11 |
| | P2 | non-spat | 0 | 1 | 5 | 11 | 27 | 11 |
| | | relocate | 0 | 1 | 3 | 12 | 26 | 12 |
| | | remove | 1 | 1 | 4 | 12 | 25 | 13 |
| Survival | P1 | non-spat | 8 | 21 | 28 | 35 | 38 | 0 |
| | | relocate | 5 | 20 | 25 | 33 | 38 | 0 |
| | | remove | 10 | 19 | 26 | 35 | 38 | 0 |
| | P2 | non-spat | 5 | 19 | 29 | 33 | 37 | 1 |
| | | relocate | 3 | 21 | 29 | 33 | 37 | 1 |
| | | remove | 3 | 22 | 29 | 35 | 36 | 2 |

| [b] mean CON_P-value | | | Number of species | | | | | |
|---|---|---|---|---|---|---|---|---|
| | | | < 0.001 | < 0.01 | < 0.05 | < 0.10 | < 0.25 | ≥ 25 |
| Growth | P1 | non-spat | 4 | 8 | 17 | 18 | 24 | 14 |
| | | relocate | 10 | 16 | 22 | 25 | 29 | 9 |
| | | remove | 7 | 16 | 21 | 23 | 31 | 7 |
| | P2 | non-spat | 3 | 5 | 9 | 13 | 22 | 16 |
| | | relocate | 5 | 9 | 16 | 20 | 26 | 12 |
| | | remove | 5 | 9 | 15 | 19 | 24 | 14 |
| Survival | P1 | non-spat | 0 | 1 | 7 | 13 | 19 | 19 |
| | | relocate | 0 | 2 | 11 | 17 | 28 | 10 |
| | | remove | 0 | 2 | 9 | 17 | 25 | 13 |
| | P2 | non-spat | 0 | 2 | 6 | 11 | 17 | 21 |
| | | relocate | 1 | 2 | 8 | 12 | 21 | 17 |
| | | remove | 1 | 2 | 7 | 10 | 18 | 20 |

| [c] mean HET_P-value | | | Number of species | | | | | |
|---|---|---|---|---|---|---|---|---|
| | | | < 0.001 | < 0.01 | < 0.05 | < 0.10 | < 0.25 | ≥ 25 |
| Growth | P1 | non-spat | 4 | 16 | 22 | 26 | 30 | 8 |
| | | relocate | 5 | 9 | 20 | 24 | 25 | 13 |
| | | remove | 5 | 10 | 15 | 24 | 30 | 8 |
| | P2 | non-spat | 5 | 8 | 17 | 23 | 30 | 8 |
| | | relocate | 4 | 5 | 16 | 23 | 28 | 10 |
| | | remove | 3 | 7 | 17 | 23 | 29 | 9 |
| Survival | P1 | non-spat | 1 | 2 | 6 | 11 | 20 | 18 |
| | | relocate | 1 | 3 | 7 | 11 | 17 | 21 |
| | | remove | 1 | 2 | 4 | 11 | 19 | 19 |
| | P2 | non-spat | 0 | 1 | 8 | 14 | 21 | 17 |
| | | relocate | 0 | 1 | 6 | 9 | 24 | 14 |
| | | remove | 0 | 1 | 6 | 14 | 24 | 14 |

Table S5. Ranges of the correlations coefficients, r, found pair-wise among the eight spatial models, with their corresponding *t*-values, for the eight combinations of conspecific vs heterospecific effect size (CON, HET), growth vs survival variables (G, S), and periods 1 and 2 ($P_1$, $P_2$). The critical *t*-value for a one-sided test at $P = 0.0001$ is 4.140 (df = 36).

| | | | *r* | *t* |
|---|---|---|---|---|
| CON | G | P1 | 0.9323 – 0.9956 | 15.471 – 63.593 |
| | | P2 | 0.9625 – 0.9973 | 21.303 – 80.880 |
| CON | S | P1 | 0.6924 – 0.9985 | 5.757 – 108.17 |
| | | P2 | 0.6707 – 0.9915 | 5.426 – 45.732 |
| HET | G | P1 | 0.8594 – 0.9919 | 10.085 – 46.892 |
| | | P2 | 0.7022 – 0.9647 | 5.918 – 21.972 |
| HET | S | P1 | 0.6779 – 0.9899 | 5.533 – 41.682 |
| | | P2 | 0.3244 – 0.9998 | 2.058 – 330.47 |

Table S6. Correlations (Pearson-*r*) between combinations of conspecific vs heterospecific effect size (CON, HET), growth vs survival variables (G, S), and periods 1 and 2 ($P_1$, $P_2$), for the "larsm/**reloc**/crown" (upper triangle) and "larsm/**remov**/crown" (lower triangle) spatial models. T-tests for *r*-values were two-tailed. Coefficients in bold face are significant at $P \leq 0.05$.

| | CON.G.P1 | CON.G.P2 | CON.S.P1 | CON.S.P2 | HET.G.P1 | HET.G.P2 | HET.S.P1 | HET.S.P2 |
|---|---|---|---|---|---|---|---|---|
| CON.G.P1 | | *** **0.521** | 0.133 | 0.027 | *****0.548** | ° 0.300 | 0.047 | −0.127 |
| CON.G.P2 | *****0.545** | | −0.234 | 0.007 | ***0.363** | ****0.440** | ***−0.327** | −0.126 |
| CON.S.P1 | −0.004 | −0.233 | | 0.114 | 0.089 | −0.090 | ***0.332** | -0.108 |
| CON.S.P2 | 0.021 | −0.032 | −0.156 | | −0.226 | 0.308 | −0.099 | −0.047 |
| HET.G.P1 | ** **0.502** | * **0.375** | −0.083 | −0.021 | | −0.008 | 0.077 | °0.318 |
| HET.G.P2 | 0.295 | * **0.399** | −0.094 | −0.111 | 0.019 | | −0.122 | ****−0.433** |
| HET.S.P1 | 0.035 | −0.300 | °0.320 | −0.209 | 0.072 | −0.042 | | −0.026 |
| HET.S.P2 | −0.118 | −0.089 | −0.116 | * **0.371** | 0.196 | *****−0.665** | −0.066 | |

***, $P \leq 0.001$; **, $P \leq 0.01$; *, $P \leq 0.05$; °, $P \leq 0.10$.

Appendix 1: Fig. S1

Frequency histograms of (a) correlations ('cor_'), and (b) slopes ('slope_'), of the relationships between fitted CON ('con') and HET ('het') neighborhood regression coefficients versus their corresponding best fitting radii, when modelling either growth ('g') or survival ('s') focus tree responses, in periods $P_1$ ('P1') or $P_2$ ('P2'), for the 38 selected species. In some panels the total frequency is < 38 when any species' regression had ≤ 2 fitted models within the band 2ΔAICc.

(a) correlations

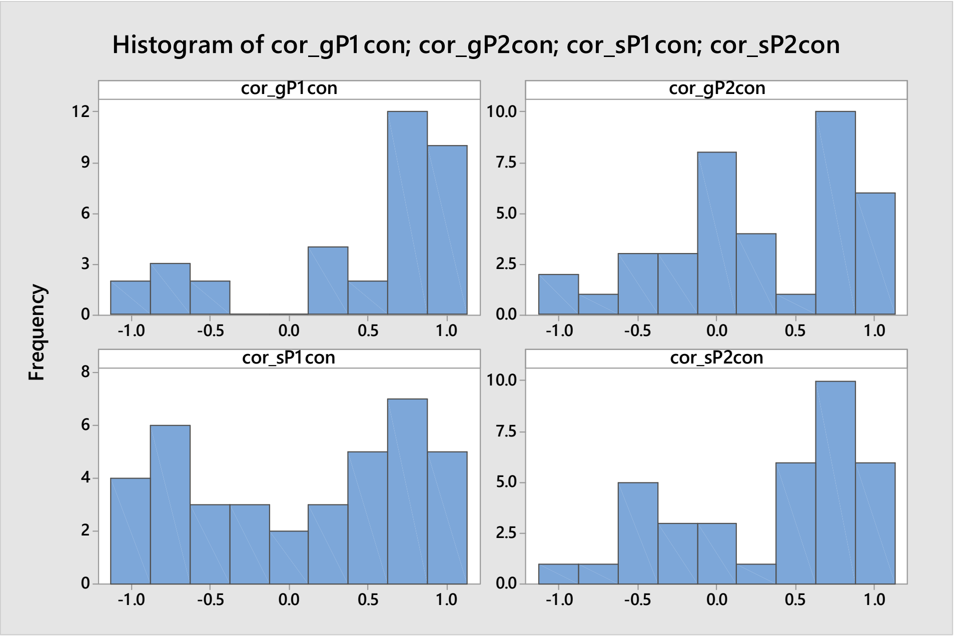


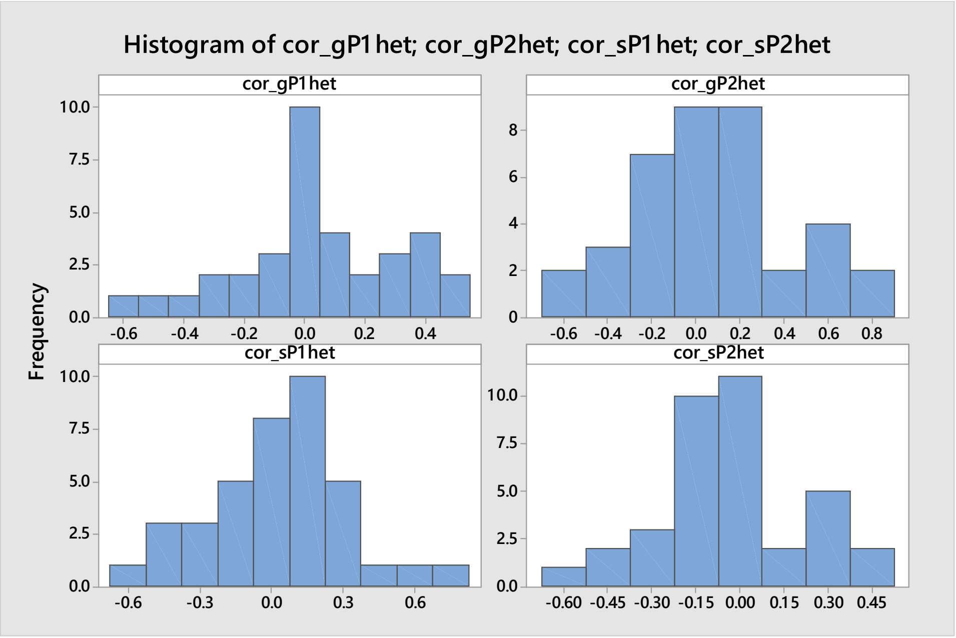

(b) slopes

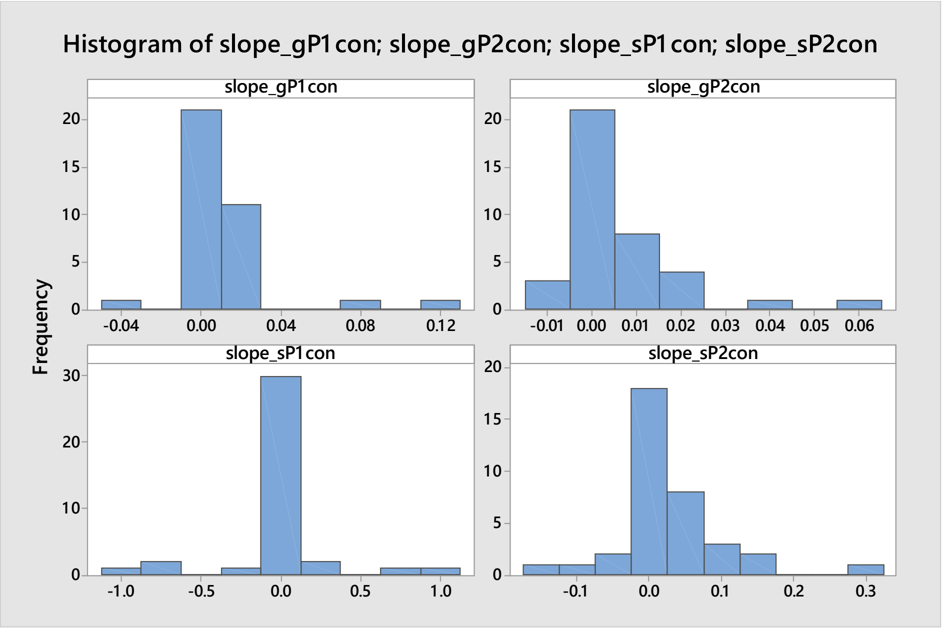


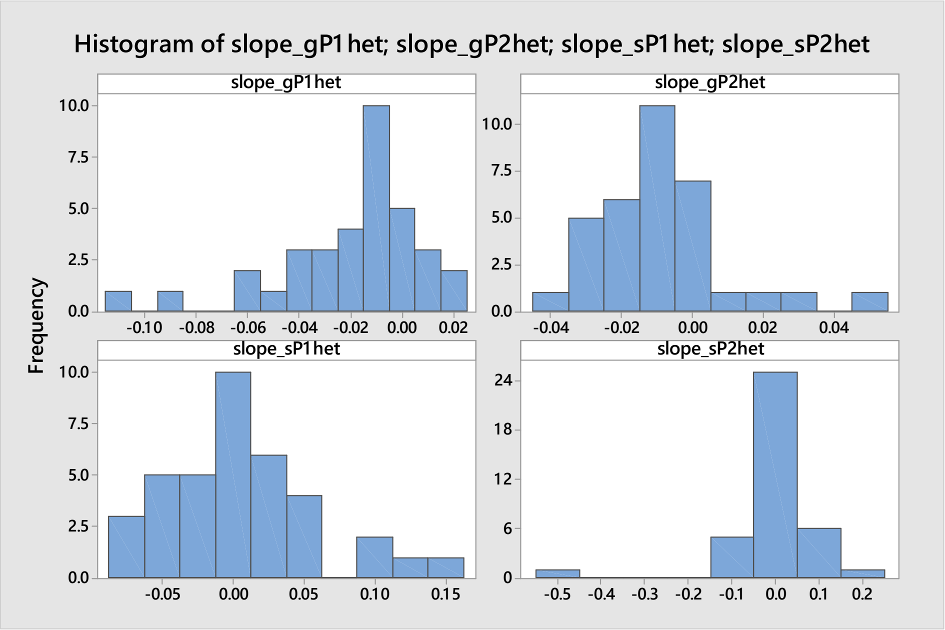


Note: In the lower left panel of the upper set of (b), i.e. for ‘slope_sP1con’, the five cases mentioned in the text with very outlying slopes were *Dimorphocalyx muricatus*, *Neoscortechinia philippinensis*, and *Parashorea malaanonan* (negative), and *Polyalthia rumphii* and *Ardisia sanguinolenta* (positive).

Appendix 1: Fig. S2.

Frequency histograms of mean $\Delta d_{CON}$ and $\Delta d_{HET}$-values corresponding to the best fitting 'CON' and 'HET' neighborhood regression coefficients when modelling either growth ('g') or survival ('s') focus tree responses, in periods $P_1$ ('P1') or $P_2$ ('P2'), for the 38 selected species, using the spatial models 'larsm/crown/remov' and (b) 'larsm/crown/reloc'. The means were those over all model fits per species within the band 2ΔAICc.

(a) Point adjustment by removal

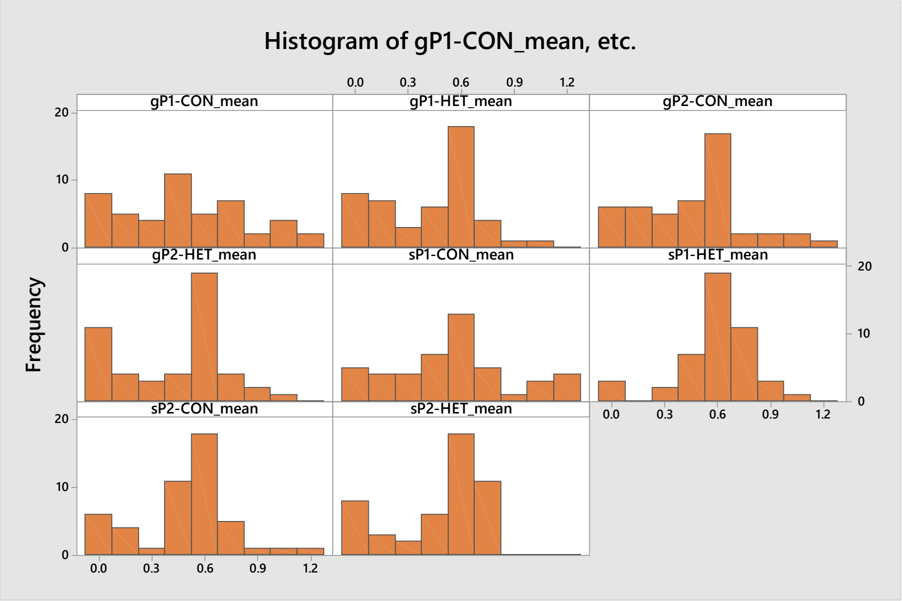


(b) Point adjustment by relocation

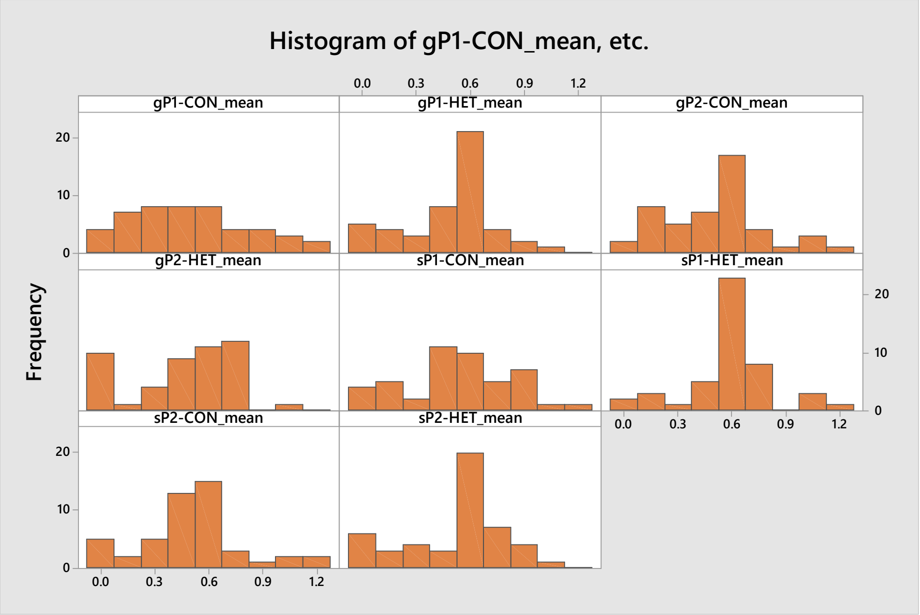

Appendix 1: Fig. S3. Relationships between *differences* in conspecific (CON) effect sizes on growth rates between periods ($P_2 - P_1$) and (a) CON and (b) HET effects on survival in period 1 for the 38 species, from non-spatial neighbourhood models with size (*ba*) and two neighbour terms (HET and CON). These figures complement the spatial models in Figs. 2 and 3 in main text, using the same labelling and color-coding.

(a)

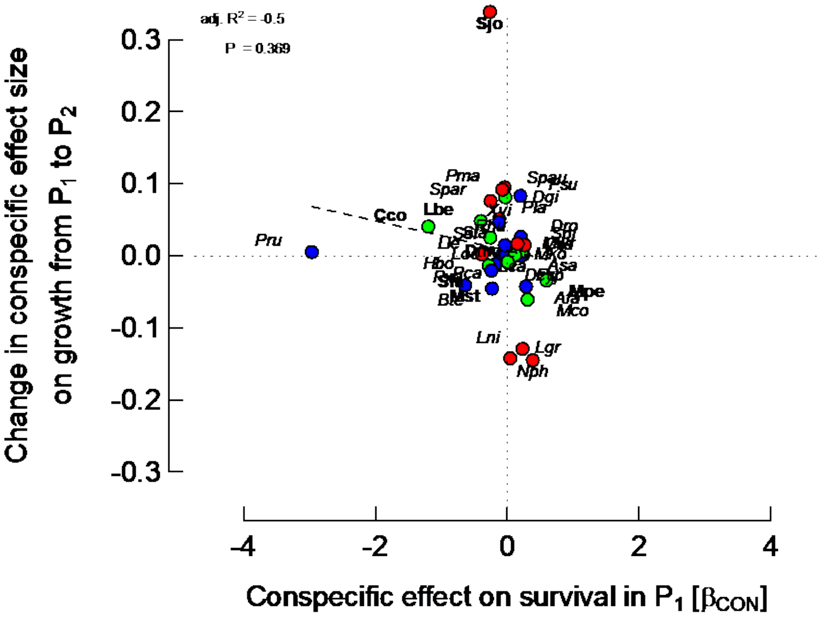


(b)

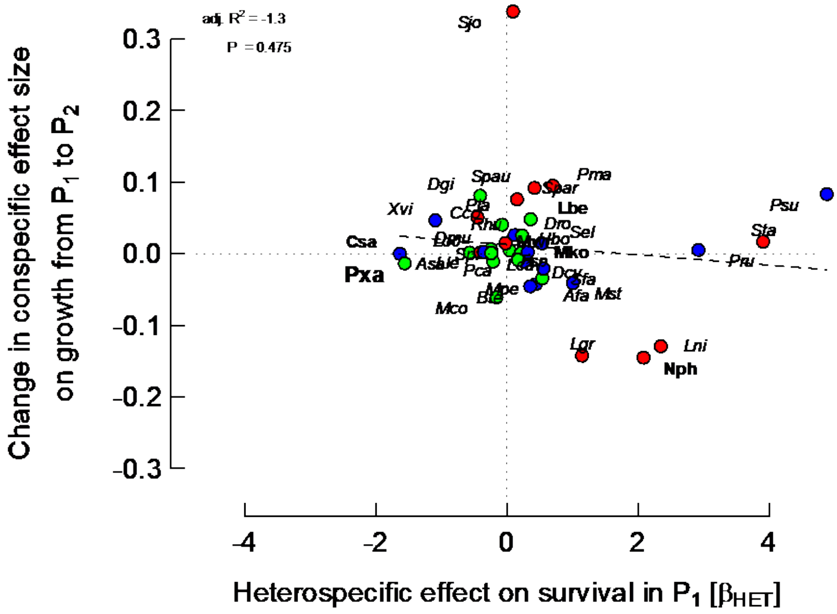

Appendix 1: Fig. S4. Relationships between *differences* in heterospecific (HET) effect sizes on growth rates between periods ($P_2 - P_1$) and (a) CON and (b) HET effects on survival in period 1 for the 38 species, from non-spatial neighbourhood models with size (*ba*) and two neighbour terms (HET and CON). These figures complement the spatial models in Figs. 4 and 5 in main text, using the same labelling and color-coding.

(a)

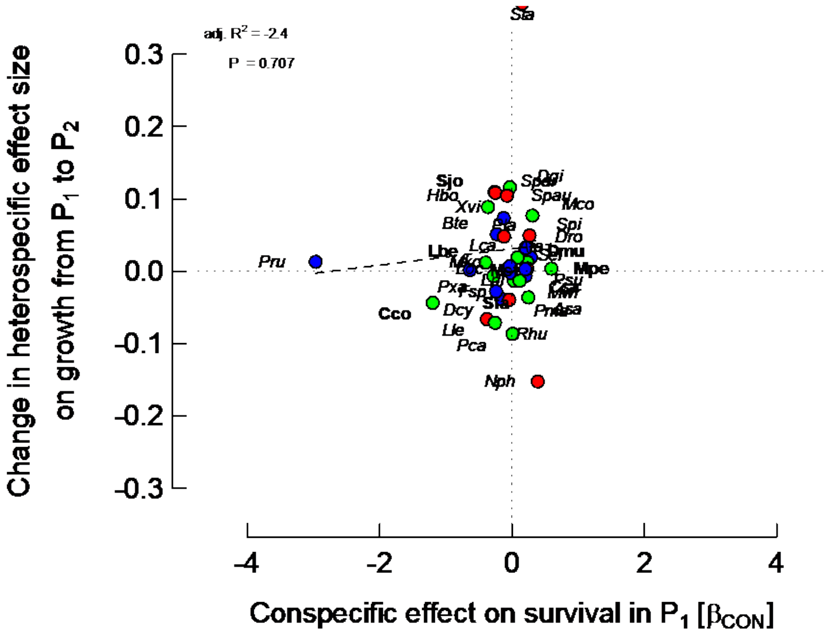


(b)

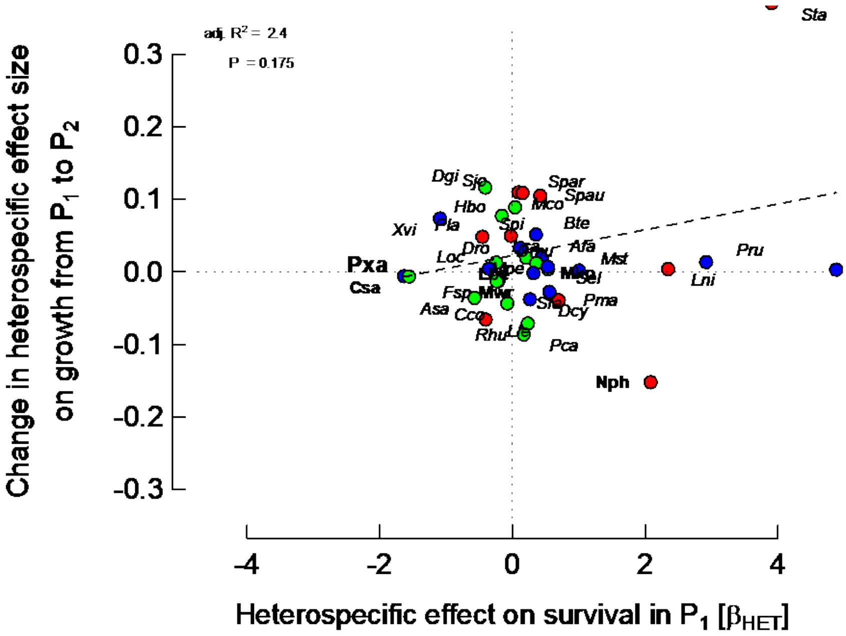

Appendix 1: Fig. S5. Relationships between conspecific (CON) effect sizes on growth rates in periods (a) 1, and (b) 2, and CON effects on survival in period 1 for the 38 species, from the ‘larsm/reloc/crown’ spatial neighbourhood model with size (*ba*) and two neighbour terms (HET and CON). These figures complement the spatial models in Fig. 2 (b) in main text, using the same labelling and color-coding.

(a)

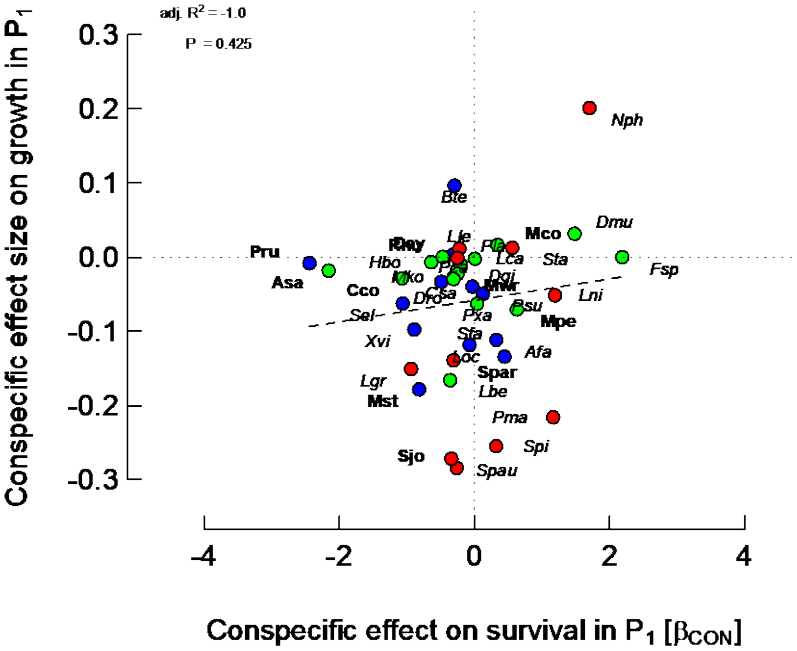


(b)

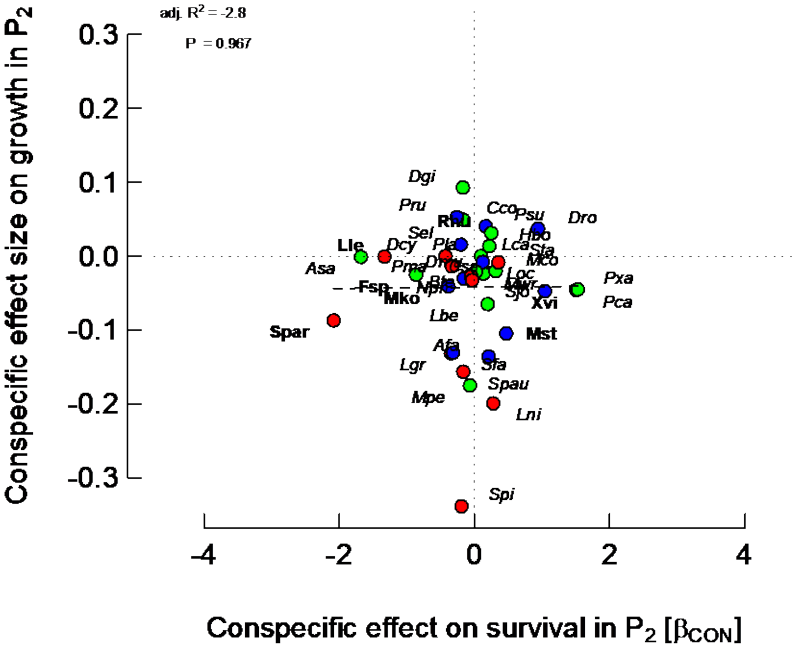

Appendix 1: Fig. S6. Relationships between (a) conspecific (CON), and (b) heterospecific (HET) effect sizes on survival in period 1, for the 38 species, from the spatial ('larsm/crown/reloc') versus non-spatial models with size (*ba*) and two neighbour terms (HET and CON). Important outliers: (a) Pru, and (b) Pru, Sta, Psu (see main text).

(a)

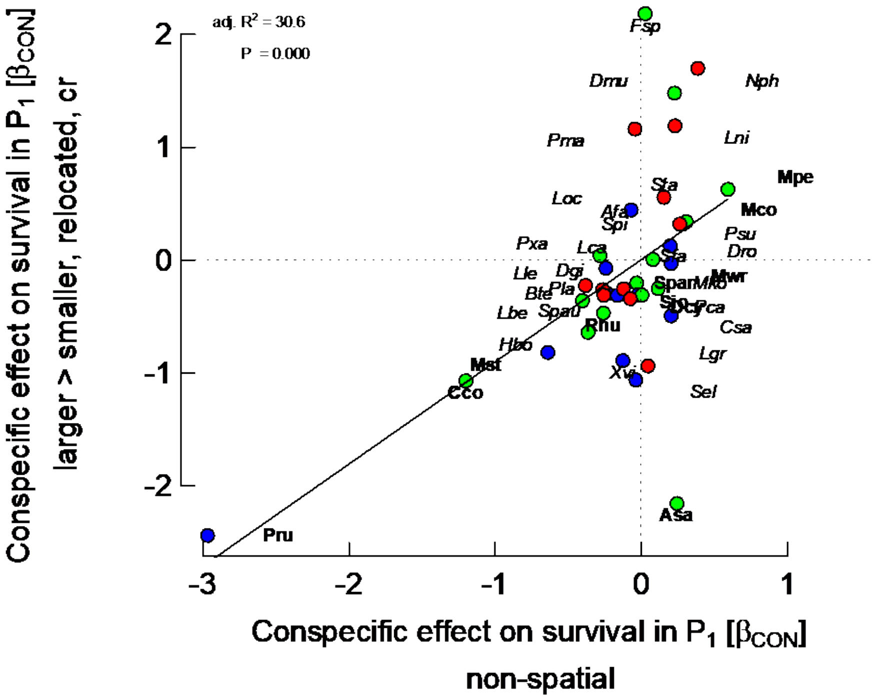


(b)

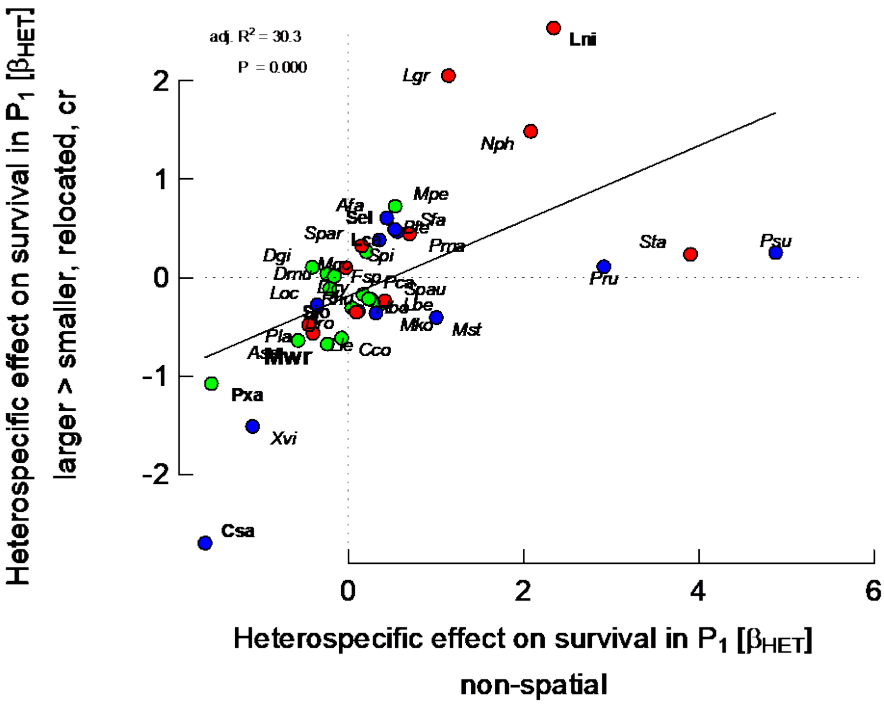
