## Supplementary material for "Including tree spatial extension in the evaluation of neighbourhood competition effects in Bornean rain forest": nncrexm appendices: App2_programs.pdf (from another version of this preprint: version 2, 2020-08-04)

**Appendix:** explaining the algorithms used to extend trees spatially, taking potential plasticity into account and evaluate their neighbourhoods.

The **R-Scripts** in folder **GitHub** are intended to demonstrate the principles. Their functionality is exactly as those used in the analyses presented in the main text. However, they were simplified as far as possible to illustrate their functionality. The data in **FULL_test.txt** are constructed to facilitate understanding of the functionality of the algorithms and are not meant to imply any biological meaning. The scripts are available at https://github.com/stollp/nncrexm, under a **Creative Commons** (CC) **license** without any warranty or support. However, the author may be contacted for informal exchange of ideas.

**FULL_test.txt**

| Plot | Tag | X | Y | F_Code07 | Code07 | GBH.86 | GBH.96 | GBH.01 | GBH.07 | col |
|---|---|---|---|---|---|---|---|---|---|---|
| MP1 | 1 | 5.0 | 5.0 | EUPH | MALLWRAY | 50 | 13.4 | 0.0 | NA | "red" |
| MP1 | 2 | 15.0 | 5.0 | MYRT | SYZYELOP | 175 | 112.8 | 113.2 | 0 | "green3" |
| MP1 | 3 | 5.0 | 15.0 | LAUR | BEILGLAU | 200 | 18.4 | 0.0 | NA | "blue" |
| MP1 | 4 | 12.1 | 12.1 | DIPT | SHORJOHO | 300 | 18.4 | 0.0 | NA | "orange" |
| MP1 | 5 | 12.1 | 10.0 | EUPH | MALLWRAY | 50 | 13.4 | 0.0 | NA | "red" |

GBH.YEAR: girth at breast height. GBH.96, .01, and .07 have no meaning and are only kept for consistency with the original data file. For example, the condition GBH.96 > 0 was used in some of the scripts to take only living trees as neighbours.

**Fig. S1**. From abstract, mathematical to impressionistic representation of trees with crown plasticity in neighbourhood models. *This figure is similar to Fig. 1 in the main text but the point density is fixed for showing how the algorithms work.* (a) Classical neighbourhood models represent trees as points without any spatial extension. Taking the red tree at X/Y = 5/5 m as focal tree, it has no neighbours at 5 m neighbourhood radius, four (1 con and 3 het) within 15 m and, depending on the exact definition of neighbours (i.e. < 10 or ≤ 10 m), one or four neighbours at 10 m neighbourhood radius. (b) Impressionistic representation of tree crowns as circles filled with 100 points (to facilitate counting) each. The crown radii (*cr*, in m) were allometrically related to girth at breast height (*gbh*, in cm). Girth of the smallest trees (red at 5/5 and 12.1/10) are 50 cm, those of their neighbours in increasing girth order 175 cm (green at 15/5), 200 cm (blue at 5/15) and 300 cm (orange at 12.1/12.1). These girths correspond to crown radii of 2.7, 5.9, 6.4 and 8.1 m, respectively (all-species regression, Table 1). Coordinates of points filling the crowns (or ZOI) were random. For the analyses presented in the main text, crowns were either filled with as many points as the trees basal area at breast height (*ba*, in $cm^2$) or 10 points per unit crown area. Filling was realised by ‘blowing up’ the original data file with as many rows per individual tree as points within the crown and initially flagging each point as ‘uncovered’. All three bigger heterospecific neighbours of the focal tree at 5/5 now have at least parts of their crowns already within the focal trees 5 m neighbourhood (4 blue, 1 green and 9 yellow = 14). The total number of conspecific points with in each 15 m neighbourhood of the red focals is 100 points. The total number of heterospecific points within each 20 m neighbourhoods of the red focals should be 300 as demonstrated in the following Output from R-Script 2_NN.R using function eval_NN.R.

```
 r Tag    x  y nos n_c n_C n_h n_H a_c o_C o_h o_H  l_c l_C l_h  l_H
 1   1  5.0  5   0   0   0   0   0   0   0   0   0  0.0   0   0  0.0
 5   1  5.0  5   3   0   0   0  14   0   0   0  14  0.0   0   0  3.7
10   1  5.0  5   4  82   0   0 139  82   0   0 139 10.2   0   0 20.5
15   1  5.0  5   4 100   0   0 270 100   0   0 270 11.9   0   0 31.2
20   1  5.0  5   4 100   0   0 300 100   0   0 300 11.9   0   0 33.1
 1   5 12.1 10   1   0   0   0   1   0   0   0   1  0.0   0   0  1.2
 5   5 12.1 10   3   0   0   0  70   0   0   0  70  0.0   0   0 23.7
10   5 12.1 10   4  78   0   0 247  78   0   0 247  9.4   0   0 48.7
15   5 12.1 10   4 100   0   0 300 100   0   0 300 11.5   0   0 53.3
20   5 12.1 10   4 100   0   0 300 100   0   0 300 11.5   0   0 53.3
```

r: neighbourhood radius, nos: number of species, n_ number of points of smaller or equally sized conspecifics (_c), heterospecifics (_h) and bigger con (_C) or heterospecific (_H) neighbours. o_ sum of points (‘basal area’) of neighbours (the number of points), l_ sum of points weighed by distance.

Taking plasticity into account

Points within ZOI’s may have points of bigger neighbours within their immediate neighbourhood as a function of some distance ($\Delta$d). Therefore, each points local neighbourhood is evaluated within $\Delta$d (0, 0.2, …, 1.2). $\Delta$d = 0 would allow complete overlap (as shown in b), whereas larger values of $\Delta$d flag individual points (open symbols in d)) as being ‘shaded’ if they have points of bigger neighbours within $\Delta$d. If a tree is completely covered as the red focal at 12.1/10, it is still used as focal, but from the perspective of the red focal at 5/5 it ‘disappears’ completely only as neighbour.

Two possibilities are used to handle these shaded points. First, (in c) the points are simply completely removed (*pruned*). Second, (in d), in an attempt to mimic plasticity, shaded points are relocated to unshaded parts of the crown using two-dimensional contour functions (function `kde2d` in MASS) to find the outline of these points. These contours were allowed to be larger as the original crowns by evaluating density contour lines using contourLines(Den, levels = c(0, 0.001, 0.01, 0.05, .1, 0.25, 0.5, 1)) and of these taking the lowest density levels at the outer edges of the crowns. The total number of points is reduced in c) but remains unaltered in d). Note that removal or relocation is only relevant for trees in their ‘role’ as neighbours. As focals, trees have a position and the neighbourhood is evaluated around this position given by the trees coordinate no matter whether or not it is completely covered. That is to say that figures 1c) and 1d) with $\Delta$d = 0, would look exactly as figure 1b), because not a single point would have a point of a bigger neighbour exactly on top of itself. The neighbourhood of each tree is evaluated by counting the number of points of crowns of other trees as a function of the focal tree’s neighbourhood radius. The focal tree’s position is either taken as the original stem coordinates or, alternatively, as the centroid (not shown) of the unshaded part of its crown. For the largest tree (orange) these two positions coincide.

The **R-scripts** and functions in the **GitHub** repository are described in the **README.pdf** and may be used to reproduce Fig. 1 using the small test data set (FULL_test.txt).

Executing **0_Impressionistic.R** produces binary files Fig_1_xxppca_c_yy_d_zz

xx: Number of points per crown
yy: non-spatial 0.00, spatial 1.00
zz: delta_d (0 = complete overlap, 1.2 highly assymetric)

The spatial version (FIG_1_100ppca_c_1.00_d_1.20) used to draw Fig. 1 as shown is included and may be directly used to redraw Fig. 1.

Executing **Fig_1.R** produces the figure, evaluates the neighbourhood of the red focals with Tags 1 and 5 (sourcing(**1_Loop.R**) and **2_NN.R**) and outputs their neighbourhood information at neighbourhood radii of 1, 5, 10, 15 and 20 m.

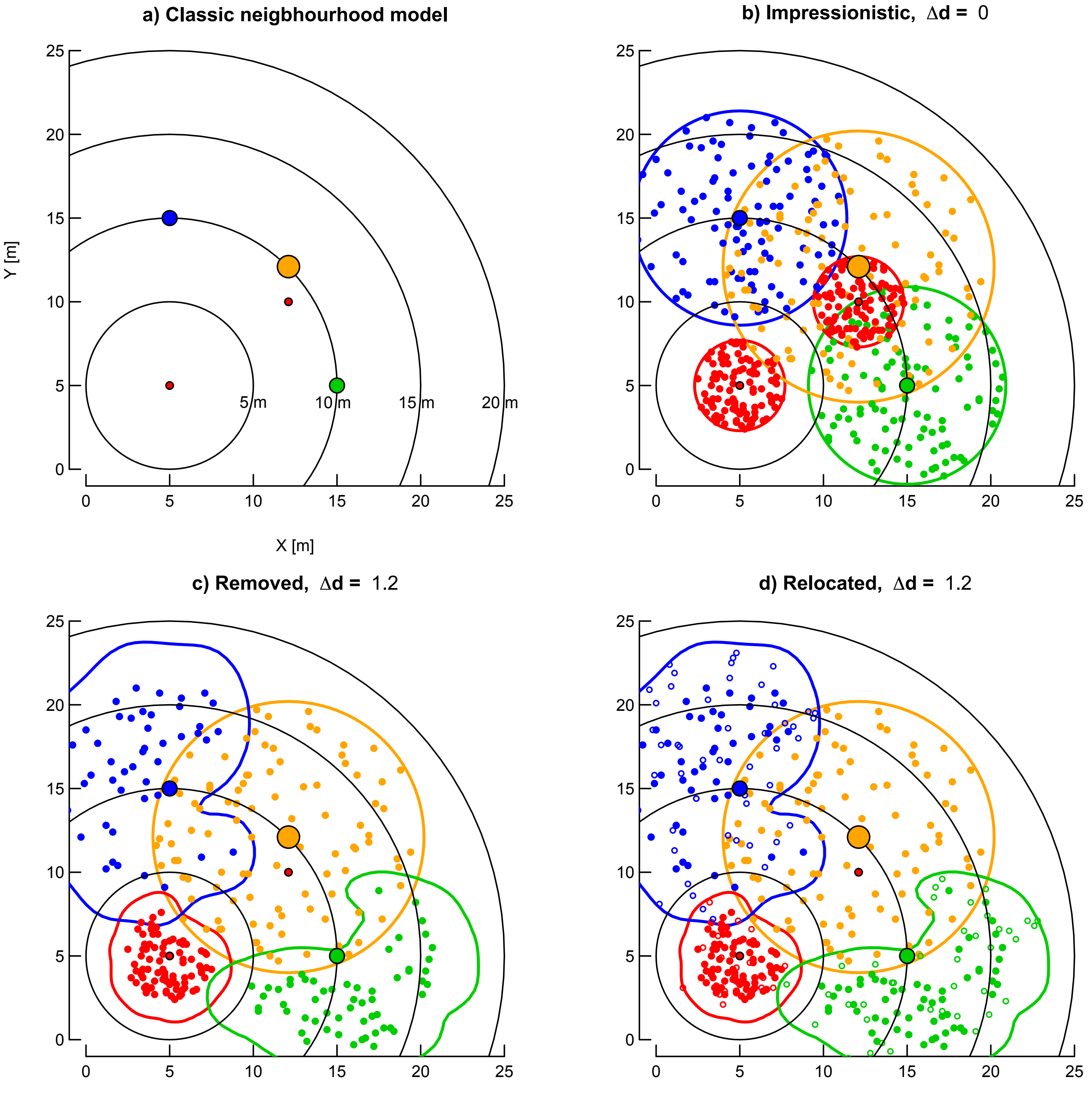

a) Classic neigbhourhood model
b) Impressionistic, Δd = 0
c) Removed, Δd = 1.2
d) Relocated, Δd = 1.2
Y [m]
X [m]
5 m
10 m
15 m
20 m
0
5
10
15
20
25
