## Supplementary material for "Including tree spatial extension in the evaluation of neighbourhood competition effects in Bornean rain forest": nncrexm appendices: App3_hist-ests.pdf (from another version of this preprint: version 2, 2020-08-04)

Appendix 3.

Frequency distributions of the fitted CON coefficients for the 48 species neighbourhood (lin-decay) 'larsm/reloc/crown' models:

Fig. S1A growth in $P_1$
S1B growth in $P_2$
S2A survival in $P_1$
S2B survival in $P_2$

Count is the number of fitted combinations of radius and $\Delta$d fits within the range of 2$\Delta$AICc.

### App_3:Fig_S1A (growth P1) − page 1

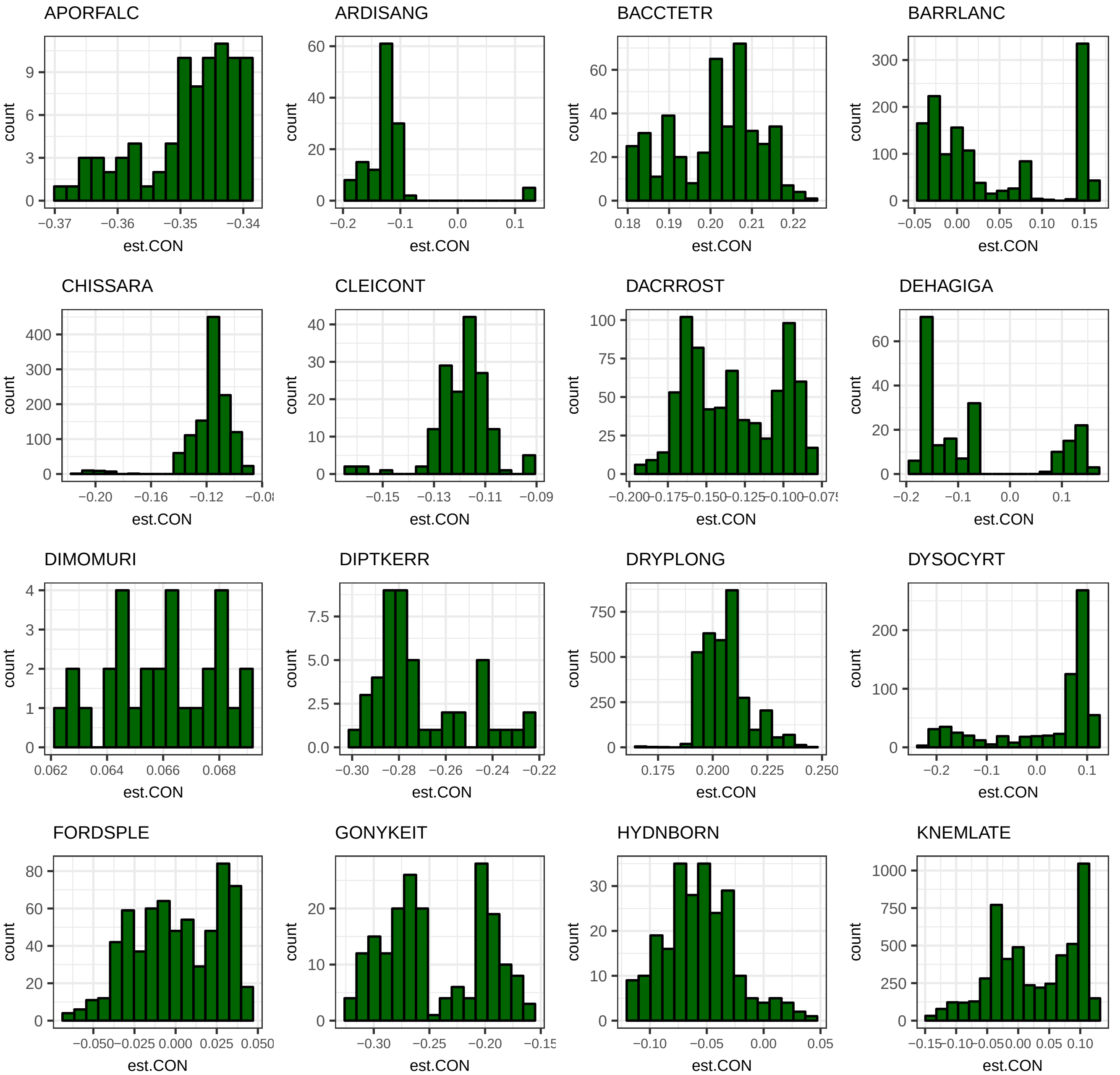

App_3:Fig_S1A (growth P1) − page 2

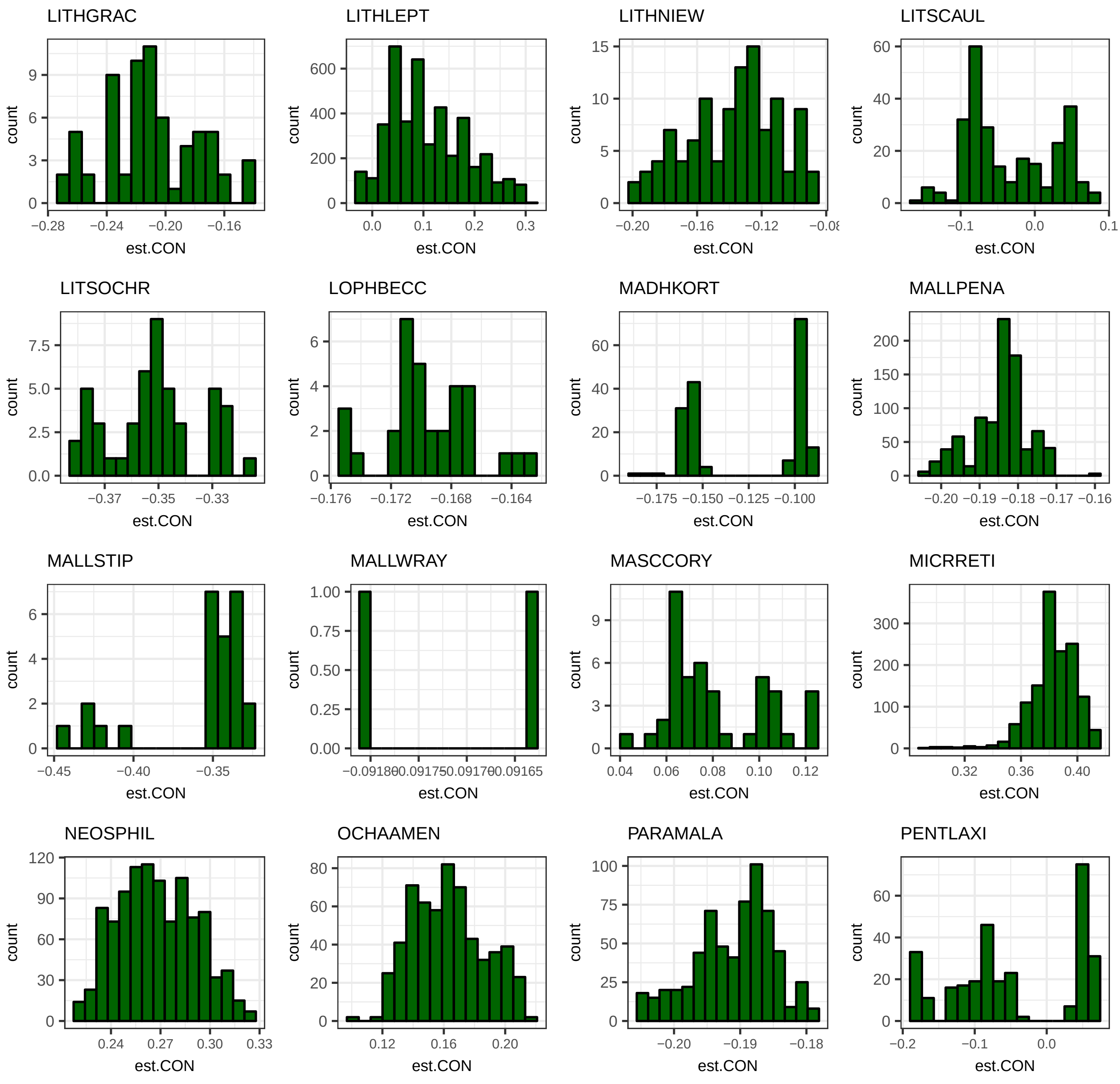

App_3:Fig_S1A (growth P1) − page 3

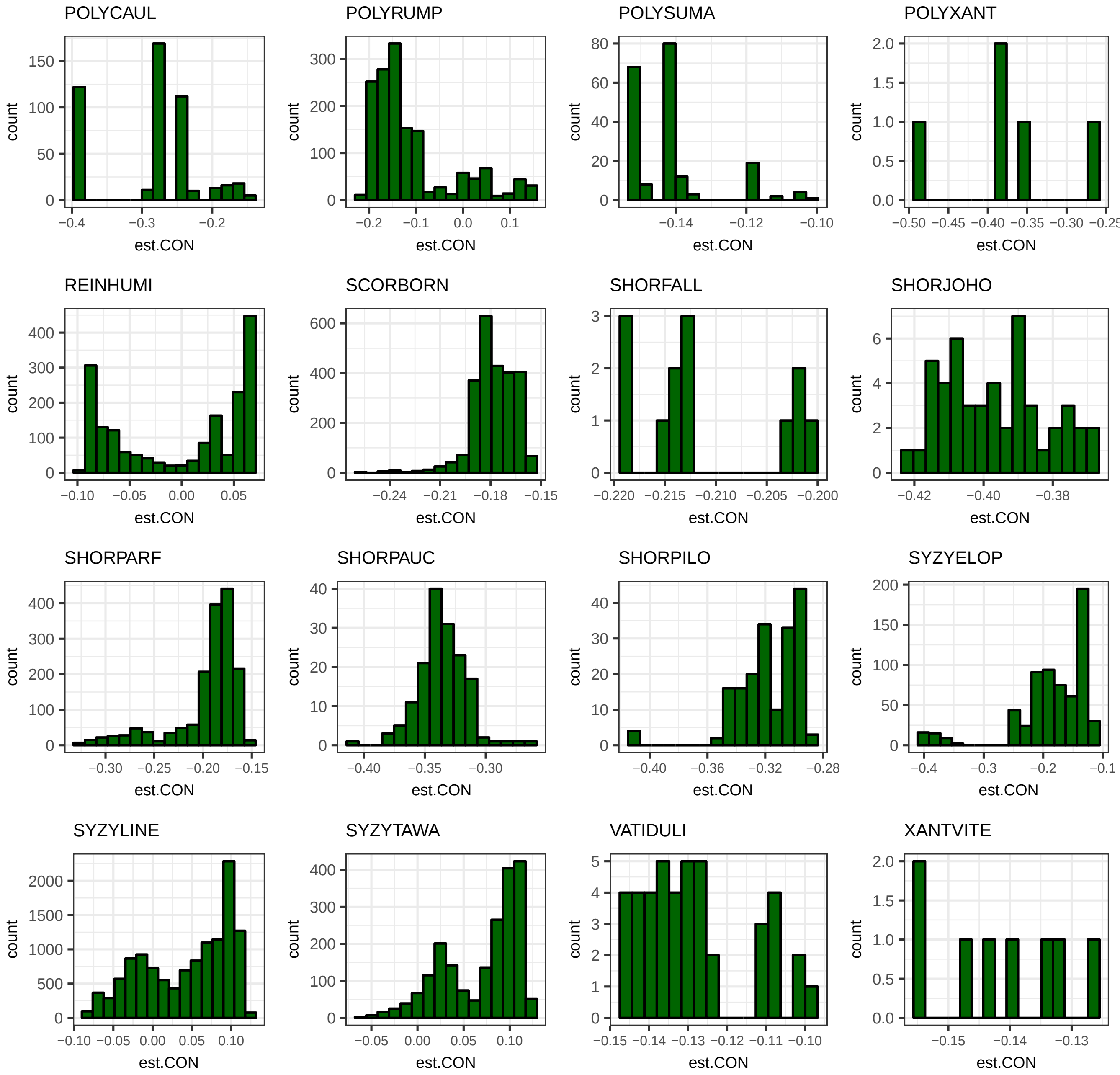

App_3:Fig_S1B (growth P2) − page 1

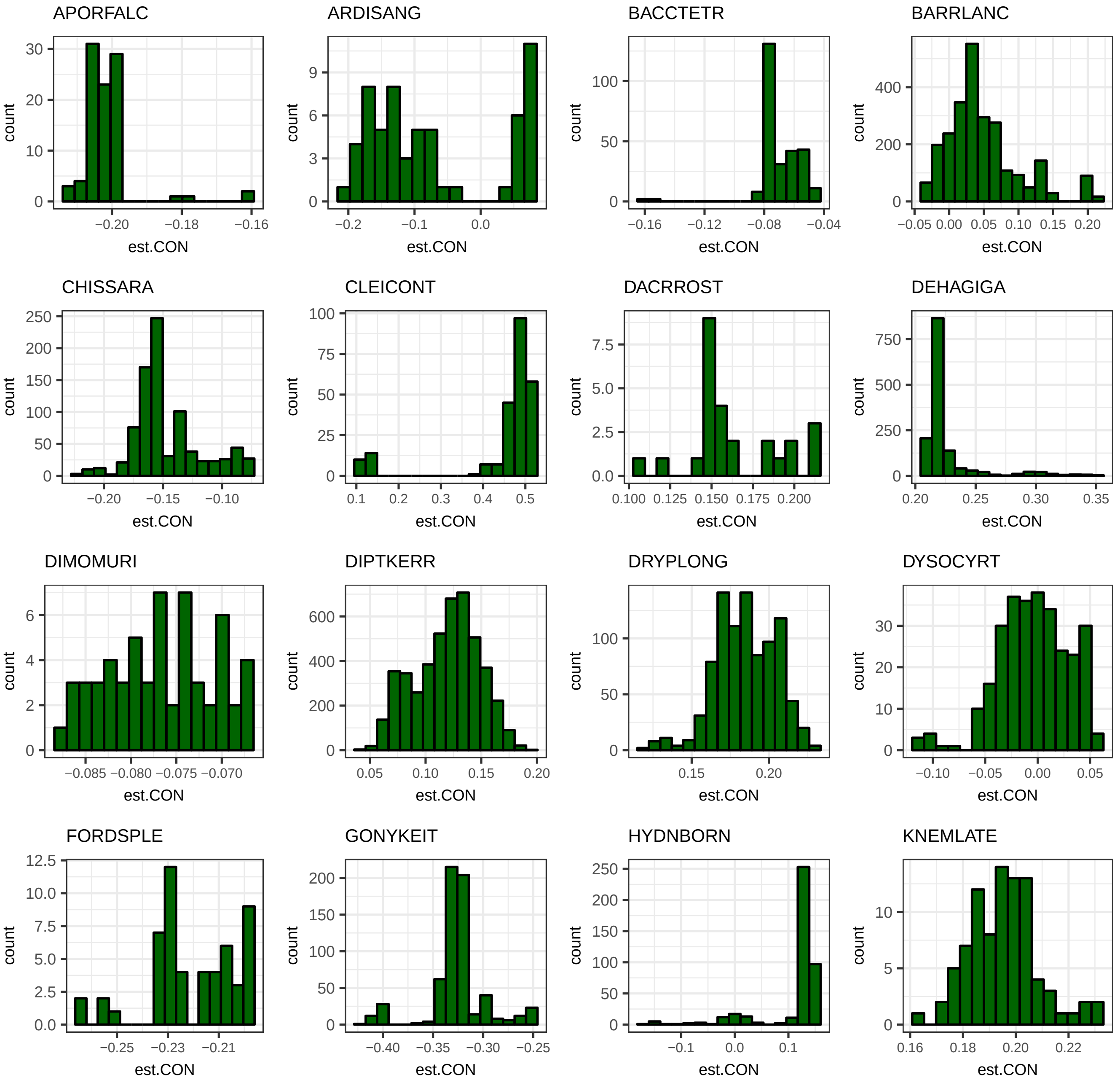

### App_3:Fig_S1B (growth P2) − page 2

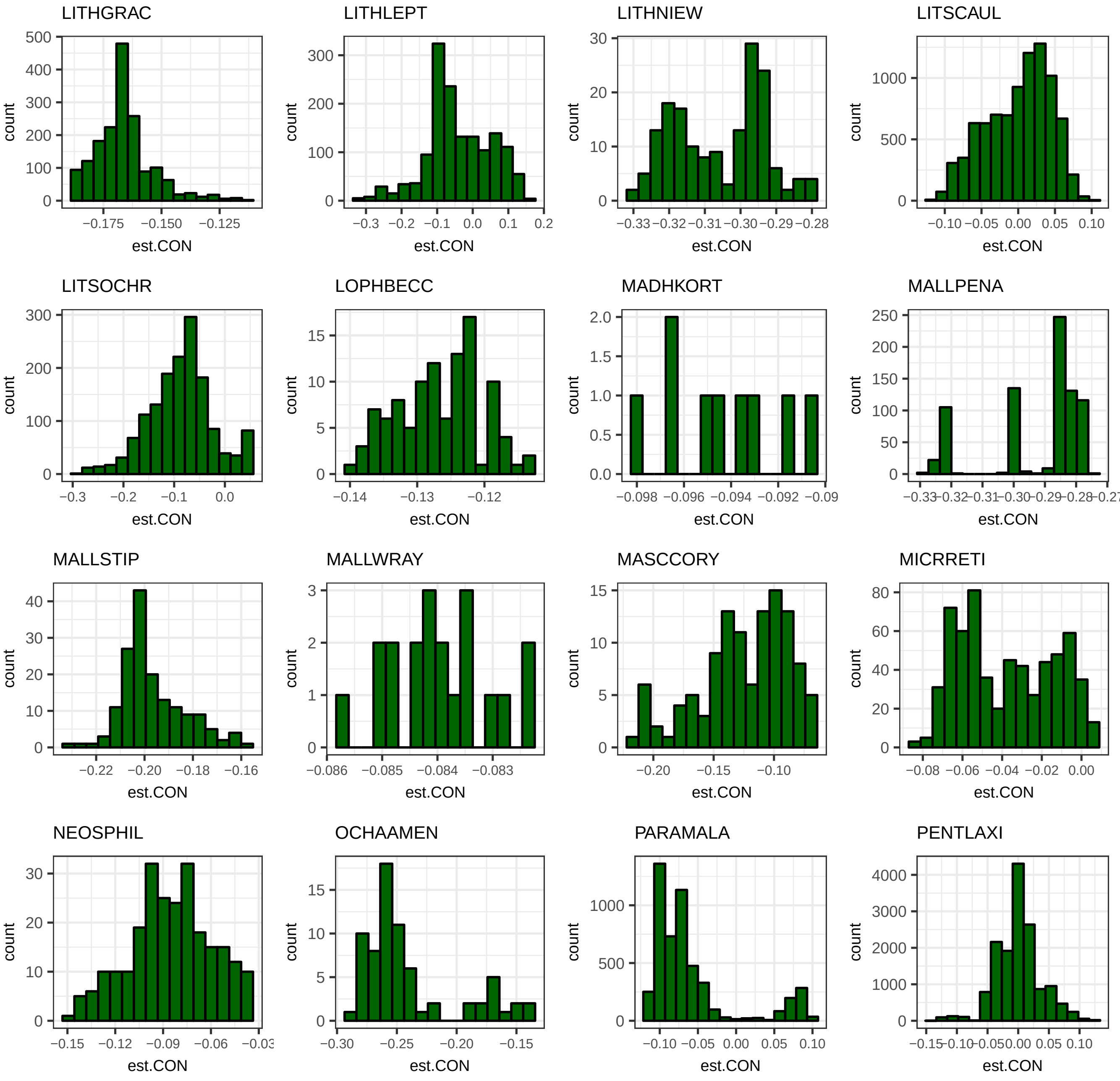

### App_3:Fig_S1B (growth P2) − page 3

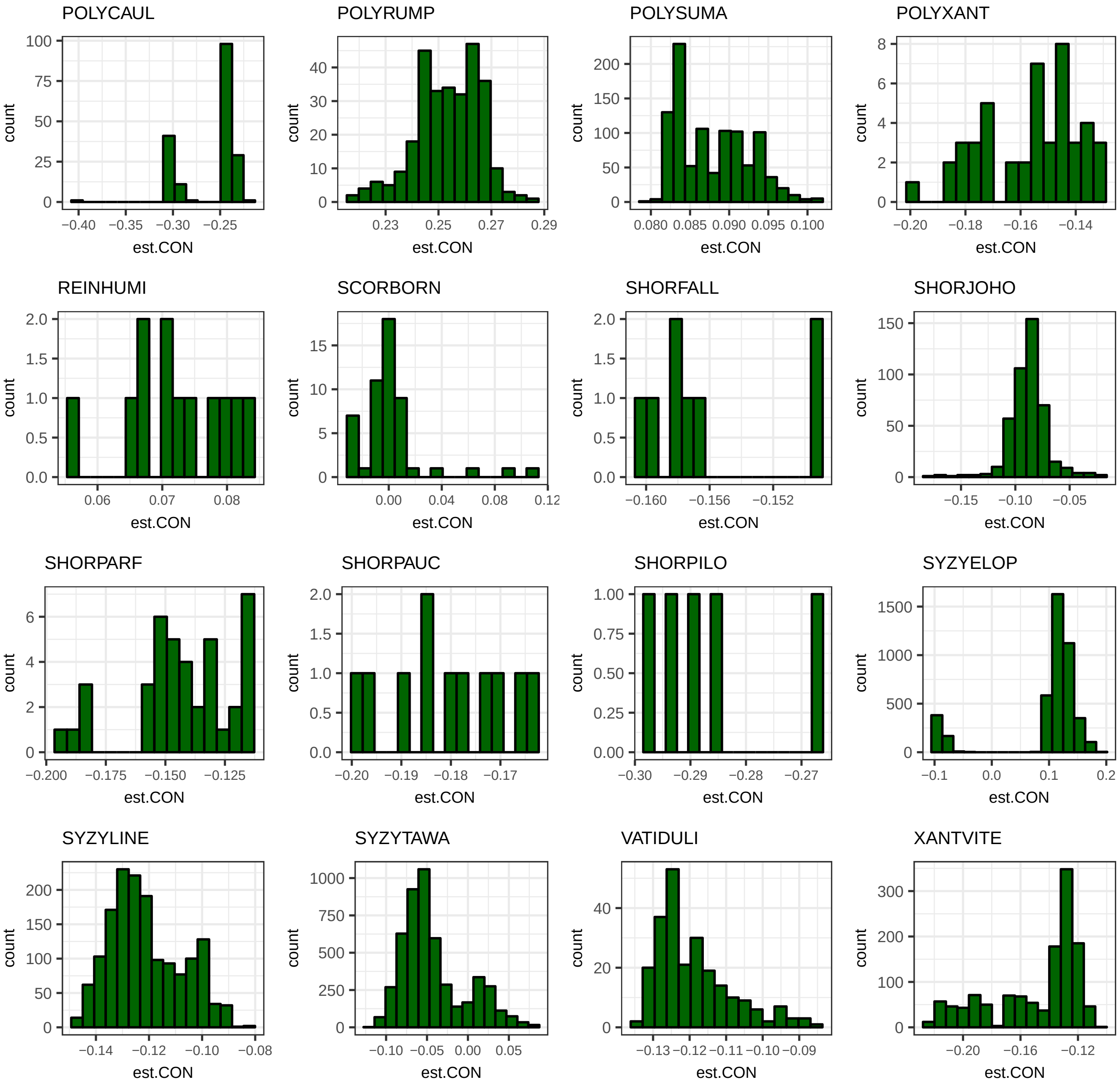

### App_3:Fig_S2A (survival P1) − page 1

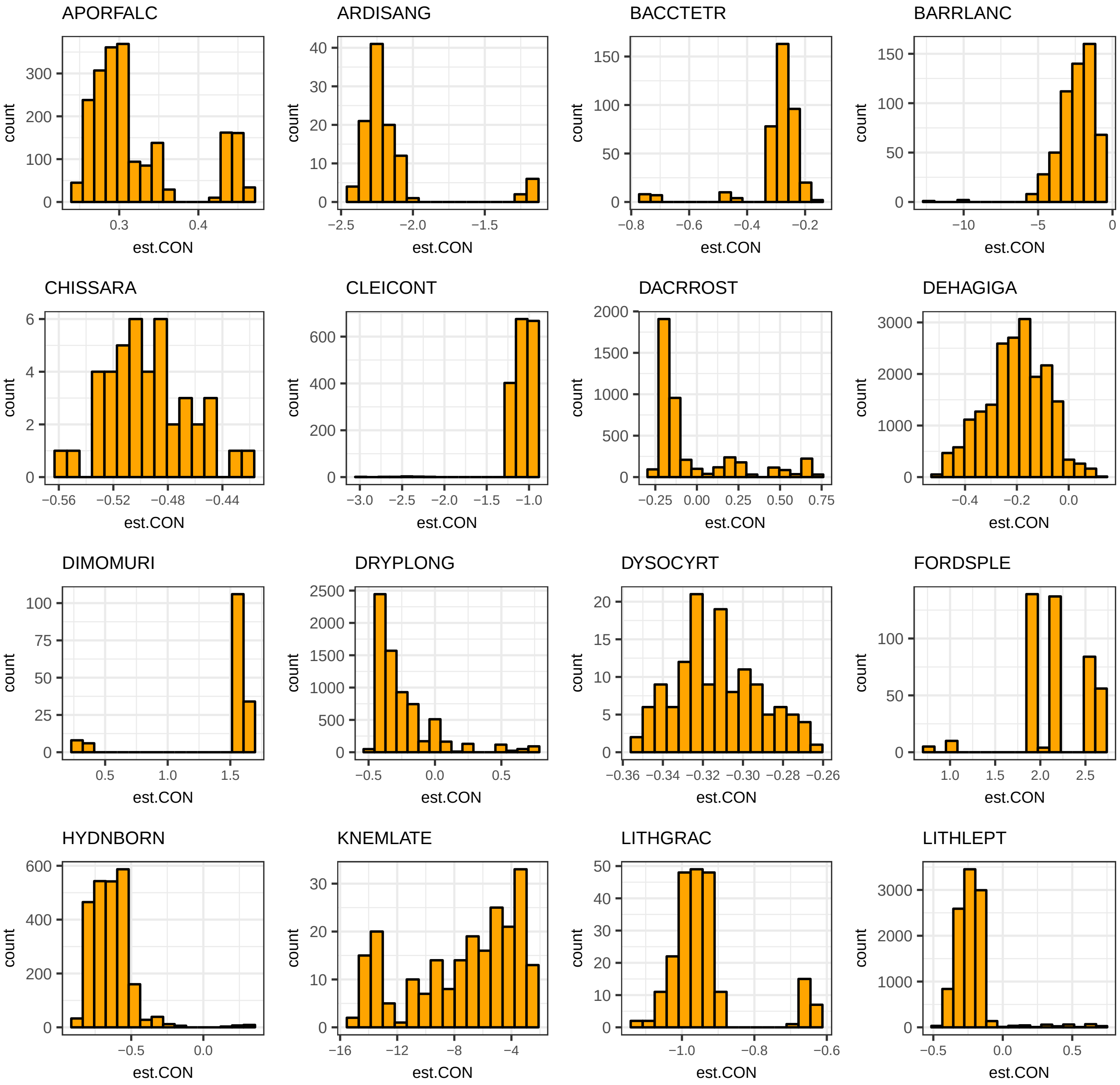

App_3:Fig_S2A (survival P1) − page 2

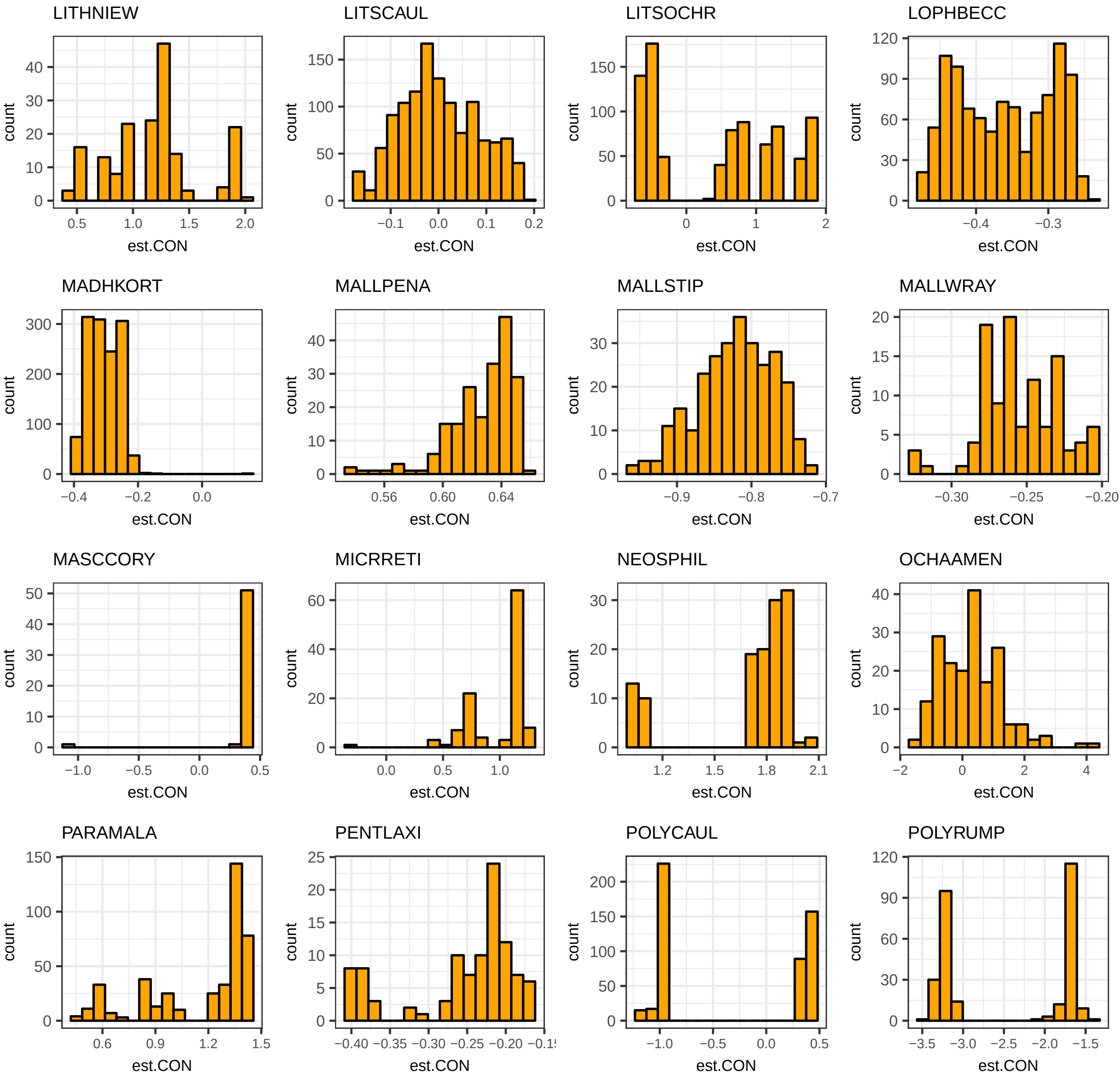

### App_3:Fig_S2A (survival P1) − page 3

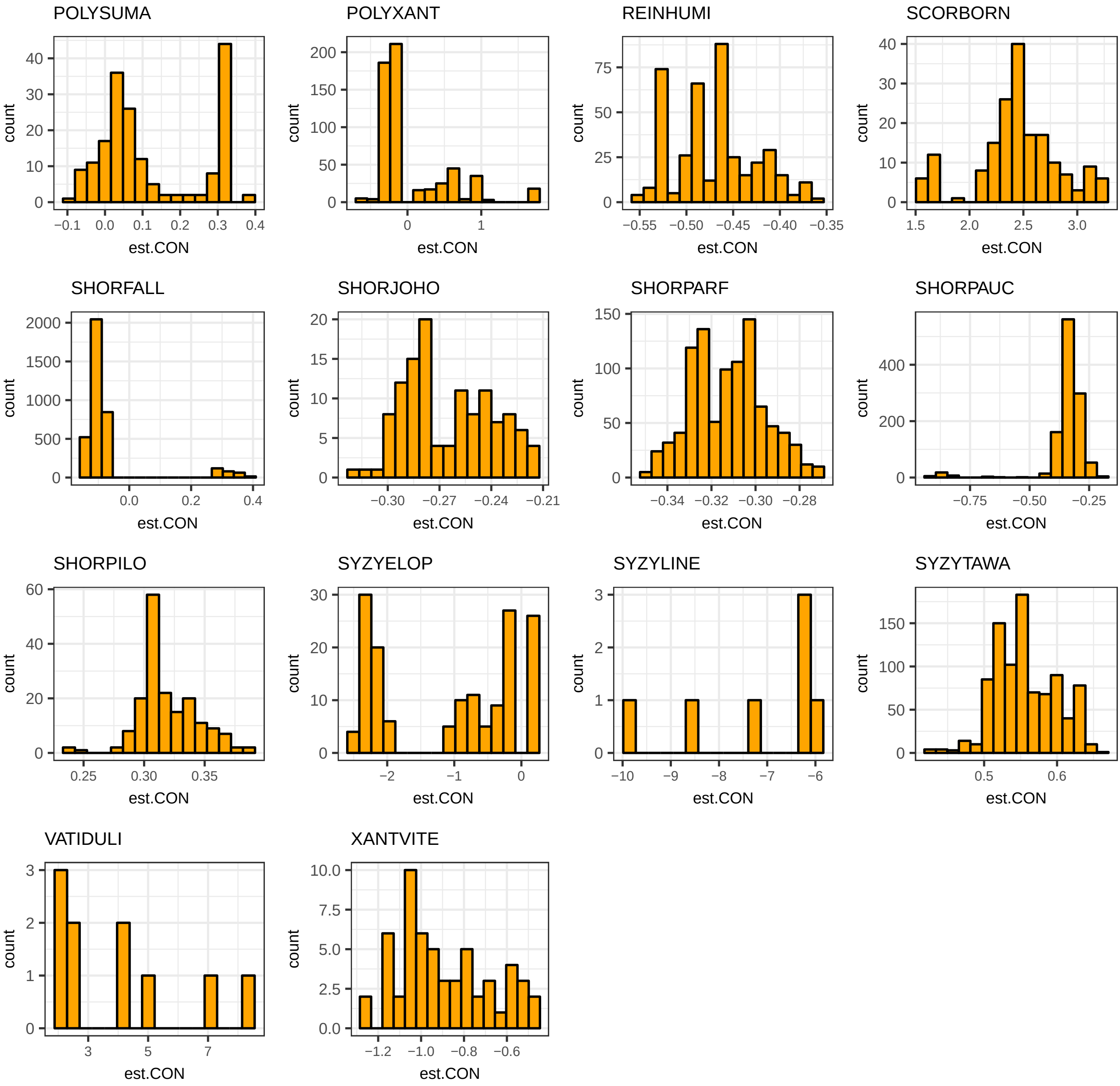

### App_3:Fig_S2B (survival P2) − page 1

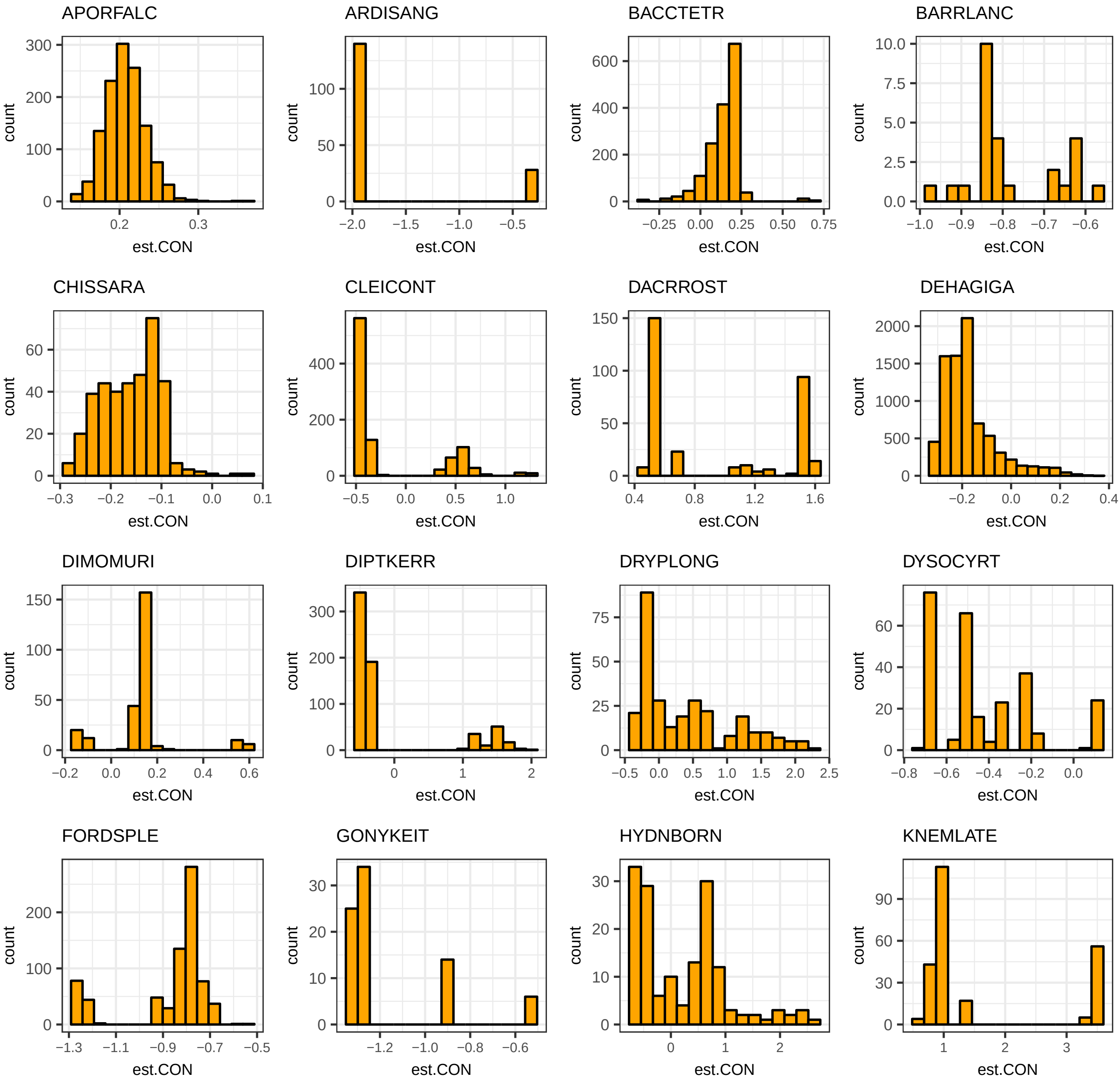

### App_3:Fig_S2B (survival P2) − page 2

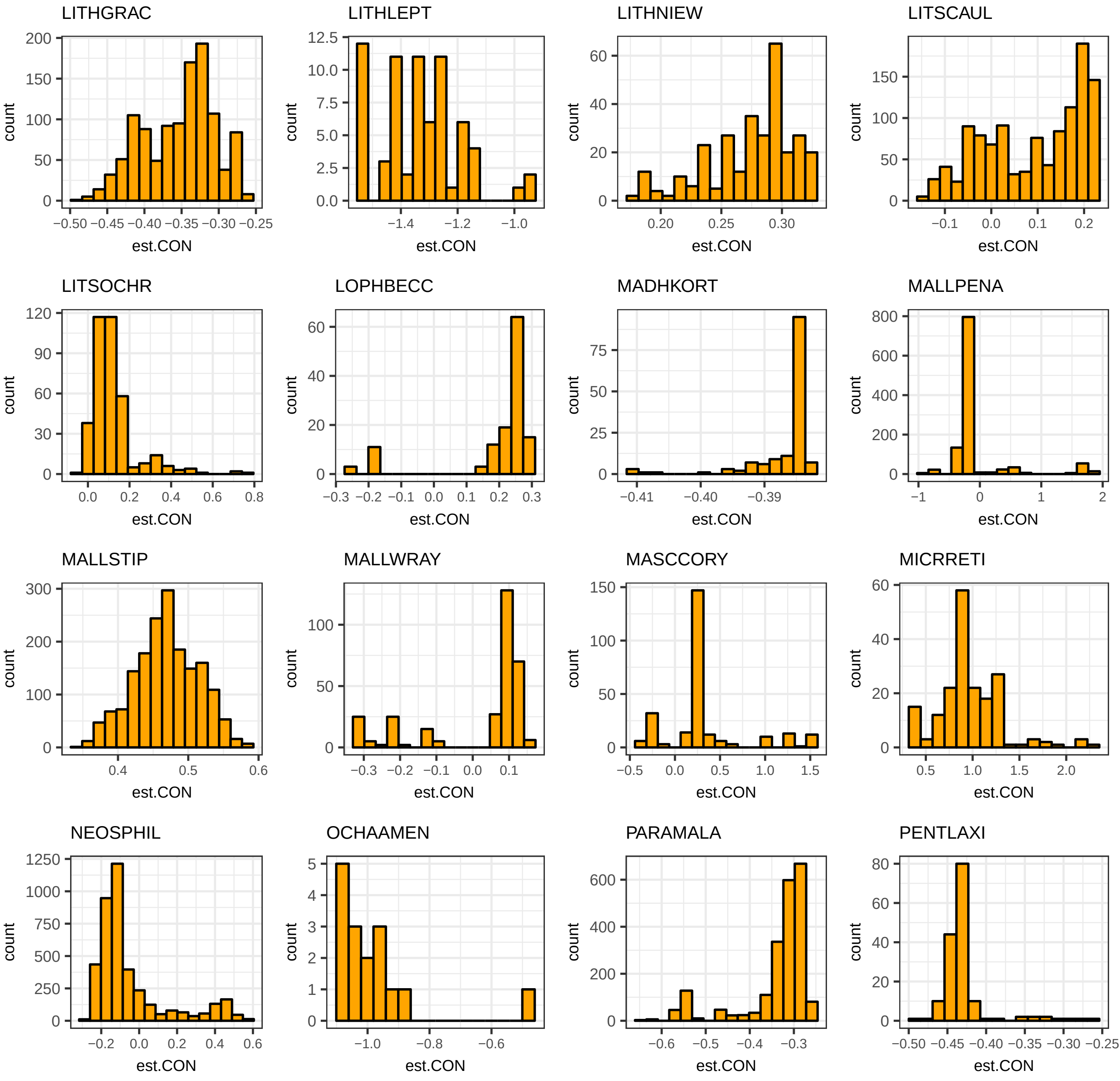

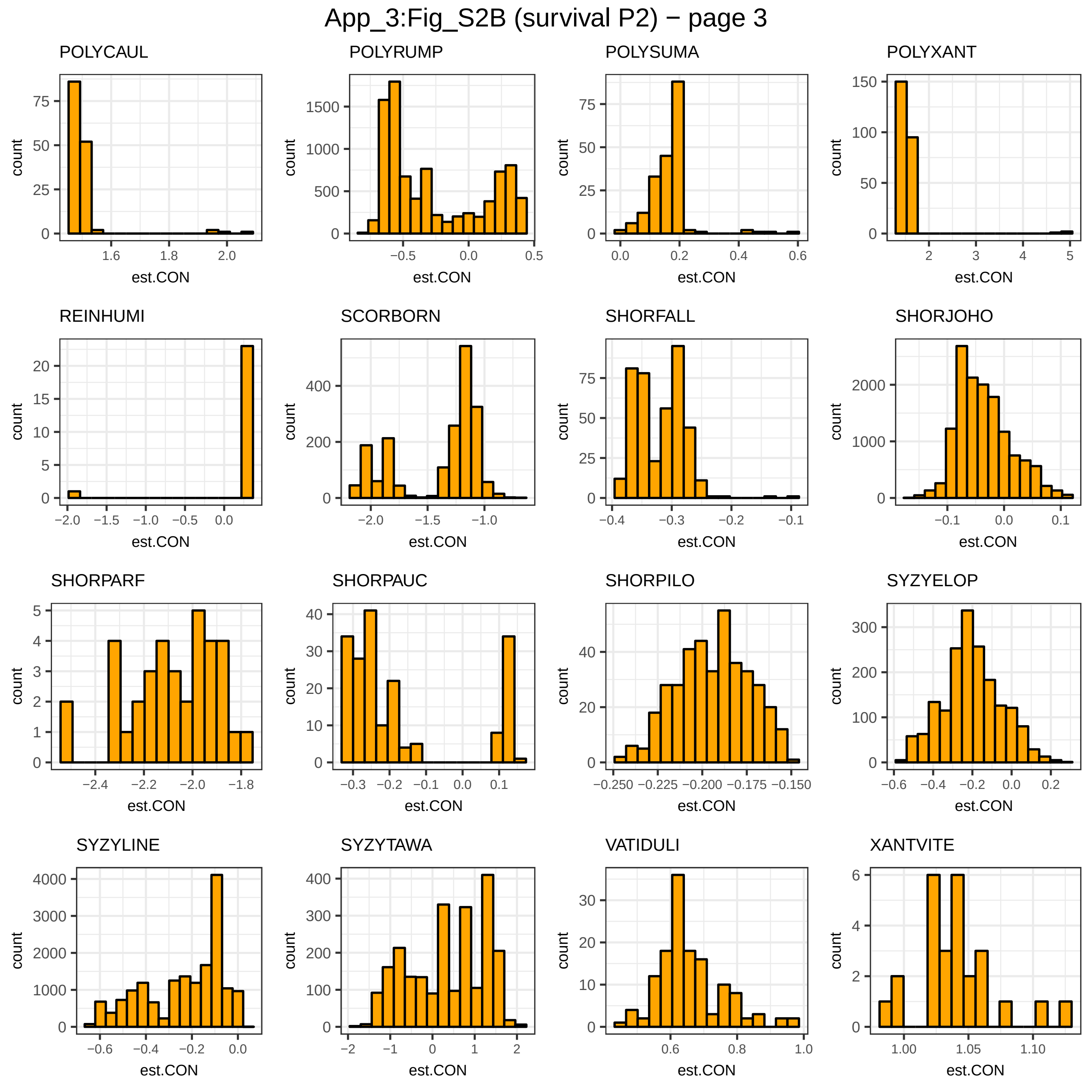
App_3:Fig_S2B (survival P2) − page 3
POLYCAUL
POLYRUMP
POLYSUMA
POLYXANT
REINHUMI
SCORBORN
SHORFALL
SHORJOHO
SHORPARF
SHORPAUC
SHORPILO
SYZYELOP
SYZYLINE
SYZYTAWA
VATIDULI
XANTVITE
count
est.CON
