## Supplementary material for "Including tree spatial extension in the evaluation of neighbourhood competition effects in Bornean rain forest": nncrexm appendices: App4_fit-stats.pdf (from another version of this preprint: version 2, 2020-08-04)

**Appendix 4. Results of regression model fitting: non-spatial versus spatial.**

Four tables of the coefficients, measures of statistical fit and significances, and best fitting radii, of the regression outcomes for all species, non-spatial and two (‘removed’ and ‘relocated’) spatial models, for growth and survival responses in periods 1 and 2 (Excel files).

One table of the effect sizes for CON and HET terms in the non-spatial and all eight combinations of spatial models, for all species, growth and survival responses in periods 1 and 2. (Excel file).

These files are too large to reproduce here when converted to pdf form, but will be available in the Supplementary materials.

App4_GROWTH P1.xls
App4_GROWTH P2.xls
App4_SURVIVAL P1.xls
App4_SURVIVAL P2.xls

App4 Effect Sizes.xls
