## Supplementary material for "Including tree spatial extension in the evaluation of neighbourhood competition effects in Bornean rain forest": nncrexm appendices: App5_radcoeff.pdf (from another version of this preprint: version 2, 2020-08-04)

Appendix 5.

Fitted CON coefficients versus corresponding best-fit radius for the 48 species neighbourhood (lin-decay) 'larsm/reloc/crown' models:

| | |
|---|---|
| Fig. S1A | growth in $P_1$ |
| S1B | growth in $P_2$ |
| S2A | survival in $P_1$ |
| S2B | survival in $P_2$ |

Number of points are fitted combinations of radius and Δd fits within the range of 2ΔAICc.

The five species referred to in the main text and footnote to Appendix 1: Fig. 6b are indicated with rad asterisks next to their names in Fig. S2A.

### App_5:Fig_S1A (growth P1) − page 1

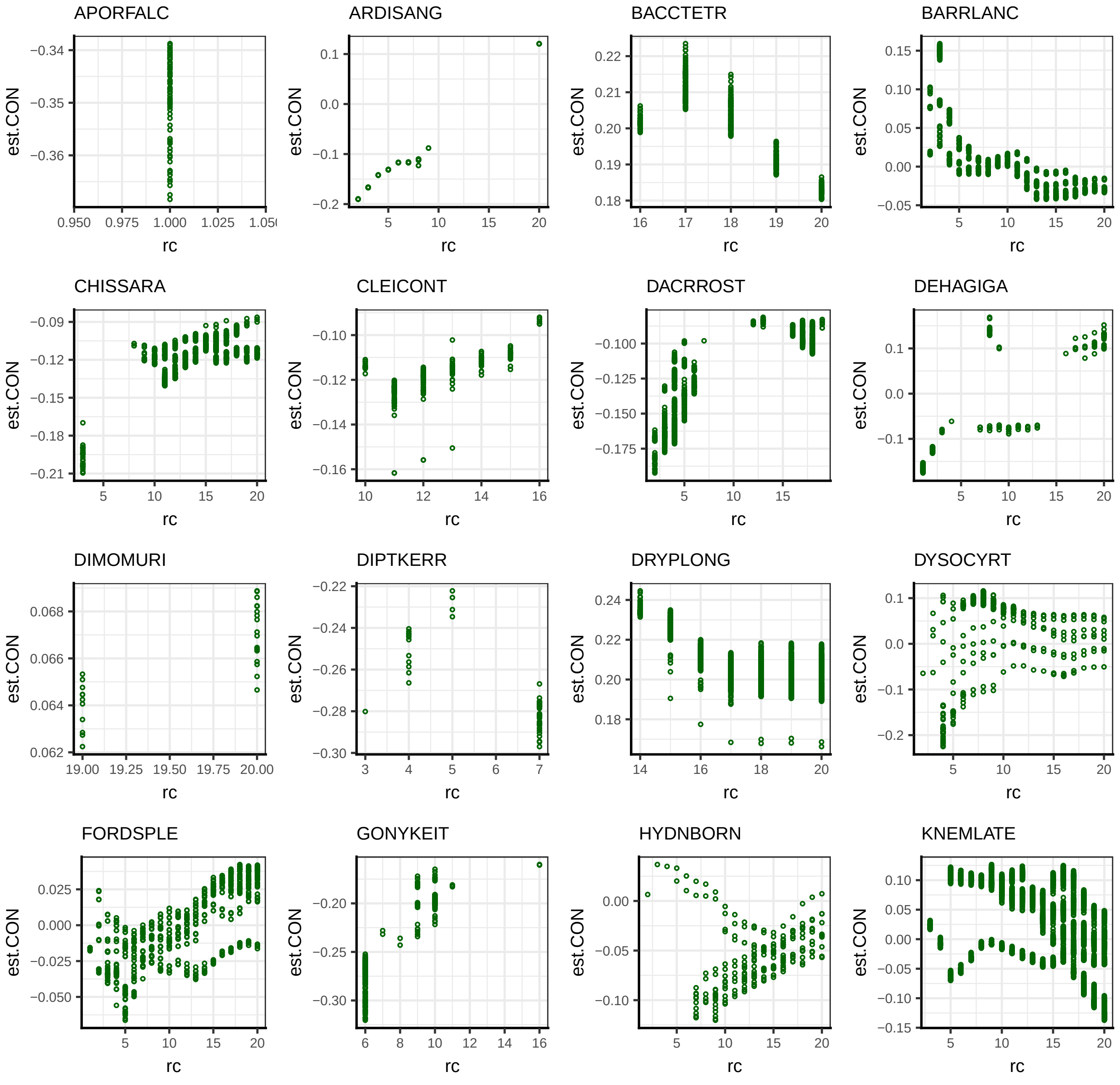

### App_5:Fig_S1A (growth P1) − page 2

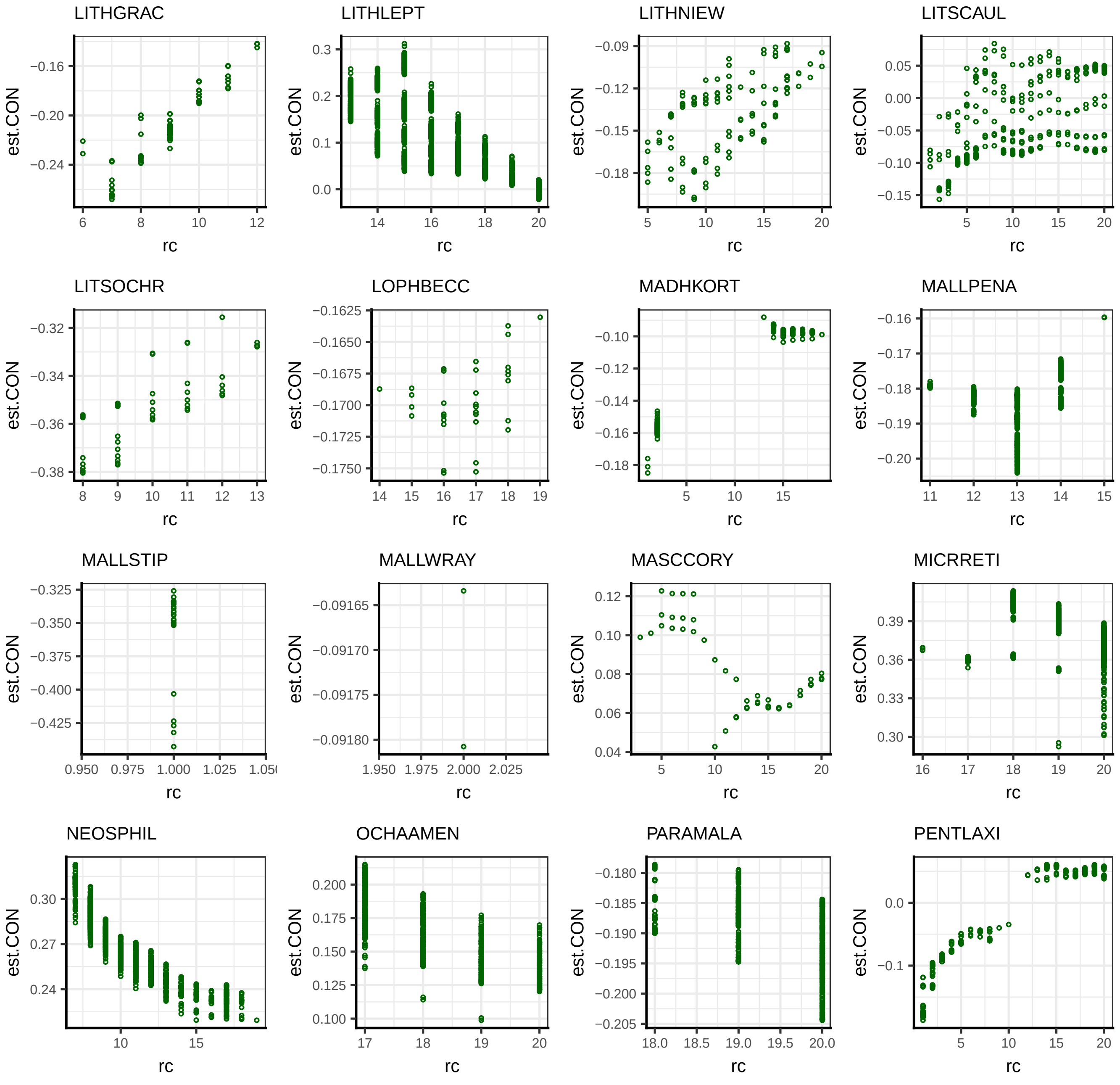

### App_5:Fig_S1A (growth P1) − page 3

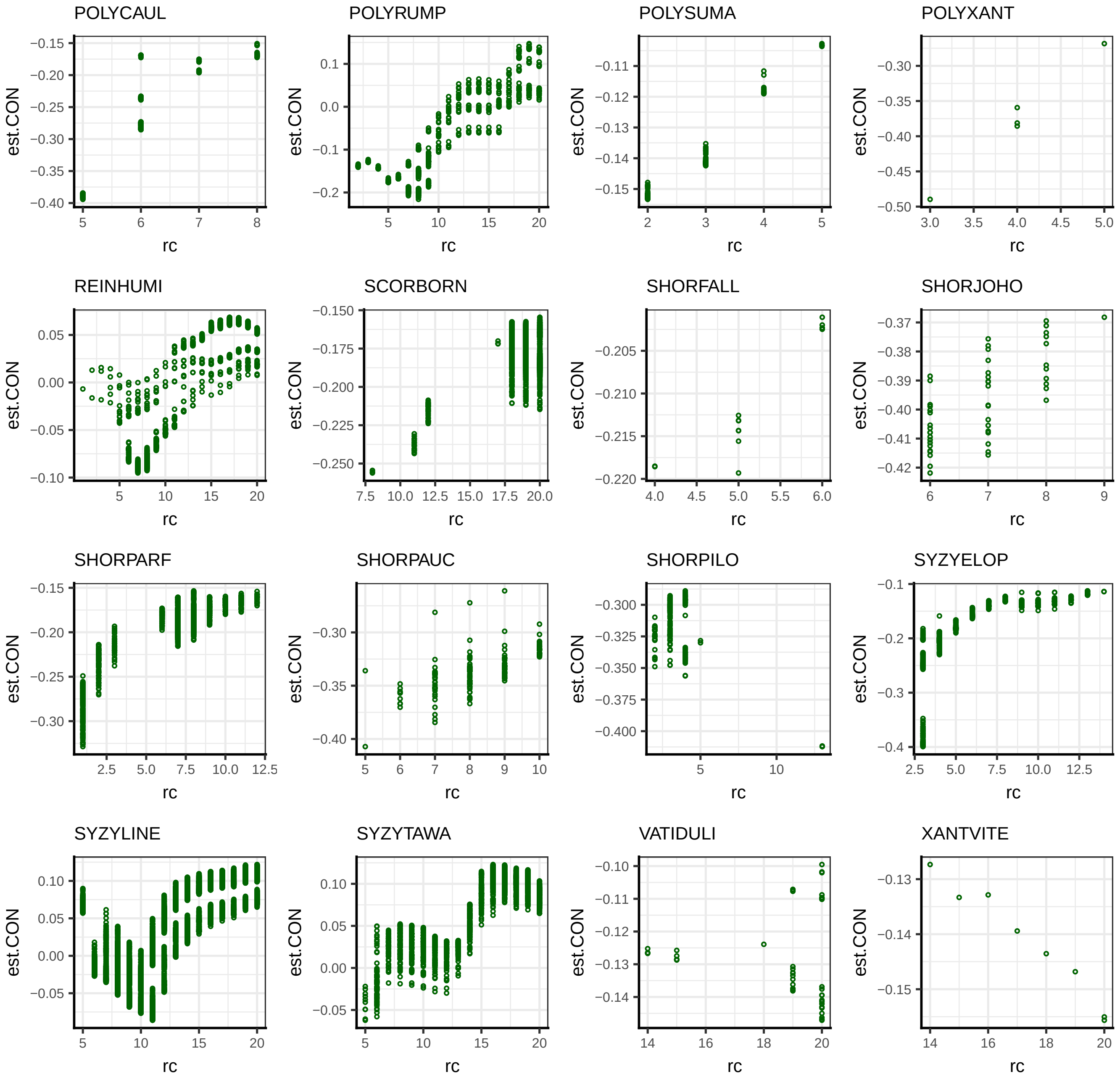

### App_5:Fig_S1B (growth P2) − page 1

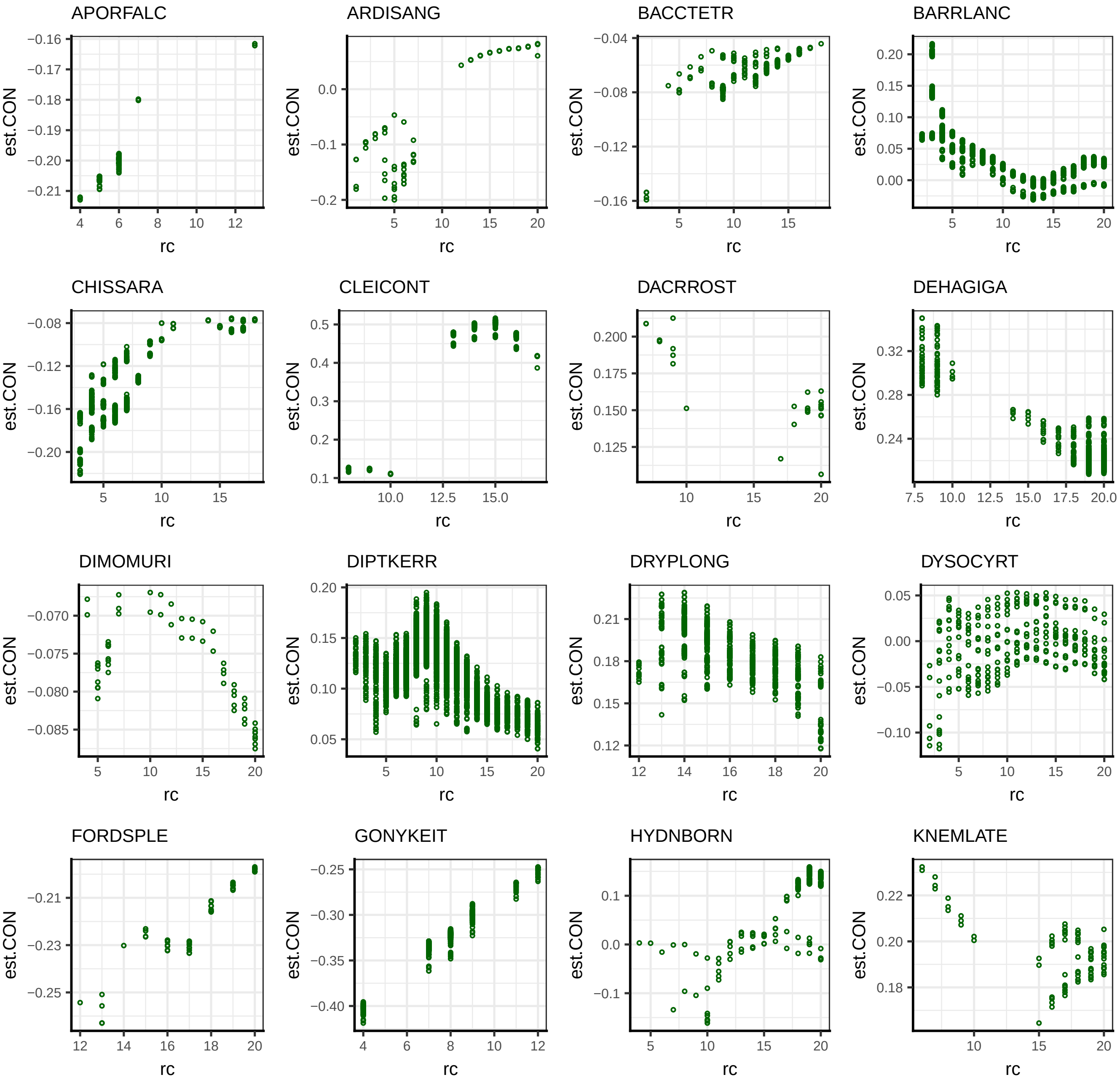

App_5:Fig_S1B (growth P2) − page 2

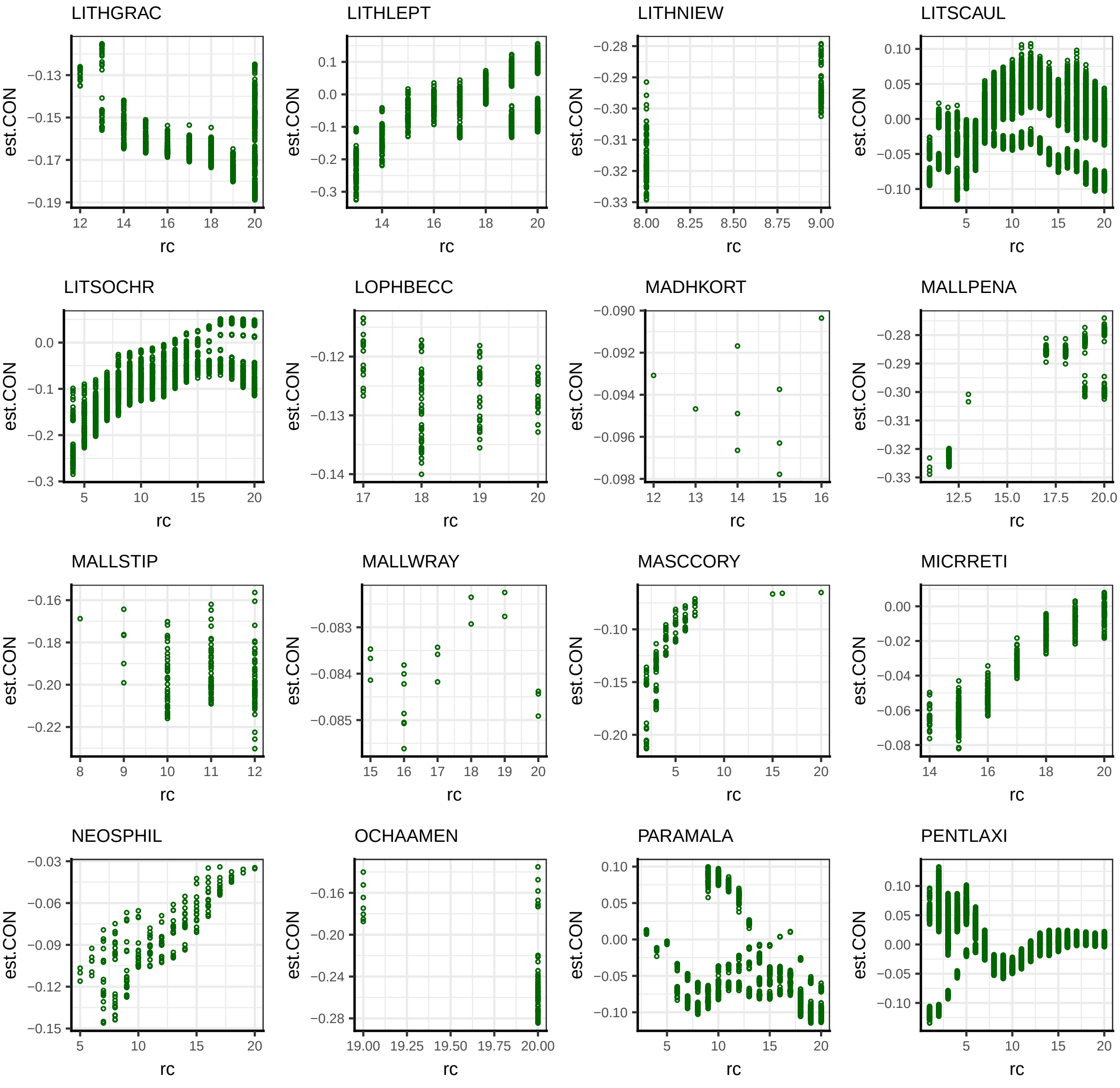

### App_5:Fig_S1B (growth P2) − page 3

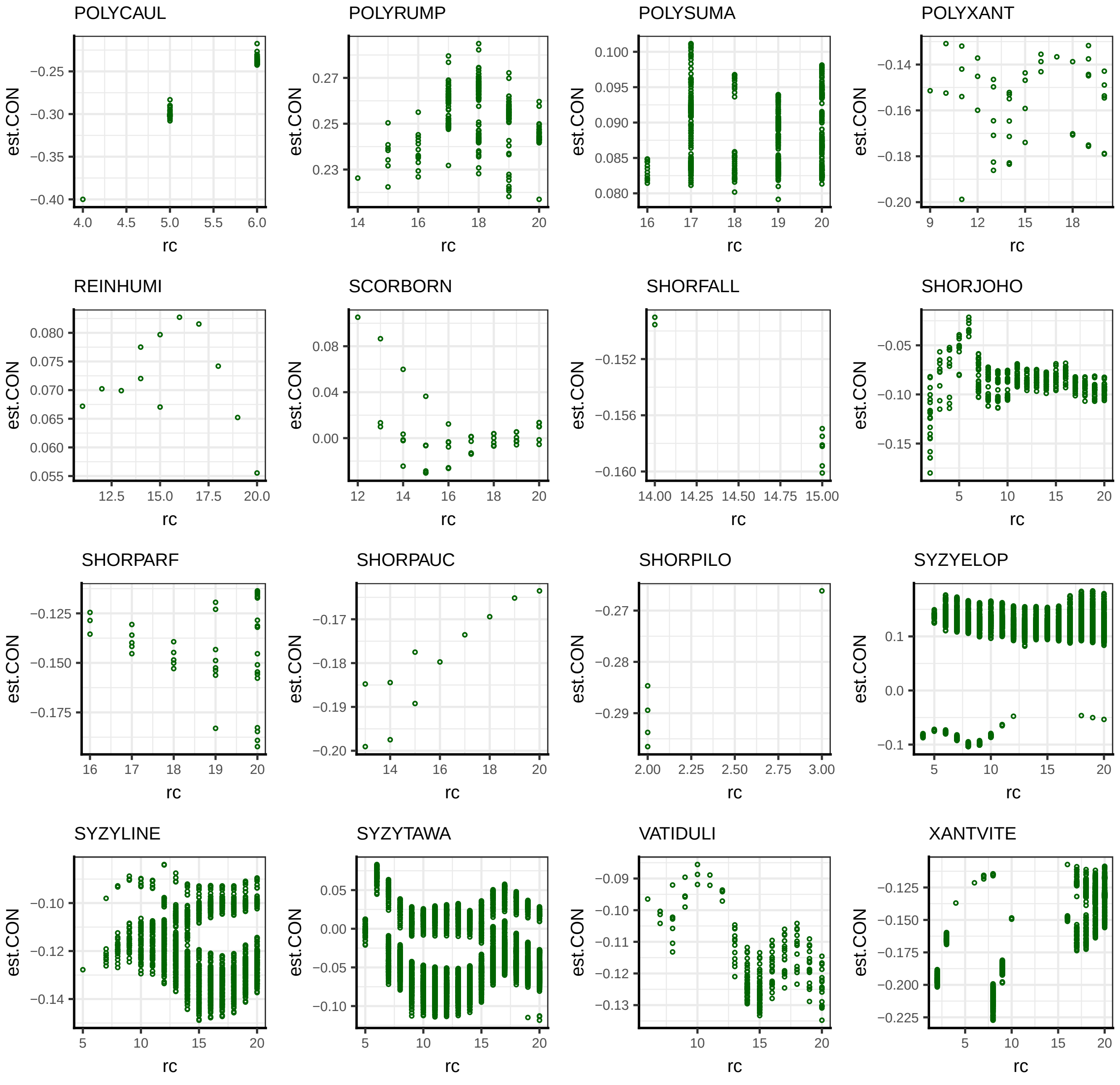

App_5:Fig_2A (survival P1) − page 1

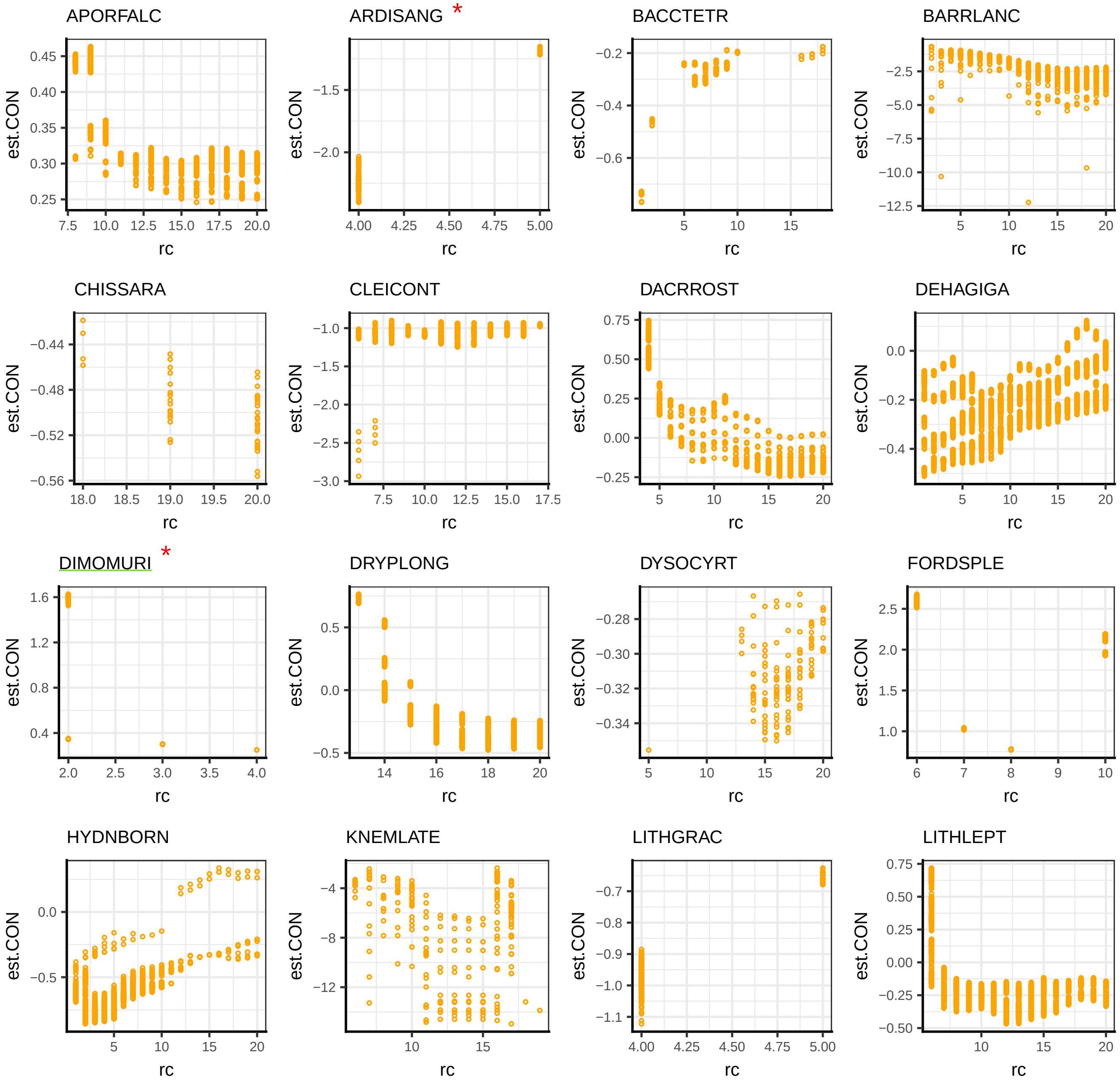

### App_5:Fig_2A (survival P1) − page 2

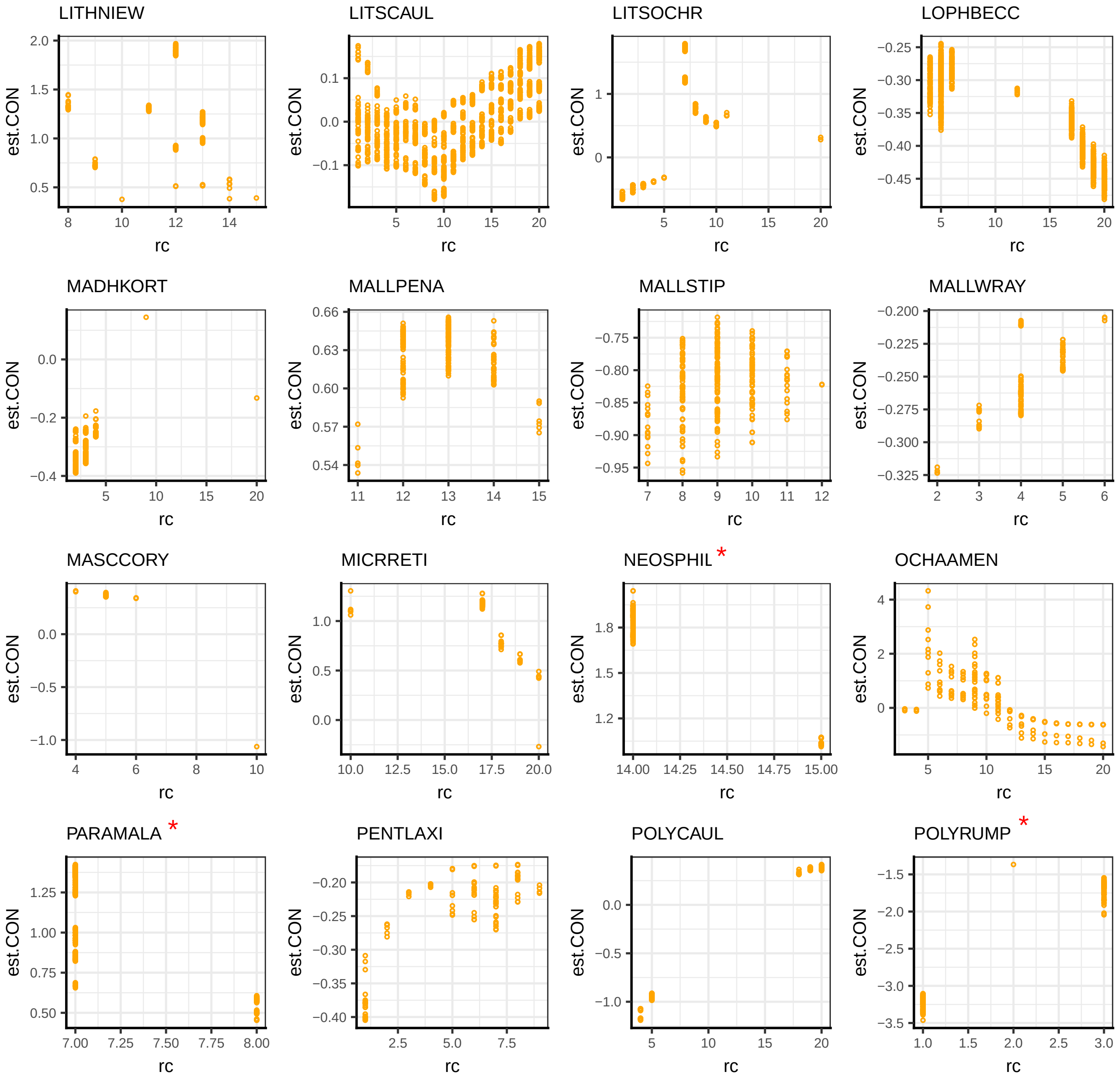

### App_5:Fig_2A (survival P1) − page 3

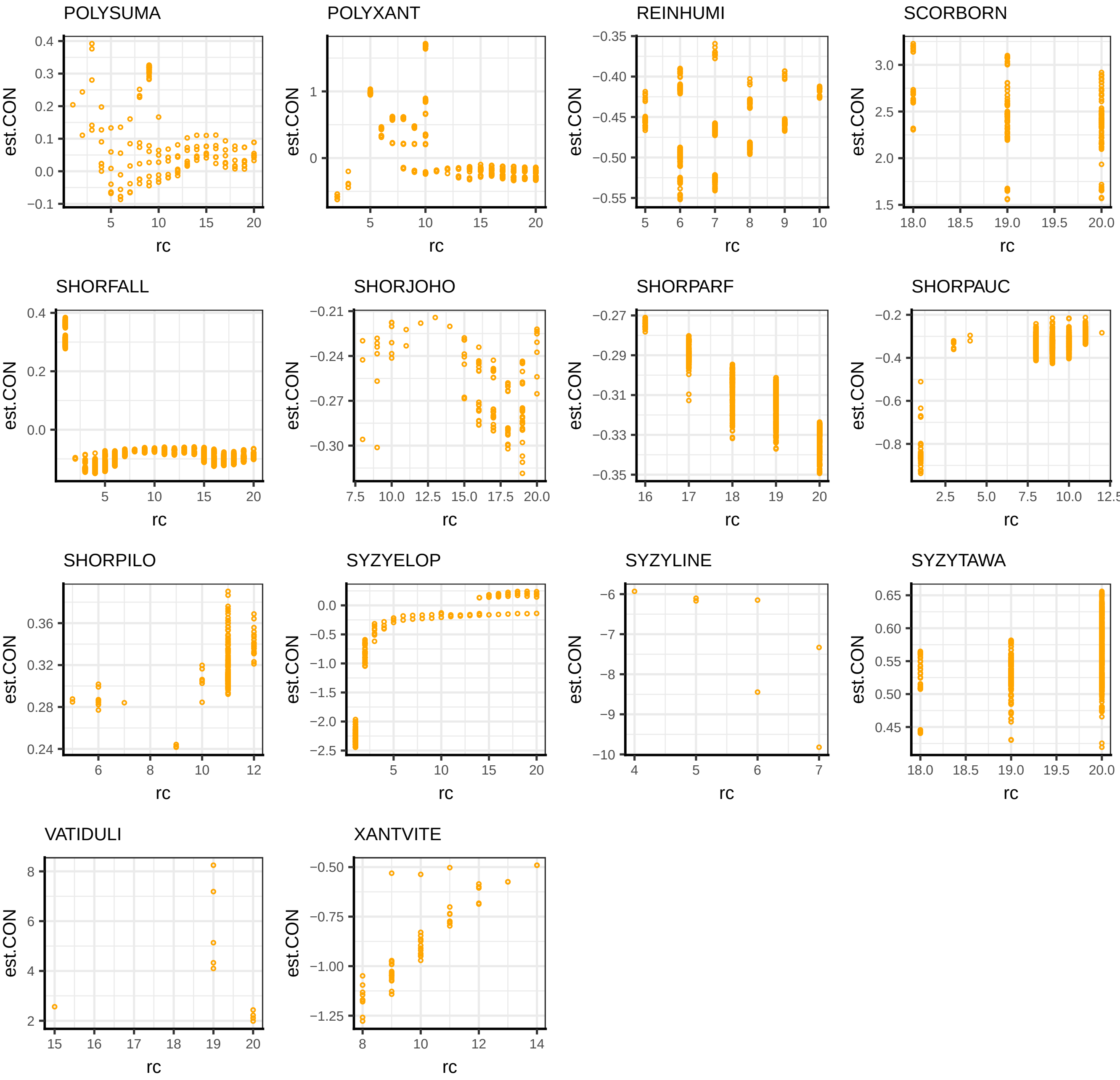

### App_5:Fig_2B (survival P2) − page 1

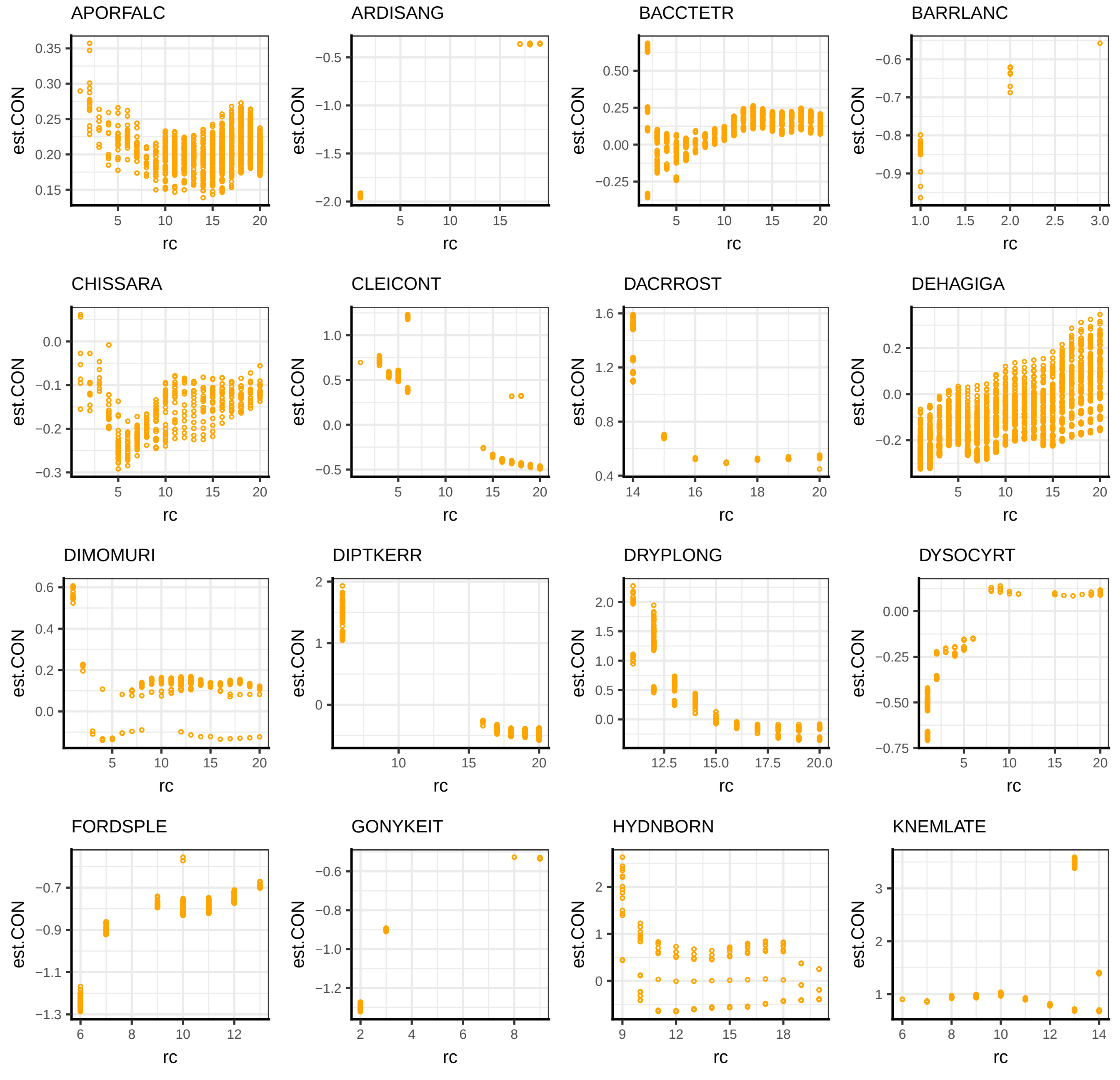

App_5:Fig_2B (survival P2) − page 2

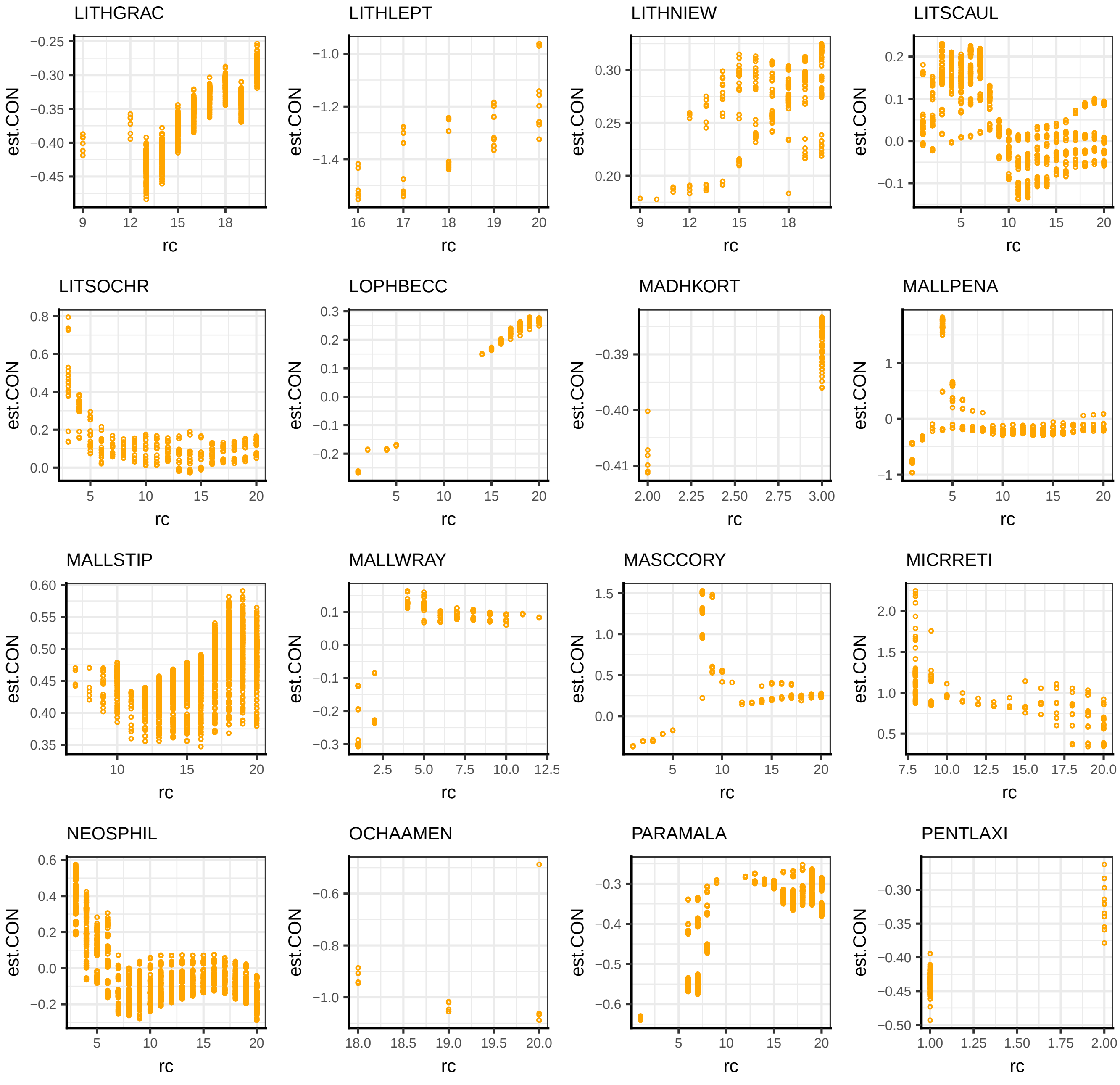

### App_5:Fig_2B (survival P2) − page 3

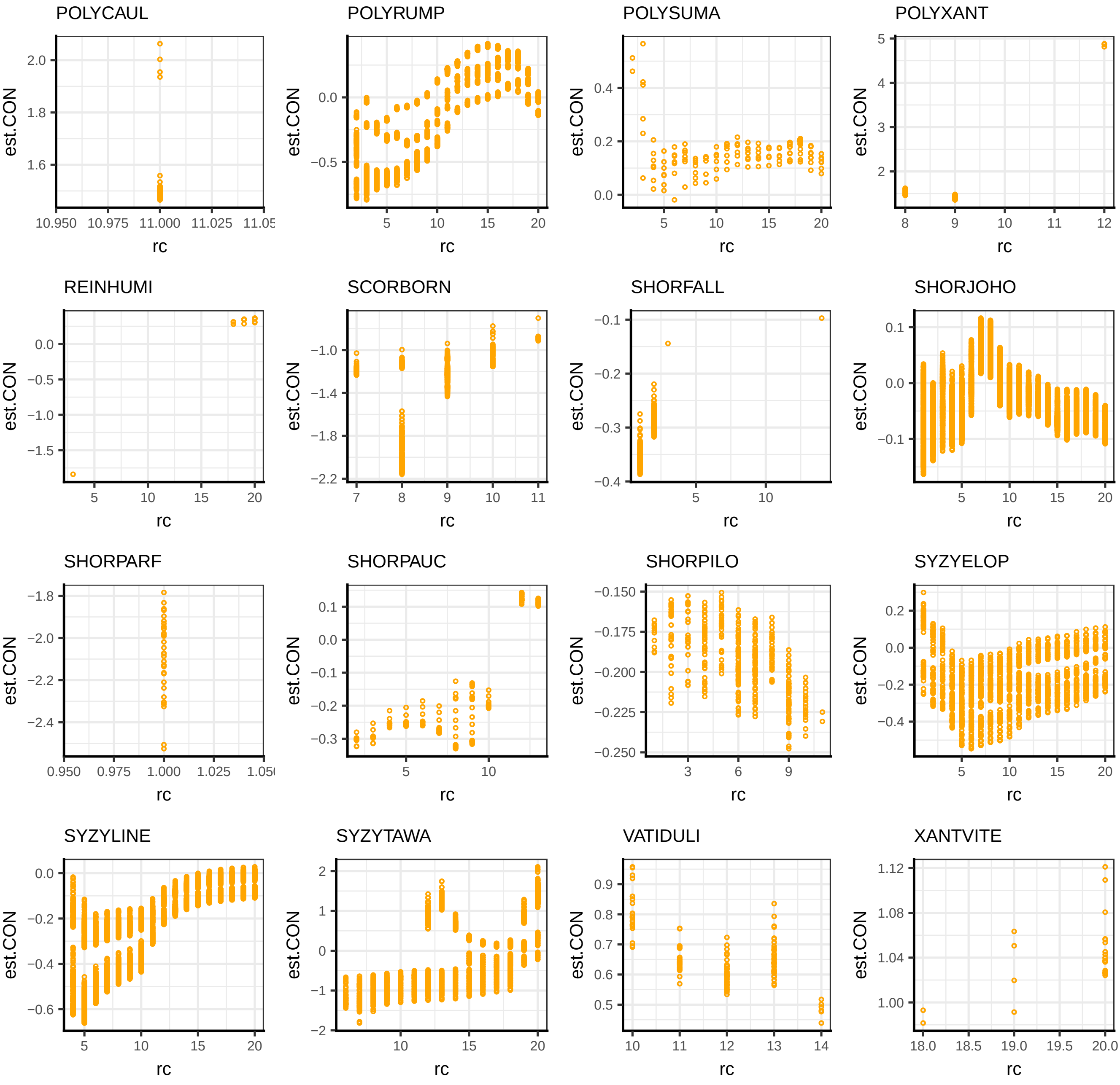
