## Supplementary material for "Including tree spatial extension in the evaluation of neighbourhood competition effects in Bornean rain forest": nncrexm appendices: App6 pattern.pdf (from another version of this preprint: version 2, 2020-08-04)

Appendix 6. Analysis of the role of structure, patterning and dynamics of the species studied in relation to CON and HET effect sizes

1. *Over-understorey structure*

The OUI was significantly negatively correlated with CON effects on growth in both $P_1$ and $P_2$ ($r = -0.371$, $P = 0.022$; $r = -0.405$, $P = 0.012$, respectively) and likewise for HET effects on growth ($r = -0.388$, $P = 0.016$; $r = -0.432$, $P = 0.007$). However, difference in CON and HET effects $P_2 - P_1$ showed no correlation ($r = 0.038$ and 0.034, $P > 0.8$). By contrast CON effects on survival in $P_1$ and $P_2$ were positive yet insignificant or negatively weaker ($r = 0.164$, $P = 0.33$; $r = -0.305$, $P = 0.063$, respectively) with HET effects on survival slightly positive in $P_1$ but not $P_2$ ($r = 0.281$, $P = 0.087$; $r = -0.249$, $P = 0.13$). The means of the OUI-classes are shown in Table S1; Figs 2-4, main text, show the individual values color-coded to match the lines in Appendix 1: Table S1. The patterns of species' points in the community-level graphs (Figs 2b and 3b, main text, especially) are not therefore accountable by over-understorey structure: differencing CON effects on growth between periods almost completely removed any influence. It is, furthermore, noticeable that CON and HET effects on survival in $P_1$ were, on average, much more positive for the over-storey species and more negative for the under- and intermediate storeys, and this difference was reversed in $P_2$.

The correlations between difference in CON effect on growth $P_2 - P_1$ versus CON or HET effect on survival when separated into the OUI classes were all insignificant (under-, intermediate, over-storey: CON −0.516, −0.113, −0.465; HET −0.448, −0.086, −0.512, respectively; range of $P = 0.059$ to 0.78)).

Table S1. Mean values of the 38 selected species' effect sizes for the three over- understorey index (OUI) classes for the eight combinations of conspecific and heterospecific effect size (CON, HET), growth and survival variables (G, S), and for periods 1 and 2 ($P_1$, $P_2$). OUI classes: under-, < 20; intermediate, ≥ 20 – 55; and over-storey, > 55 (scale of 0 – 100). The classes are correspondingly color-coded in Figs 2-5, main text, and in Appendix 1: Table S1.

| | | OUI: | under- | intermediate | over- |
|---|---|---|---|---|---|
| | | *n* | 14 | 13 | 11 |
| CON | G | $P_1$ | −0.026 | −0.058 | −0.104 |
| | | $P_2$ | −0.017 | −0.029 | −0.091 |
| CON | S | $P_1$ | −0.057 | −0.450 | 0.235 |
| | | $P_2$ | 0.092 | 0.101 | −0.396 |
| HET | G | $P_1$ | −0.031 | −0.055 | −0.118 |
| | | $P_2$ | −0.028 | −0.035 | −0.081 |
| HET | S | $P_1$ | −0.210 | −0.270 | 0.505 |
| | | $P_2$ | 0.363 | 0.131 | −1.056 |

1. *Population dynamics*

Mortality, recruitment and stem relative growth rates, *rgr*, of small trees (10 - < 50 cm *gbh*) for the 38 selected species (D. M. Newbery and others, unpublished data) in $P_1$ were all strongly negatively correlated ($P \leq 0.01$) with CON effects on growth in that same period, but only *rgr* was similarly negatively correlated with CON effects on growth in $P_2$ (Table S2). Mortality for $P_1$ and $P_2$, and *rgr* just for $P_{2,}$ were also negatively correlated ($P \leq 0.05$) with HET effects on growth in the same periods. These growth rates were limited to ones that were valid, however. The negative correlations between *rgr* and CON effects on growth in $P_1$ were reaffirmed in more limited comparisons involving 22 and 32 species out of the 38, from two previously published analyses for the same plots, and also for $P_{2a}$ (the first part of $P_2$, 1996-1999; [19, 20; main text]), the short span that included the 1998 ENSO drought, yet not in the $P_{2b}$ following (1999-2001). With one exception, other variables were not significantly correlated; and all correlations between these variables small-tree population variables and CON and HET effects on survival, and differences in CON and HET effects on growth, were non-significant. Reactivity measured the relative change in *rgr* between periods or part periods, for instance between $P_{2a}$ to $P_{2b}$ (as *rea*), or how the *rgr* in these part periods (as $rea_1$ and $rea_2$) differed from the average *rgr* of the periods before ($P_1$) and after ($P_{2b}$) the ENSO disturbance (see Table S2). Surprisingly, *rea* was just weakly positively correlated with CON and HET effects on growth in $P_1$, and not in $P_2$ — the period in which the ENSO disturbance itself occurred. A PCA first axis though the community-level graph of difference in CON effects on growth versus CON effects on survival (Fig. 2b, main text) was not significantly correlated with any of the dynamics variables or reactivity.

Table S2**.** Correlations (Pearson's *r*) between Danum plots' population dynamics variables and CON and HET effects on growth in periods 1 and 2 ($P_1$, $P_2$) from the 'larsm/reloc/crown' spatial nearest-neighbour model: (a) annualized mortality ($m_a$) and recruitment ($r_a$) and mean relative stem growth rate (*rgr*) for small trees [10 -< 50 cm gbh] in the full periods for which the effects estimates corresponded (D. M. Newbery, M. Lingenfelder & C. E. Ridsdale unpubl. data); (b) $m_a$ and *rgr*, in two size classes of small tree as shown, for respectively the 22 species analyzed by Lingenfelder and Newbery [19, main text], and 32 of the species matching those of the present study analyzed by Newbery et al. [20, main text], in period 1 and in the first two of three parts of period 2 [a, 1996-1999; b, 1999-2001; c, 2001-2007], and part-period dynamics correlated with their corresponding full-period estimates; and (c) reactivity, in terms of *rgr* (*rea*, partitioned into $rea_1$ and $rea_2$), as calculated in Newbery et al.[20, main text].

| (a) period | effect | $\ln(m_a)$ | $\ln(r_a)$ | $\ln(rgr)$ |
|---|---|---|---|---|
| $P_1$ | CON | −0.454** | −0.425** | −0.482** |
| | HET | −0.339* | $-0.200^{ns}$ | $-0.234^{ns}$ |
| $P_2$ | CON | $-0.236^{ns}$ | $-0.133^{ns}$ | −0.420** |
| | HET | −0.340* | $-0.284^{o}$ | −0.387* |

| (b) | size: | 10 - < 50 cm | | 12.5 - < 50 cm |
|---|---|---|---|---|
| period | effect | $\ln(m_a)$ | $\ln(rgr)$ | $\ln(rgr)$ |
| $P_1$ | CON | $0.228^{ns}$ | −0.438* | −0.364* |
| | HET | −0.496* | $-0.373^{o}$ | $-0.071^{ns}$ |
| $P_{2a}$ | CON | $0.046^{ns}$ | $-0.192^{ns}$ | −0.384* |
| | HET | $-0.098^{ns}$ | $0.201^{ns}$ | $-0.129^{ns}$ |
| $P_{2b}$ | CON | $0.331^{ns}$ | $0.241^{ns}$ | $-0.296^{o}$ |
| | HET | $-0.024^{ns}$ | $0.104^{ns}$ | $-0.253^{ns}$ |

| (c) period | effect | $rea^a$ | $rea_1$ | $rea_2$ |
|---|---|---|---|---|
| $P_1$ | CON | 0.389* | $0.296^{o}$ | $0.148^{ns}$ |
| | HET | 0.414* | $0.292^{ns}$ | $0.182^{ns}$ |
| $P_2$ | CON | $0.195^{ns}$ | $0.215^{ns}$ | $0.005^{ns}$ |
| | HET | $-0.170^{ns}$ | $-0.261^{ns}$ | $0.073^{ns}$ |

**, $P \leq 0.01$; *, $P \leq 0.05$; $^{o}$, $P \leq 0.10$, $^{ns}$ $P > 0.05$.
$^a$, $rea = (rgr_{2b} - rgr_{2a})/mrgr_{1,2c}$; $rea_1 = (mrgr_{1,2c} - rgr_{2a})/mrgr_{1,2c}$; $rea_2 = (rgr_{2b} - mrgr_{1,2c})/mrgr_{1,2c}$.

2. *Spatial pattern*

The second-order spatial statistic 'L(r) – r' (Ripley 1977) was calculated for each of the 38 selected species, for the four combinations of plots 1 and 2 times $P_1$ and $P_2$, range r = 0 to 60 m, and for either trees of focal size (10 - < 100 cm *gbh*) or all trees (≥ 10 cm *gbh*). Most species showed very slight departures from randomness towards aggregation, more so in the range r = 40 – 60 m ($P \leq 0.05$), but differing often between plots, not periods. Five of 20 extended understorey species (OUI ≤ 30) had very strong aggregations in both plots (*Cleistanthus contractus, Dimorphocalyx muricatus, Lophopetalum beccarianum, Mallotus penangensis, M. stipularis*; see Appendix 1: Table S1 for full names). CON and HET effects on growth, and on survival, in $P_1$ and $P_2$ (eight *t*-tests), showed no significant differences in their means between aggregated and non-aggregated species, however ($P$ = 0.17 – 0.76). Among the 18 intermediate-overstorey species (OUI > 30), just two were clearly aggregated (*Shorea fallax, S. pilosa*), four others more weakly so and not always consistently so across both plots (*Lithocarpus gracilis, Pentace laxiflora, Parashorea malaanonan, Syzygium elopurae*). Among the dipterocarps, comparing the two strongly aggregated dipterocarp species (*S. fallax, S. pilosa*) with the four others weakly or non-aggregated (*P. malaanonan, S. johorensis, S. parvifolia, S. pauciflora*), CON and HET effects (again the eight *t*-tests), showed no significant differences in their means ($P$ = 0.09 – 0.89). The more outlying species' points in Figs 2b and 3b (main text), showed no particular associations with degree of aggregation. Taken together these comparisons provide little support for spatial patterning explaining the community-level relationships.

Reference: Ripley BD. Modelling spatial patterns. Journal of the Royal Statistical Society B. 1977; 39:172-212.

Fig. S1. Spatial pattern analysis of the 38 species used in the nearest-neighbour modelling (on the following pages).

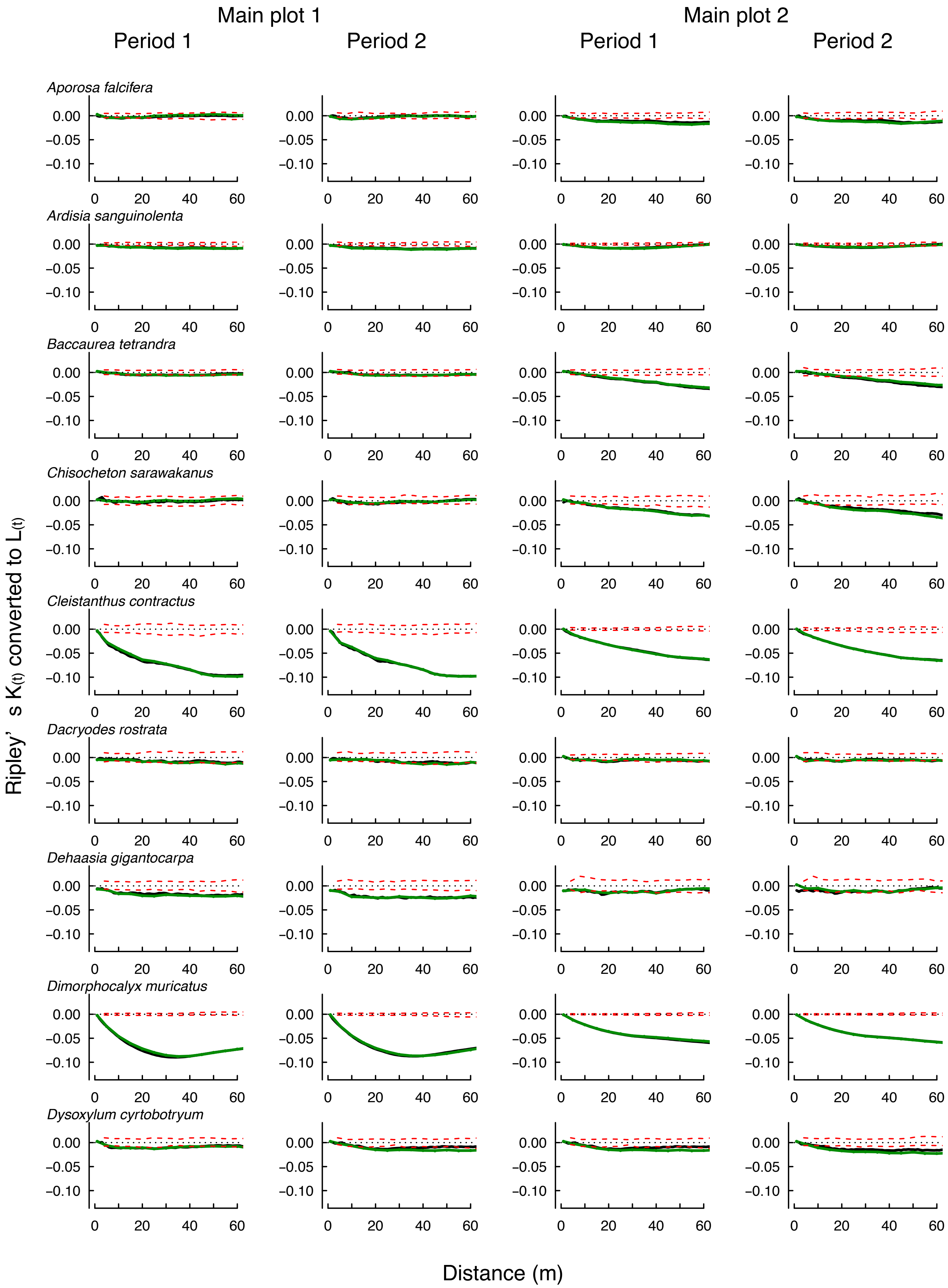

Main plot 1
Main plot 2
Period 1
Period 2
Period 1
Period 2
Aporosa falcifera
Ardisia sanguinolenta
Baccaurea tetrandra
Chisocheton sarawakanus
Cleistanthus contractus
Dacryodes rostrata
Dehaasia gigantocarpa
Dimorphocalyx muricatus
Dysoxylum cyrtobotryum
0.00
−0.05
−0.10
0
20
40
60
Ripley's K(t) converted to L(t)
Distance (m)

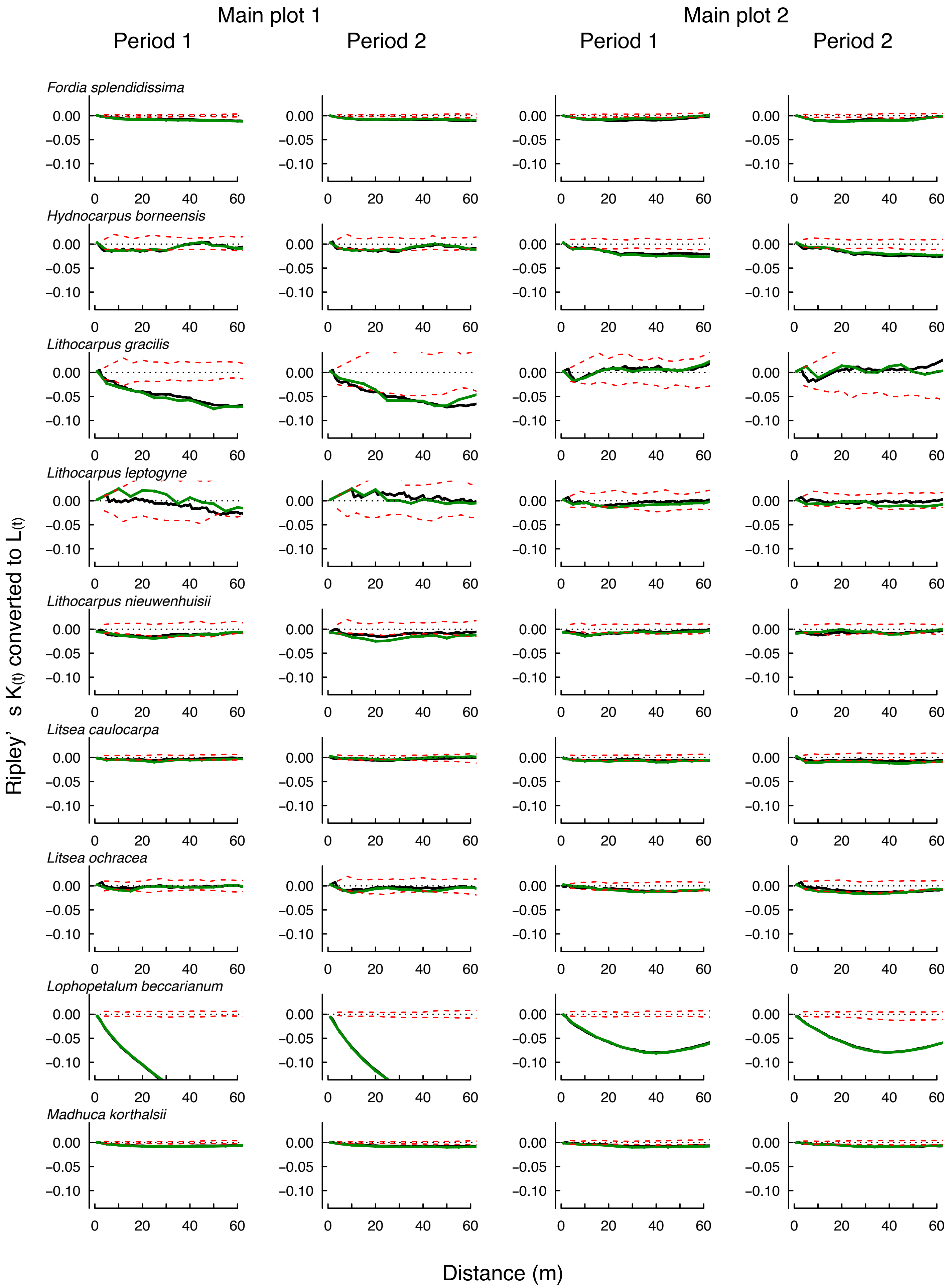

Main plot 1
Main plot 2
Period 1
Period 2
Period 1
Period 2
Fordia splendidissima
Hydnocarpus borneensis
Lithocarpus gracilis
Lithocarpus leptogyne
Lithocarpus nieuwenhuisii
Litsea caulocarpa
Litsea ochracea
Lophopetalum beccarianum
Madhuca korthalsii
0.00
−0.05
−0.10
0
20
40
60
Ripley's K(t) converted to L(t)
Distance (m)

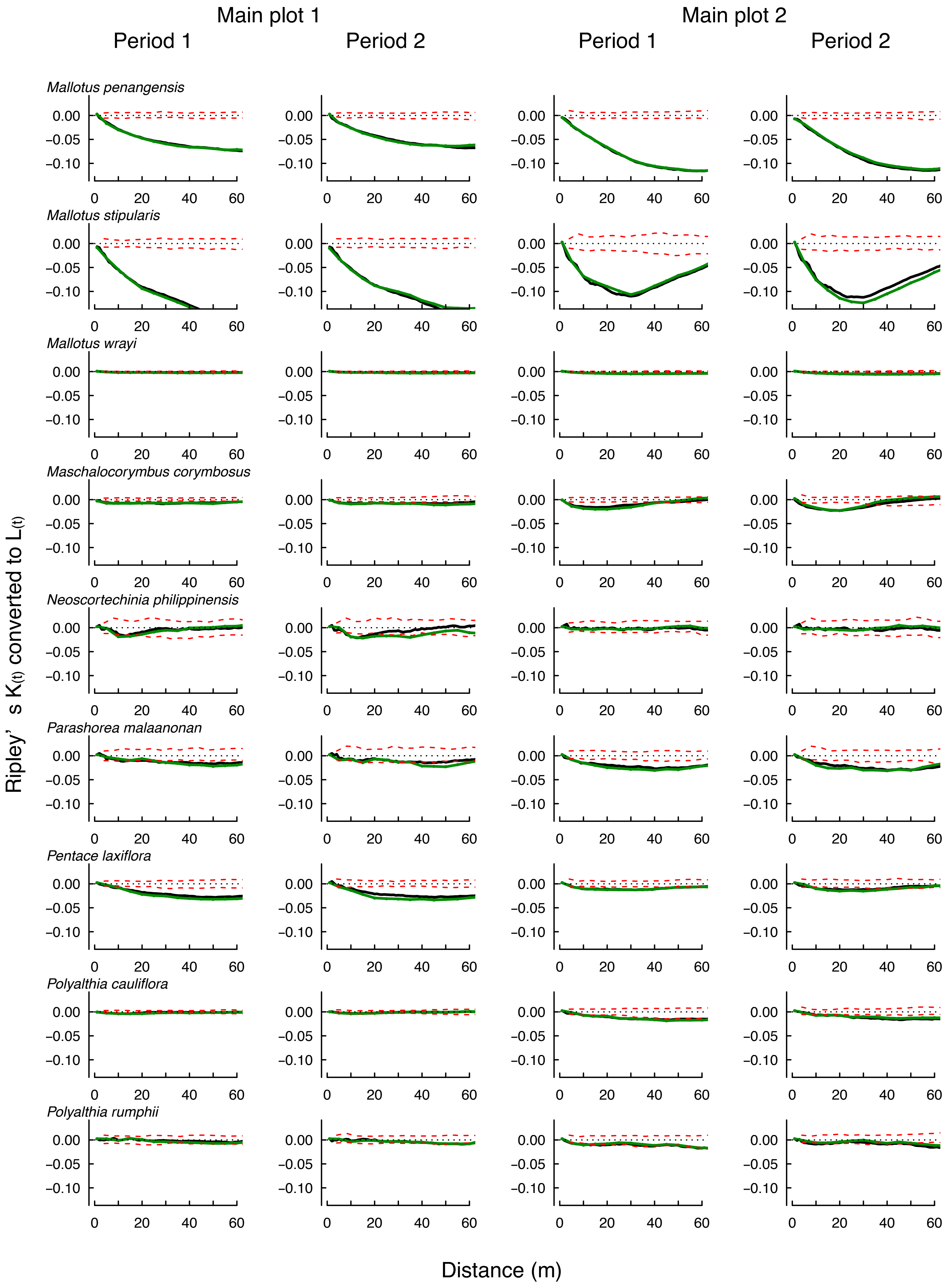

Main plot 1
Main plot 2
Period 1
Period 2
Period 1
Period 2
Mallotus penangensis
Mallotus stipularis
Mallotus wrayi
Maschalocorymbus corymbosus
Neoscortechinia philippinensis
Parashorea malaanonan
Pentace laxiflora
Polyalthia cauliflora
Polyalthia rumphii
0.00
−0.05
−0.10
0
20
40
60
Ripley’ s K(t) converted to L(t)
Distance (m)

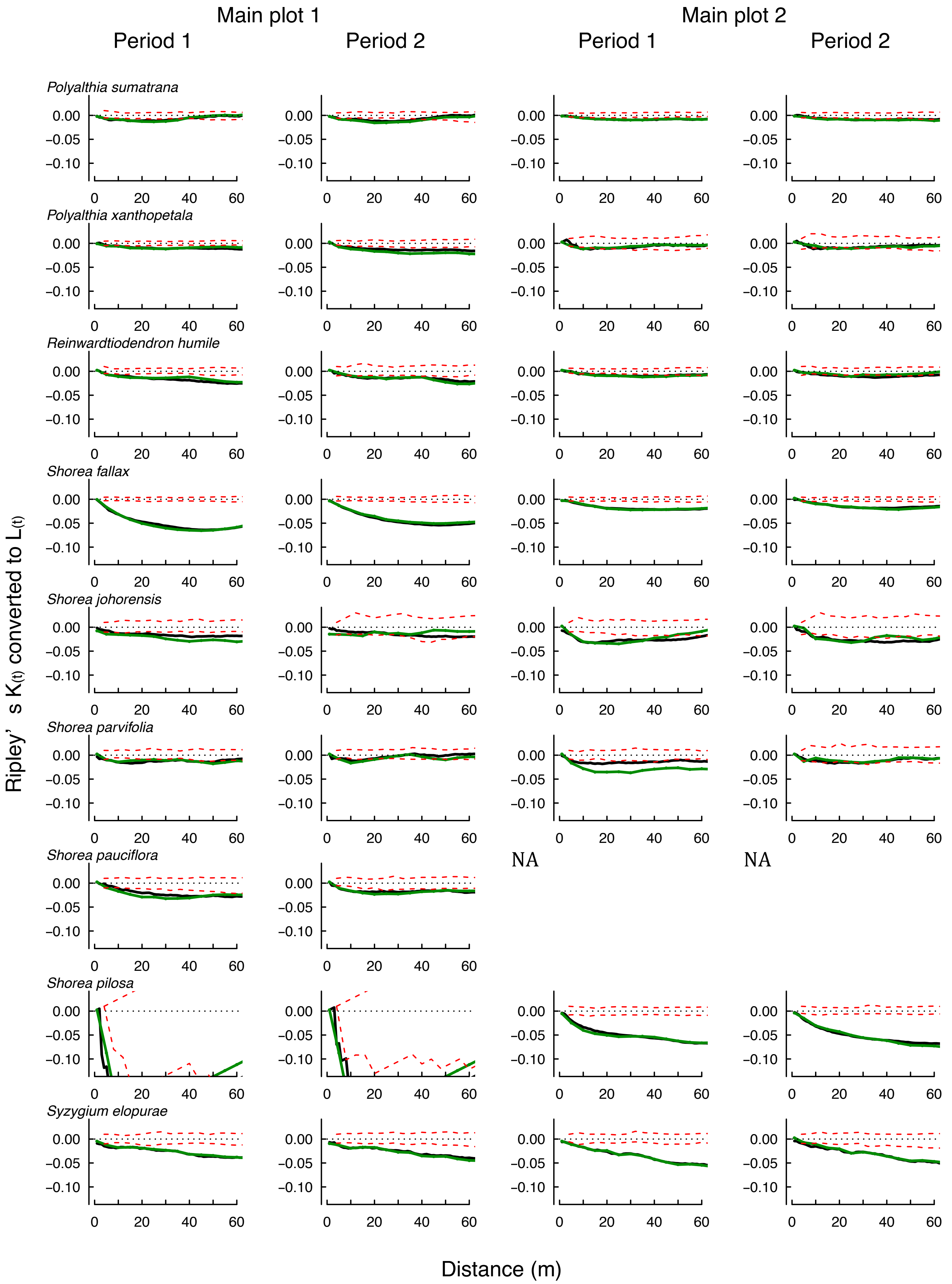

Main plot 1
Main plot 2
Period 1
Period 2
Period 1
Period 2
Polyalthia sumatrana
Polyalthia xanthopetala
Reinwardtiodendron humile
Shorea fallax
Shorea johorensis
Shorea parvifolia
Shorea pauciflora
Shorea pilosa
Syzygium elopurae
NA
NA
0.00
−0.05
−0.10
0
20
40
60
Ripley's K(t) converted to L(t)
Distance (m)

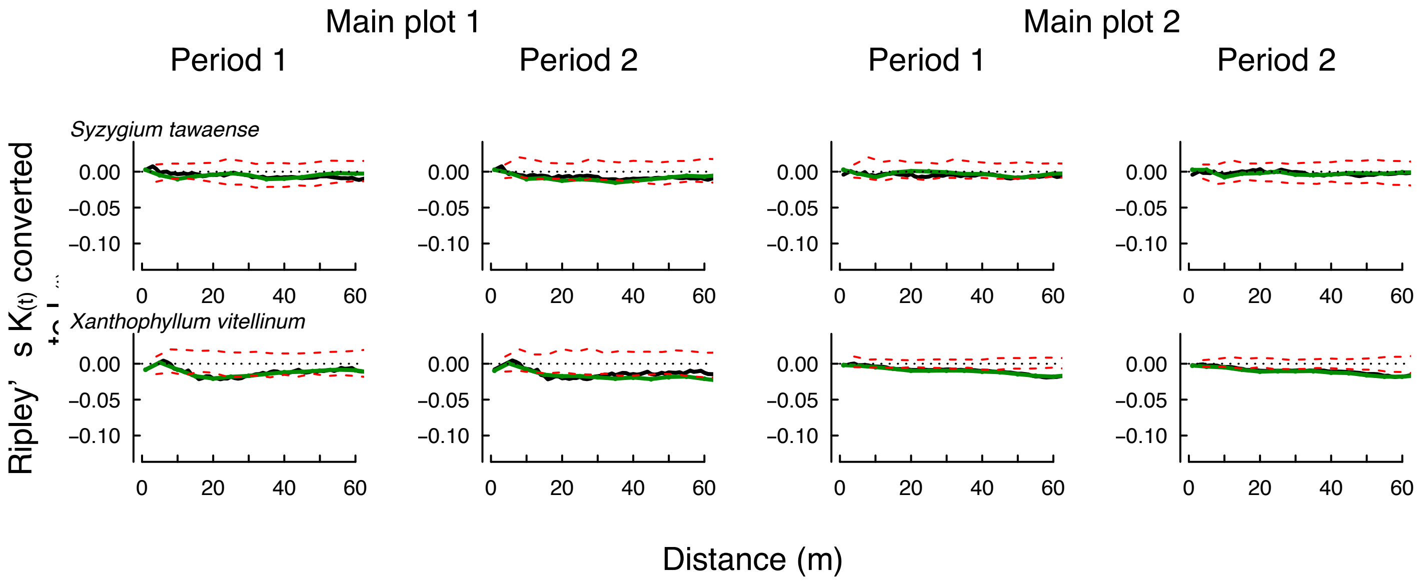

Main plot 1
Main plot 2
Period 1
Period 2
Period 1
Period 2
Syzygium tawaense
Xanthophyllum vitellinum
Ripley's K(t) converted
0.00
−0.05
−0.10
0
20
40
60
Distance (m)
