## Supplementary material for "Including tree spatial extension in the evaluation of neighbourhood competition effects in Bornean rain forest": nncrexm appendices: App7_randruns.pdf (from another version of this preprint: version 2, 2020-08-04)

Fig. S1. Single species panel-graphs (CON)

Outcome of the 48 species' analyses using 100 randomizations of tree positions (method of Newbery & Stoll, 2013), for the spatial model 'larsm/reloc/crown' [linear decay], shown in alphabetical order. (The excluded 10 species have their names in grey.) Averages of the runs are shown as means ± 3 SE's, in green if the zero-value for the variables growth ($P_1$, $P_2$ and change $P_2$-$P_1$; CON effect size) and survival ($P_1$ and $P_2$; raw CON estimate, $\beta$) was within the implied confidence limits, otherwise in red to indicate a significant difference. The estimates from the empirical data are shown as black circles, open if they lay within the limits and closed if outside of them.

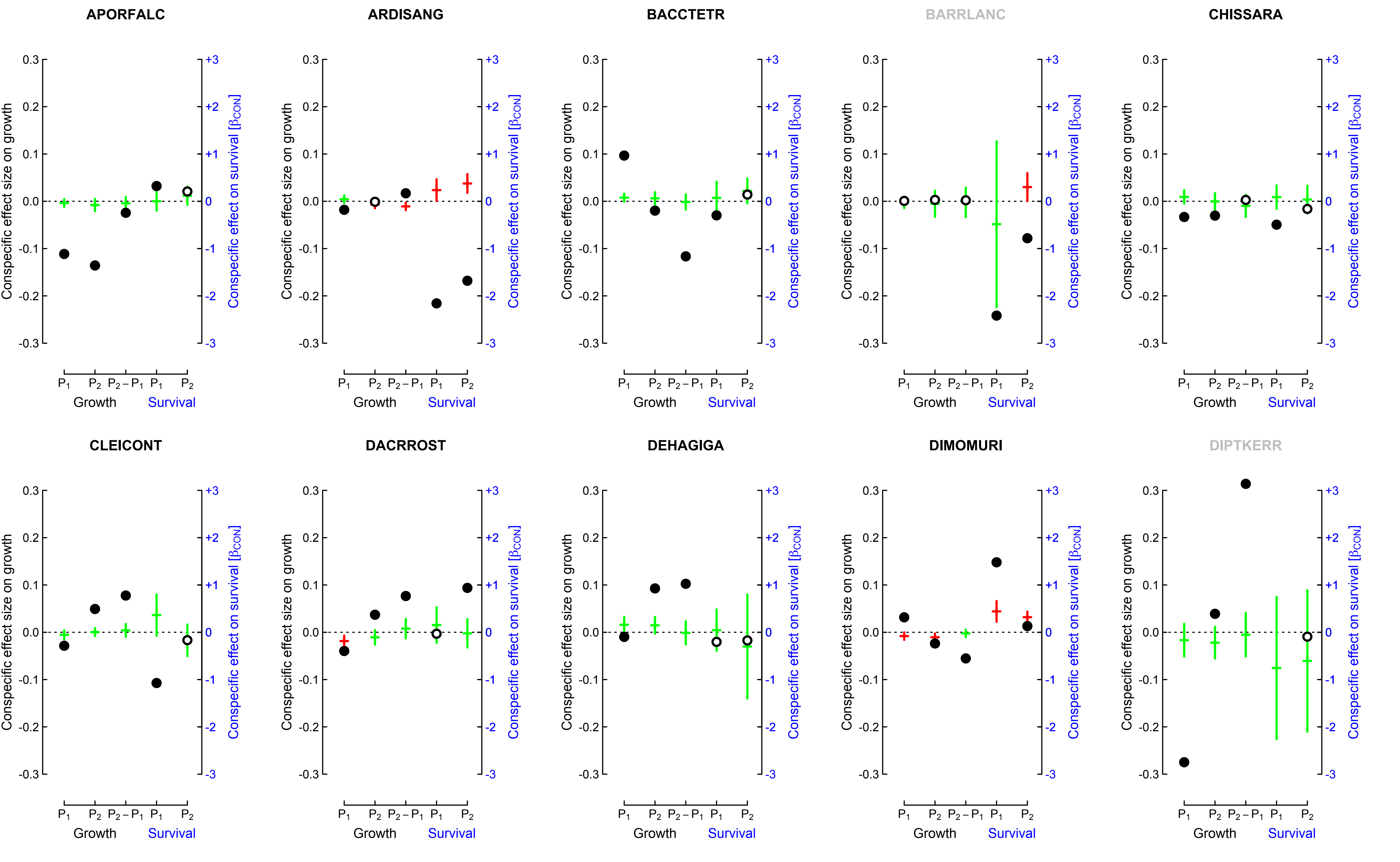
APORFALC
ARDISANG
BACCTETR
BARRLANC
CHISSARA
CLEICONT
DACRROST
DEHAGIGA
DIMOMURI
DIPTKERR
Conspecific effect size on growth
Conspecific effect on survival [βCON]
0.3
0.2
0.1
0.0
-0.1
-0.2
-0.3
+3
+2
+1
0
-1
-2
-3
P1
P2
P2 – P1
P1
P2
Growth
Survival

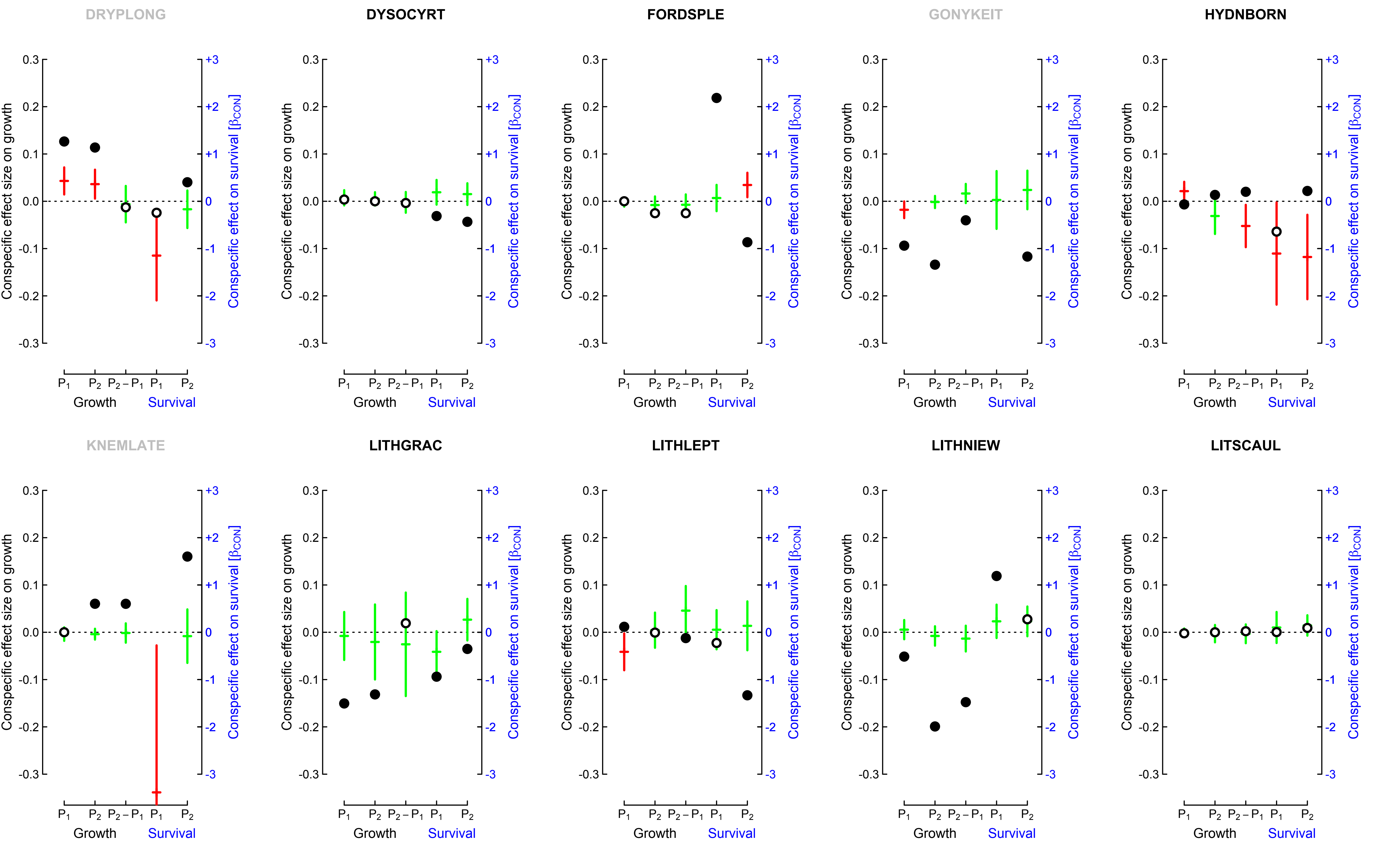
DRYPLONG
DYSOCYRT
FORDSPLE
GONYKEIT
HYDNBORN
KNEMLATE
LITHGRAC
LITHLEPT
LITHNIEW
LITSCAUL
Conspecific effect size on growth
Conspecific effect on survival [$\beta_{CON}$]
0.3
0.2
0.1
0.0
-0.1
-0.2
-0.3
+3
+2
+1
0
-1
-2
-3
$P_1$
$P_2$
$P_2 - P_1$
$P_1$
$P_2$
Growth
Survival

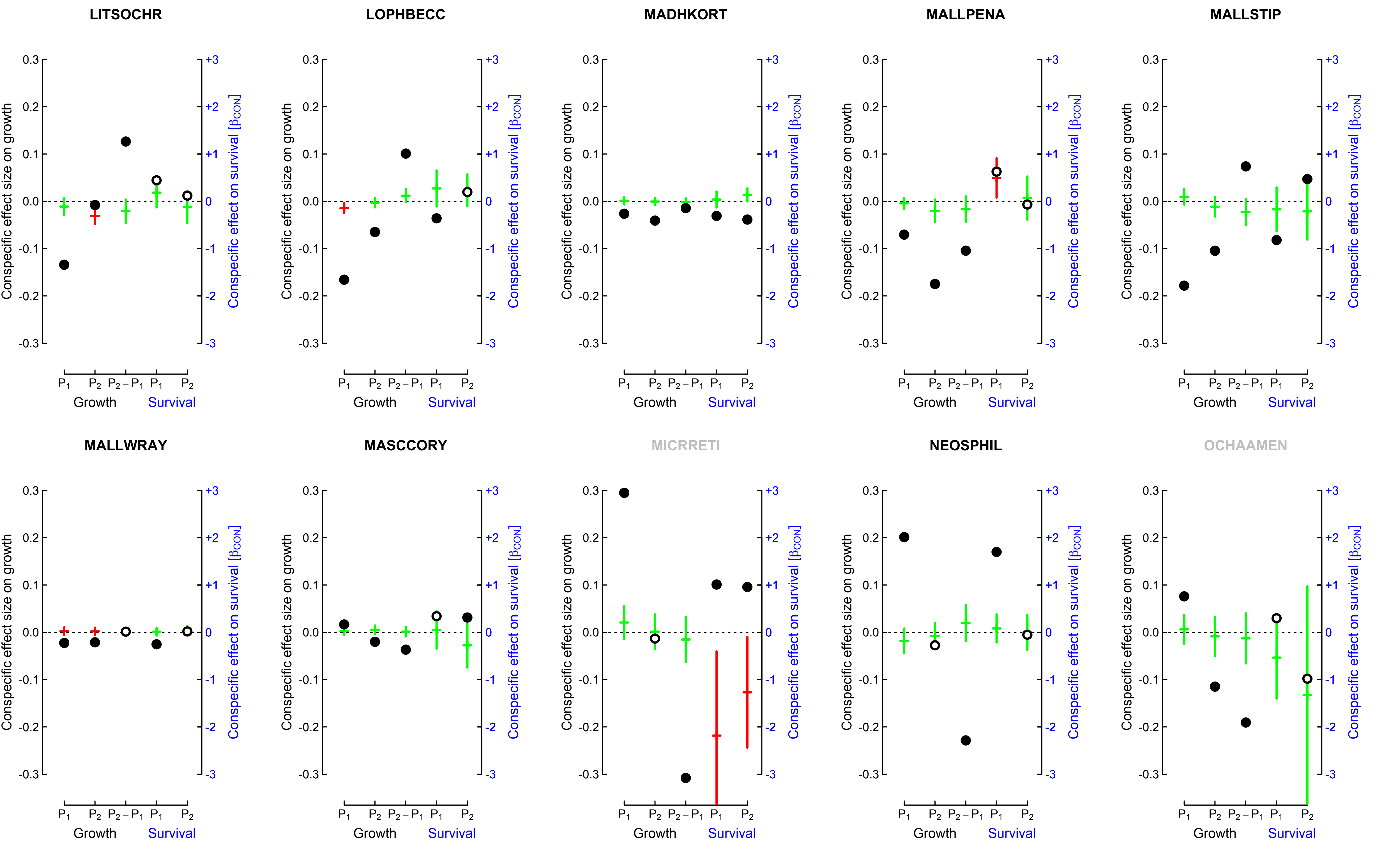

LITSOCHR
LOPHBECC
MADHKORT
MALLPENA
MALLSTIP
MALLWRAY
MASCCORY
MICRRETI
NEOSPHIL
OCHAAMEN
Conspecific effect size on growth
Conspecific effect on survival [βCON]
0.3
0.2
0.1
0.0
-0.1
-0.2
-0.3
+3
+2
+1
0
-1
-2
-3
P1
P2
P2 – P1
P1
P2
Growth
Survival

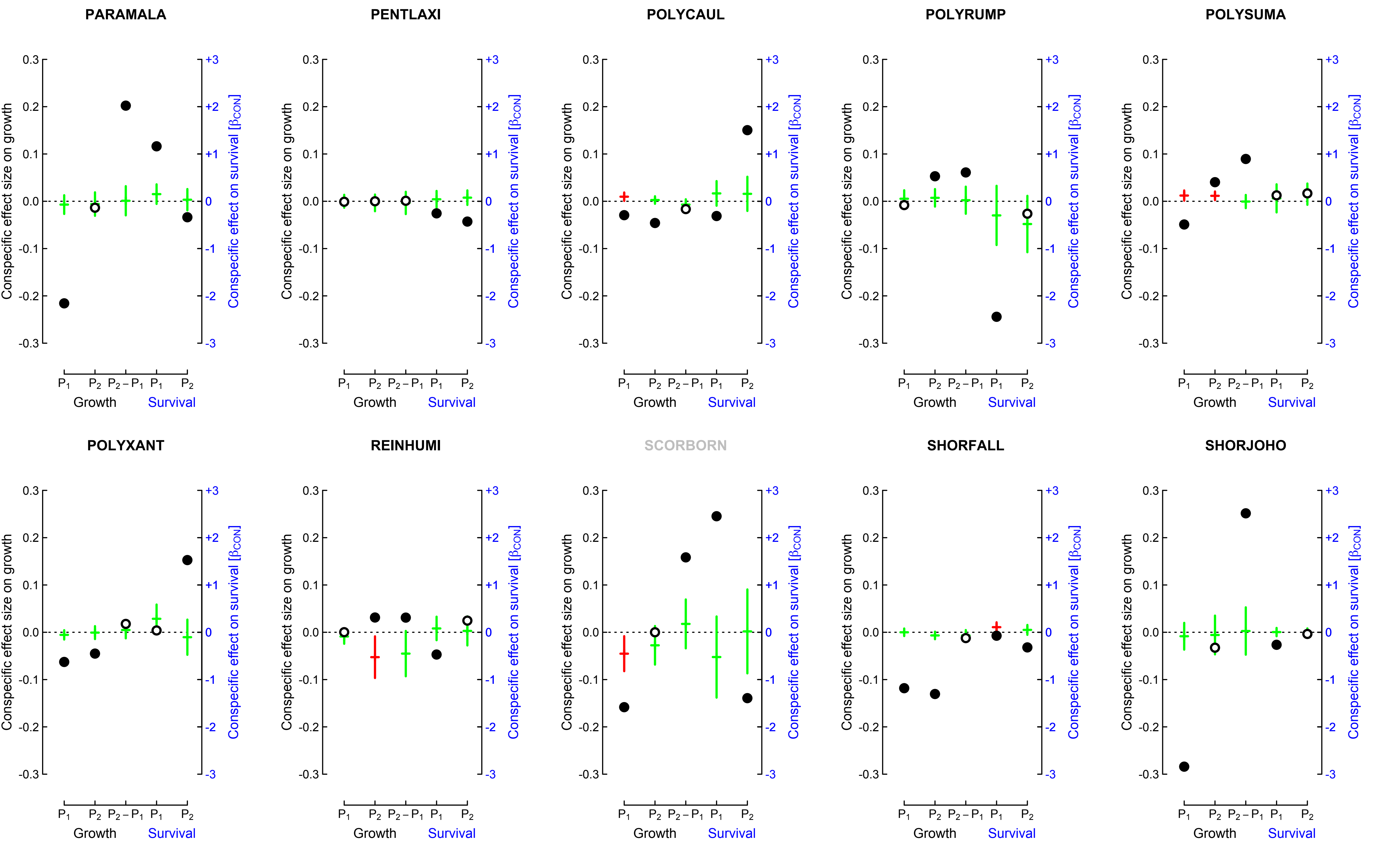
PARAMALA
PENTLAXI
POLYCAUL
POLYRUMP
POLYSUMA
POLYXANT
REINHUMI
SCORBORN
SHORFALL
SHORJOHO
Conspecific effect size on growth
Conspecific effect on survival [βCON]
P1
P2
P2 – P1
Growth
Survival

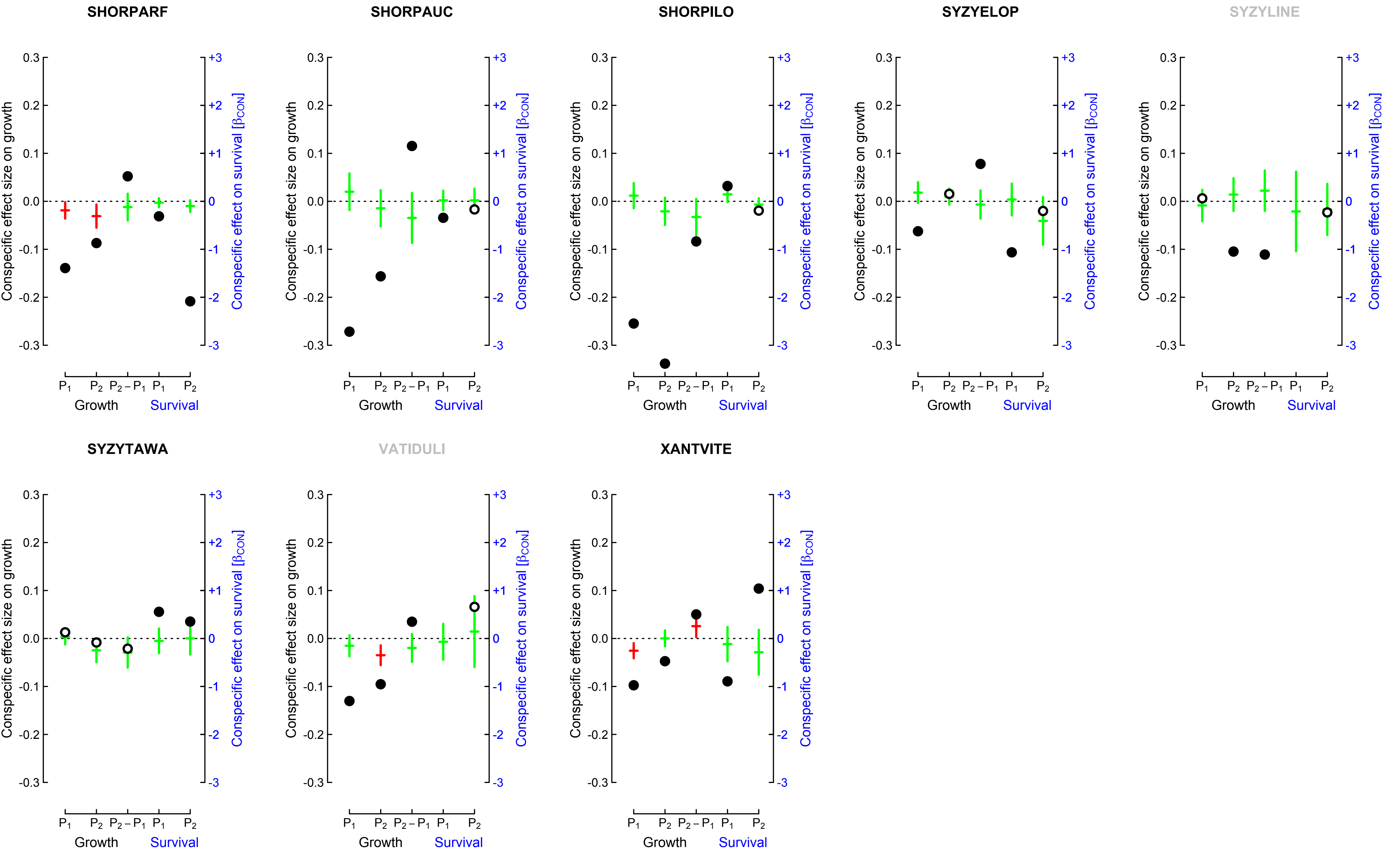
SHORPARF
SHORPAUC
SHORPILO
SYZYELOP
SYZYLINE
SYZYTAWA
VATIDULI
XANTVITE
Conspecific effect size on growth
Conspecific effect on survival [$\beta_{CON}$]
0.3
0.2
0.1
0.0
-0.1
-0.2
-0.3
+3
+2
+1
0
-1
-2
-3
$P_1$ $P_2$ $P_2 - P_1$ $P_1$ $P_2$
Growth
Survival

Fig. S2. Single species panel-graphs (HET)

Outcome of the 48 species' analyses using 100 randomizations of tree positions (method of Newbery & Stoll, 2013), for the spatial model 'larsm/reloc/crown' [linear decay], shown in alphabetical order. (The excluded 10 species have their names in grey.) Averages of the runs are shown as means ± 3 SE's, in green if the zero-value for the variables growth ($P_1$, $P_2$ and change $P_2$-$P_1$; HET effect size) and survival ($P_1$ and $P_2$; raw HET estimate, β) was within the implied confidence limits, otherwise in red to indicate a significant difference. The estimates from the empirical data are shown as black circles, open if they lay within the limits and closed if outside of them.

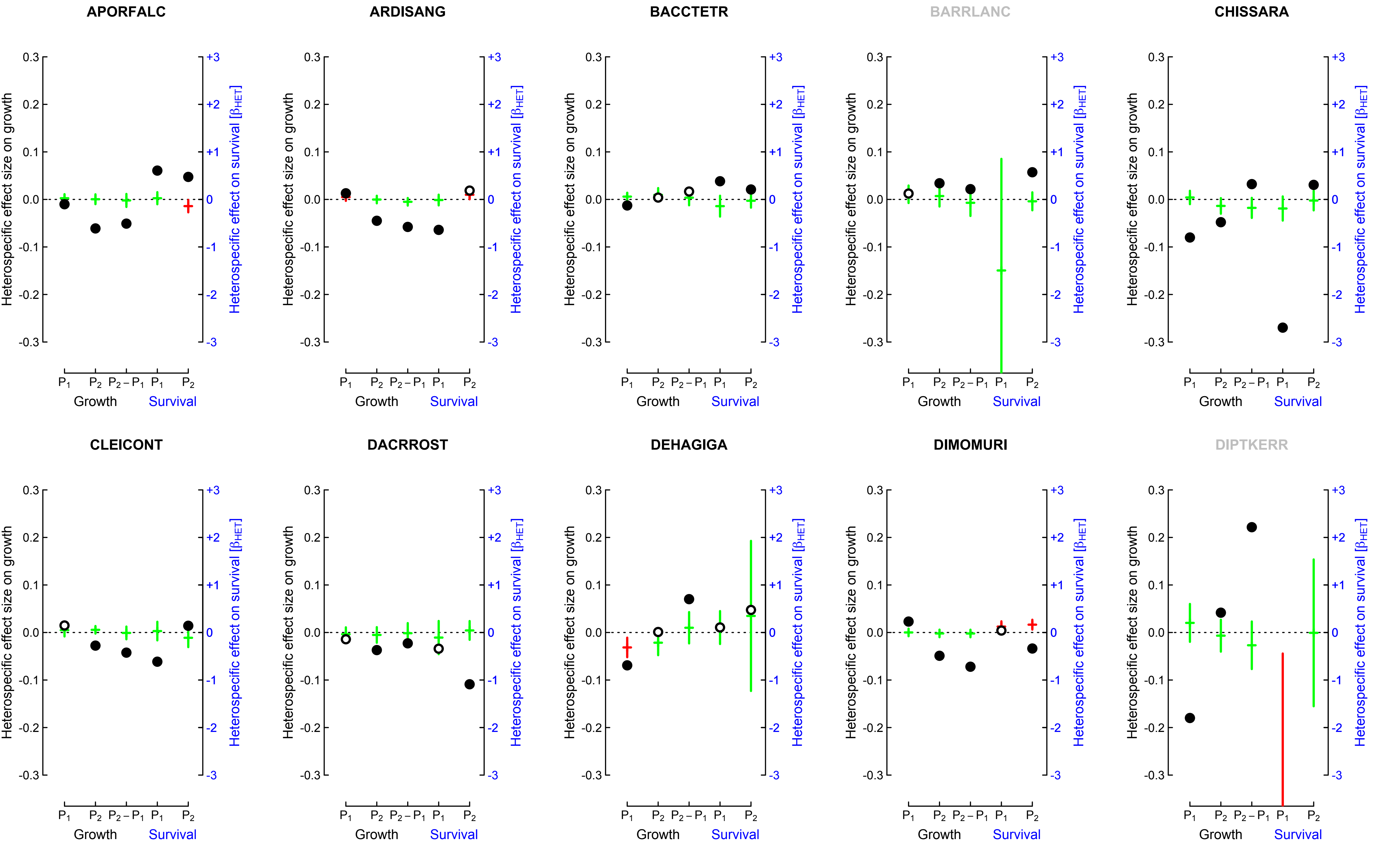
APORFALC
ARDISANG
BACCTETR
BARRLANC
CHISSARA
CLEICONT
DACRROST
DEHAGIGA
DIMOMURI
DIPTKERR
Heterospecific effect size on growth
Heterospecific effect on survival [$\beta_{HET}$]
0.3
0.2
0.1
0.0
-0.1
-0.2
-0.3
+3
+2
+1
0
-1
-2
-3
$P_1$ $P_2$ $P_2 - P_1$ $P_1$ $P_2$
Growth
Survival

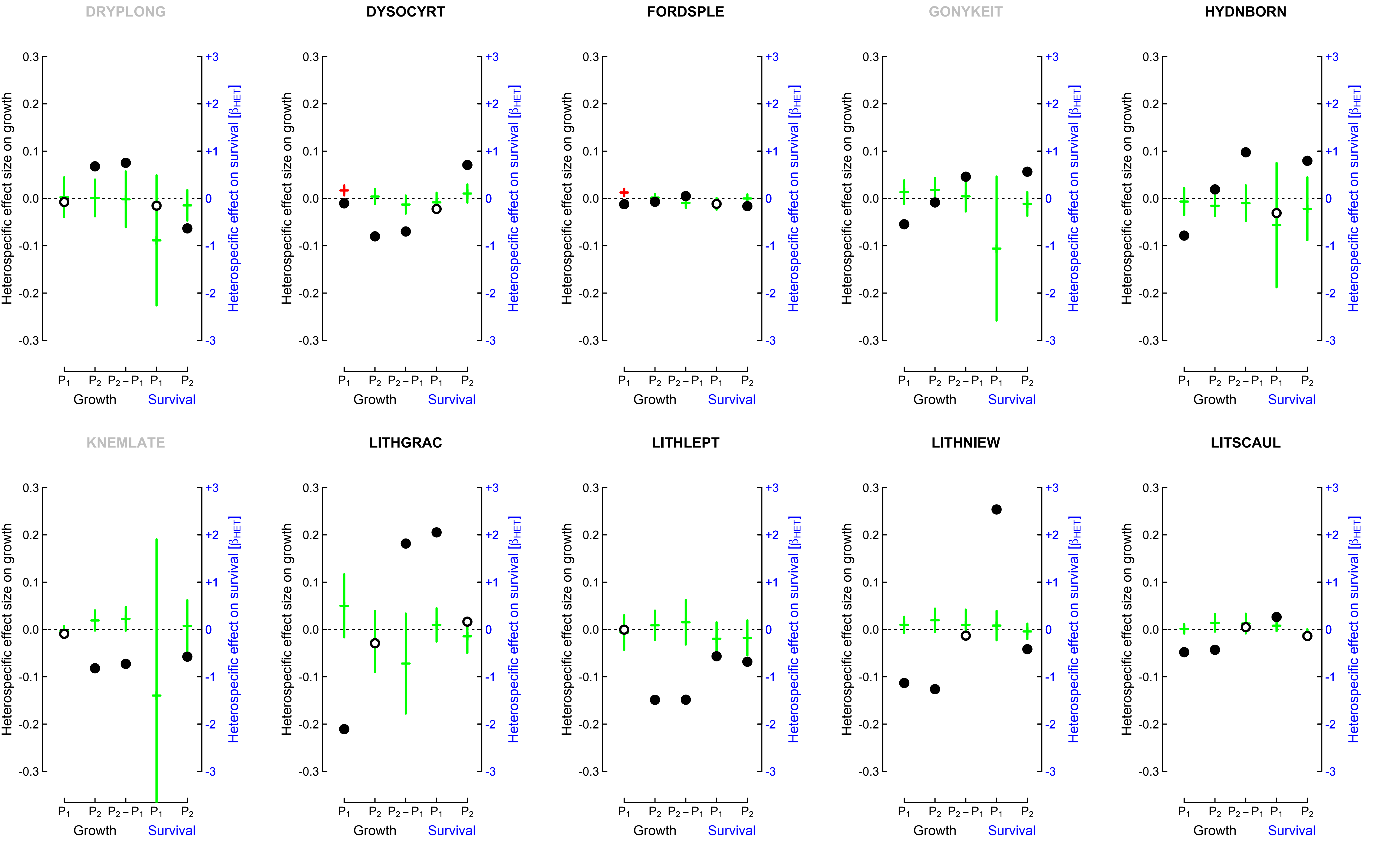
DRYPLONG
DYSOCYRT
FORDSPLE
GONYKEIT
HYDNBORN
KNEMLATE
LITHGRAC
LITHLEPT
LITHNIEW
LITSCAUL
Heterospecific effect size on growth
Heterospecific effect on survival [β HET]
0.3
0.2
0.1
0.0
-0.1
-0.2
-0.3
+3
+2
+1
0
-1
-2
-3
P1
P2
P2 – P1
P1
P2
Growth
Survival

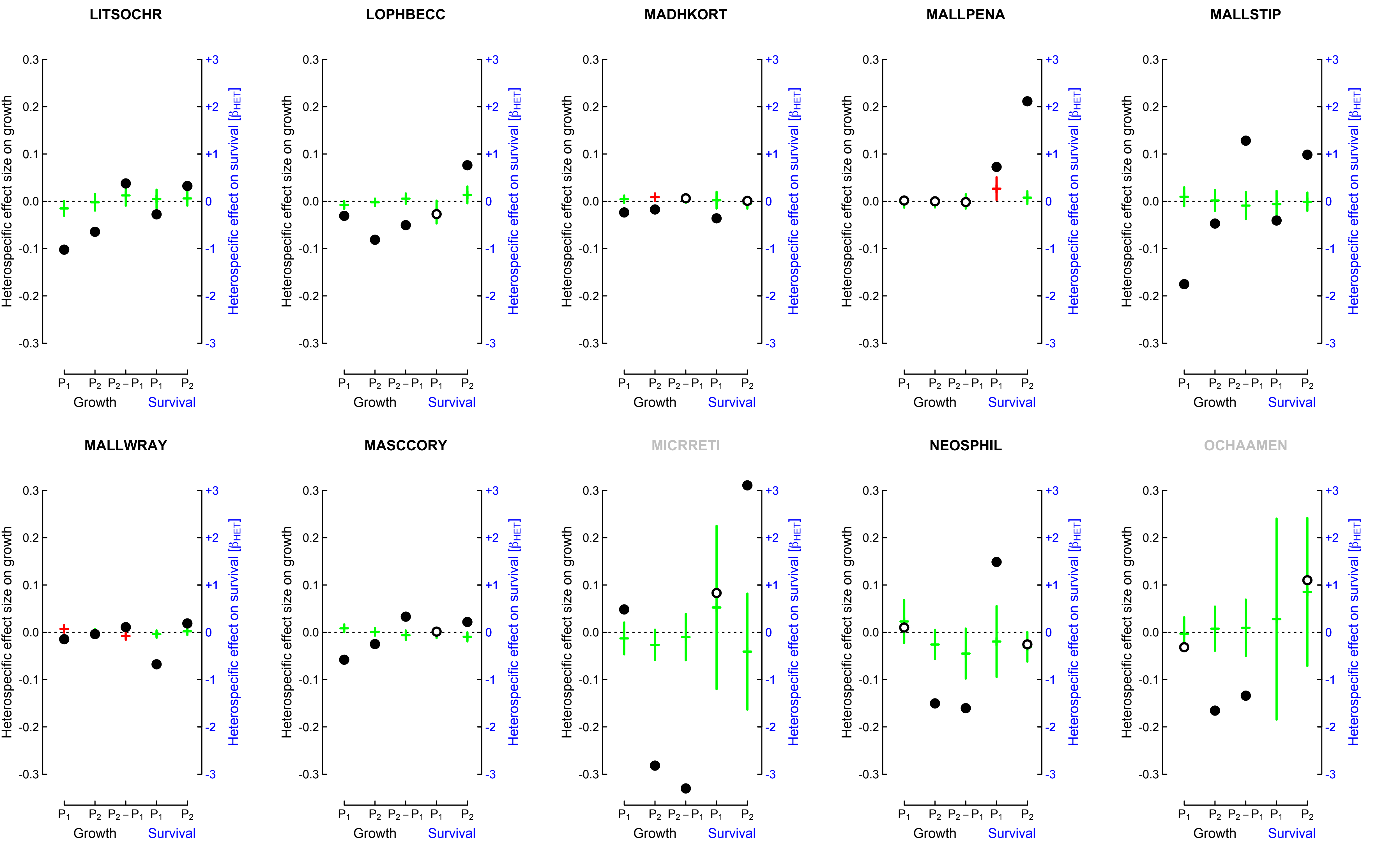
LITSOCHR
LOPHBECC
MADHKORT
MALLPENA
MALLSTIP
MALLWRAY
MASCCORY
MICRRETI
NEOSPHIL
OCHAAMEN
Heterospecific effect size on growth
Heterospecific effect on survival [$\beta_{HET}$]
0.3
0.2
0.1
0.0
-0.1
-0.2
-0.3
+3
+2
+1
0
-1
-2
-3
$P_1$
$P_2$
$P_2 - P_1$
$P_1$
$P_2$
Growth
Survival

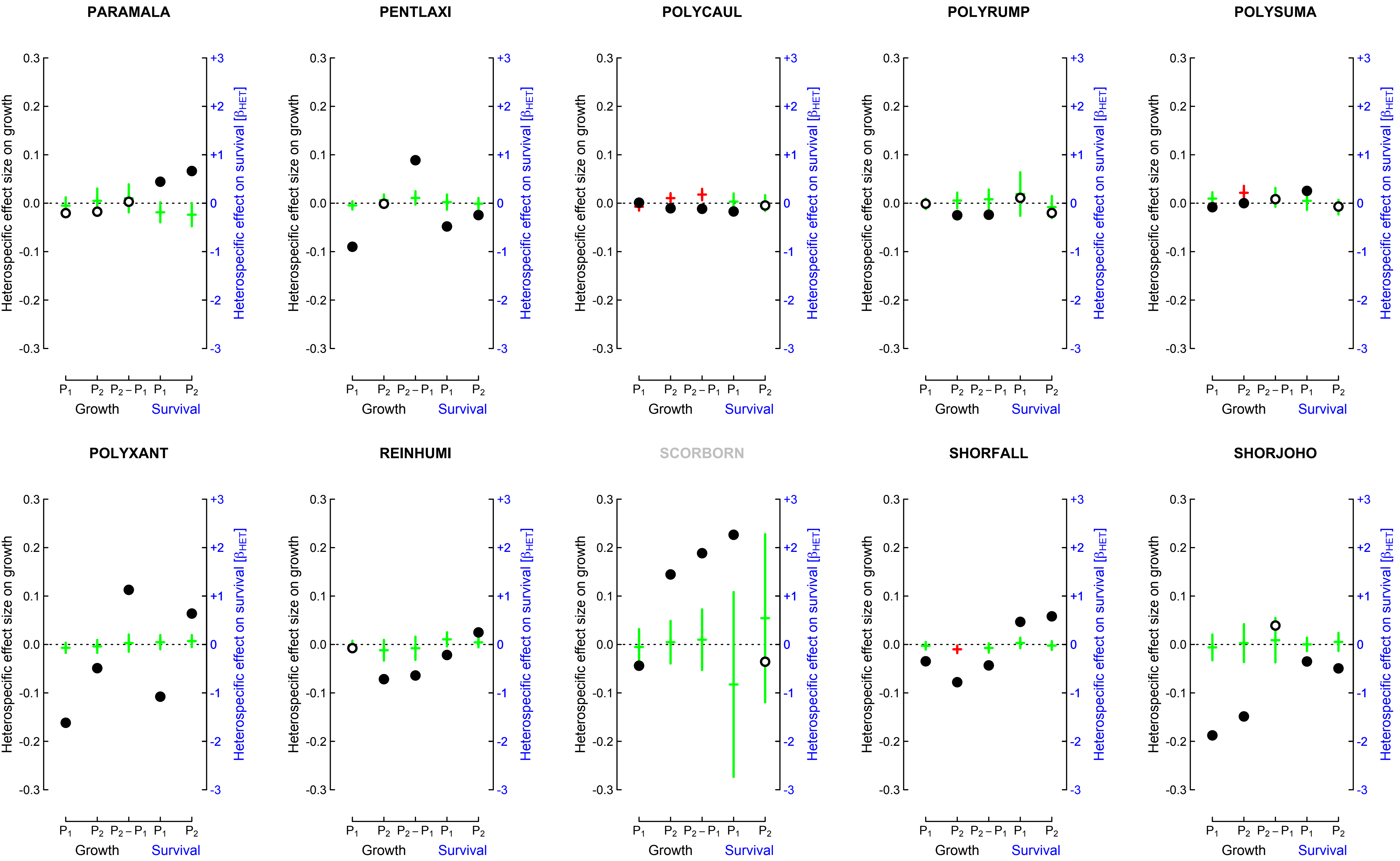

PARAMALA
PENTLAXI
POLYCAUL
POLYRUMP
POLYSUMA
POLYXANT
REINHUMI
SCORBORN
SHORFALL
SHORJOHO
Heterospecific effect size on growth
Heterospecific effect on survival [$\beta_{HET}$]
0.3
0.2
0.1
0.0
-0.1
-0.2
-0.3
+3
+2
+1
0
-1
-2
-3
$P_1$ $P_2$ $P_2 - P_1$ $P_1$ $P_2$
Growth
Survival

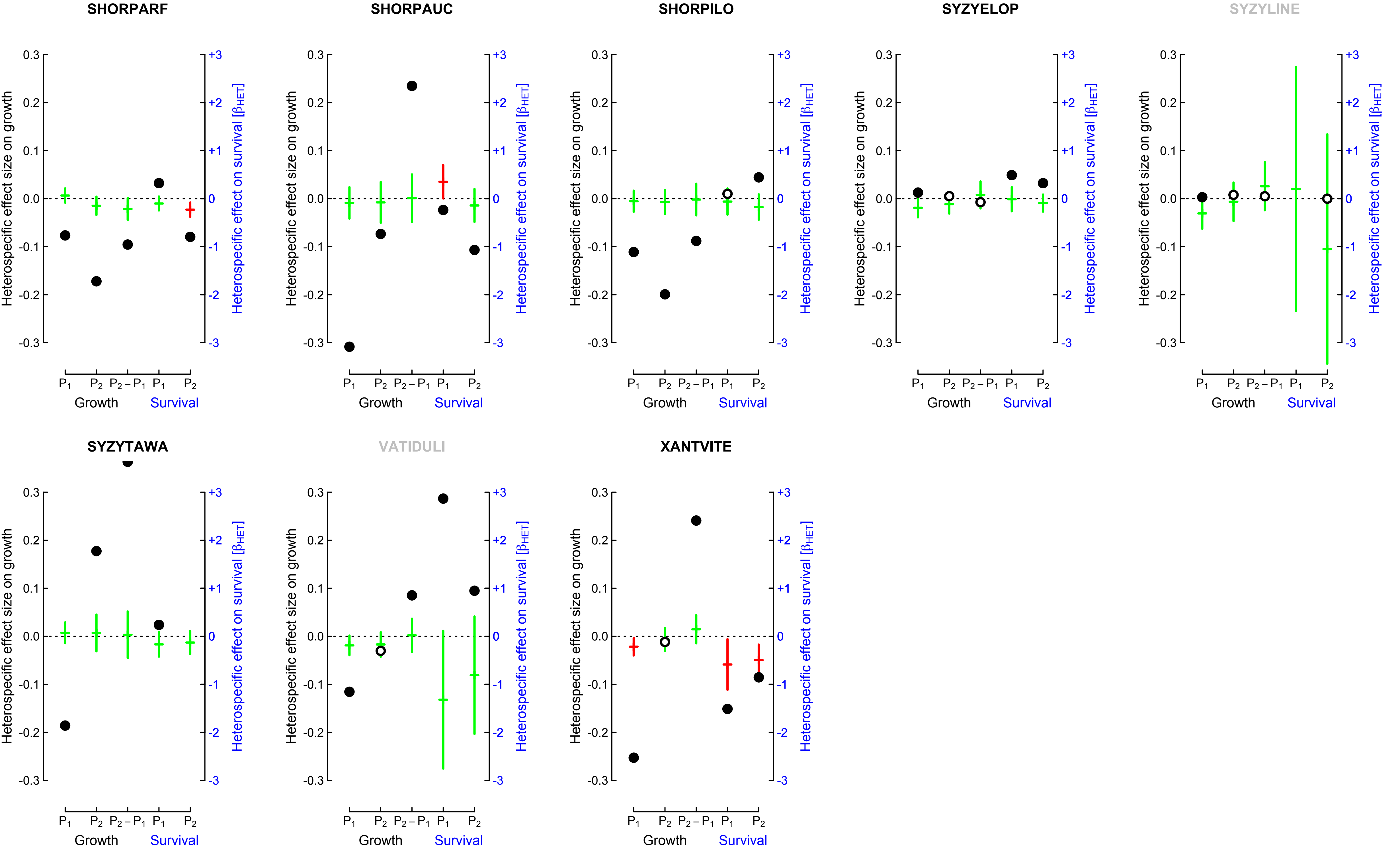
SHORPARF
SHORPAUC
SHORPILO
SYZYELOP
SYZYLINE
SYZYTAWA
VATIDULI
XANTVITE
Heterospecific effect size on growth
Heterospecific effect on survival [$\beta_{HET}$]
$P_1$
$P_2$
$P_2 - P_1$
Growth
Survival

Figure S3. The change in effect size on growth between $P_1$ and $P_2$ versus effect size on survival in $P_1$, as fitted regression lines from the 100 randomizations of tree positions (method of Newbery & Stoll, 2013), for the spatial model ‘larsm/reloc/crown’ [lin decay]: (a) CON effects based using all 38 selected species, (b) CON effects limited to the 24 species where *P* was < 0.1 for survival effects in $P_{1;}$ and (c) and (d) correspondingly the same for HET effects. Lines with positive and negative slopes are shown in green and blue respectively: continuous lines are significant at $P \geq 0.05$, dashed lines are not significant. The line from the regression using the empirical data is in red (continuous), as also shown for (a) on the last line of Table 5a and on Fig. 2b), for (b) on the last line of Table 5b, and for (c) on Fig. 3b.

**(a) CON, 38 spp.**

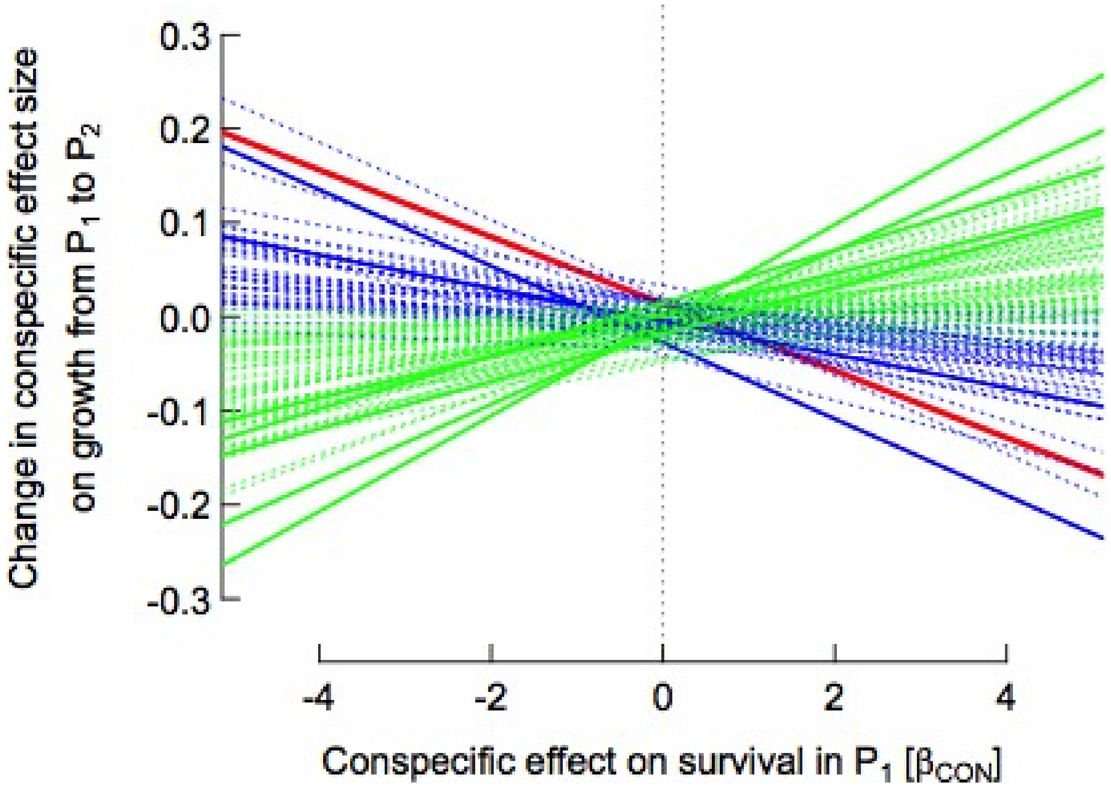

Change in conspecific effect size on growth from $P_1$ to $P_2$
0.3
0.2
0.1
0.0
-0.1
-0.2
-0.3
-4
-2
0
2
4
Conspecific effect on survival in $P_1$ [$\beta_{CON}$]

**(b) CON, 24 spp.**

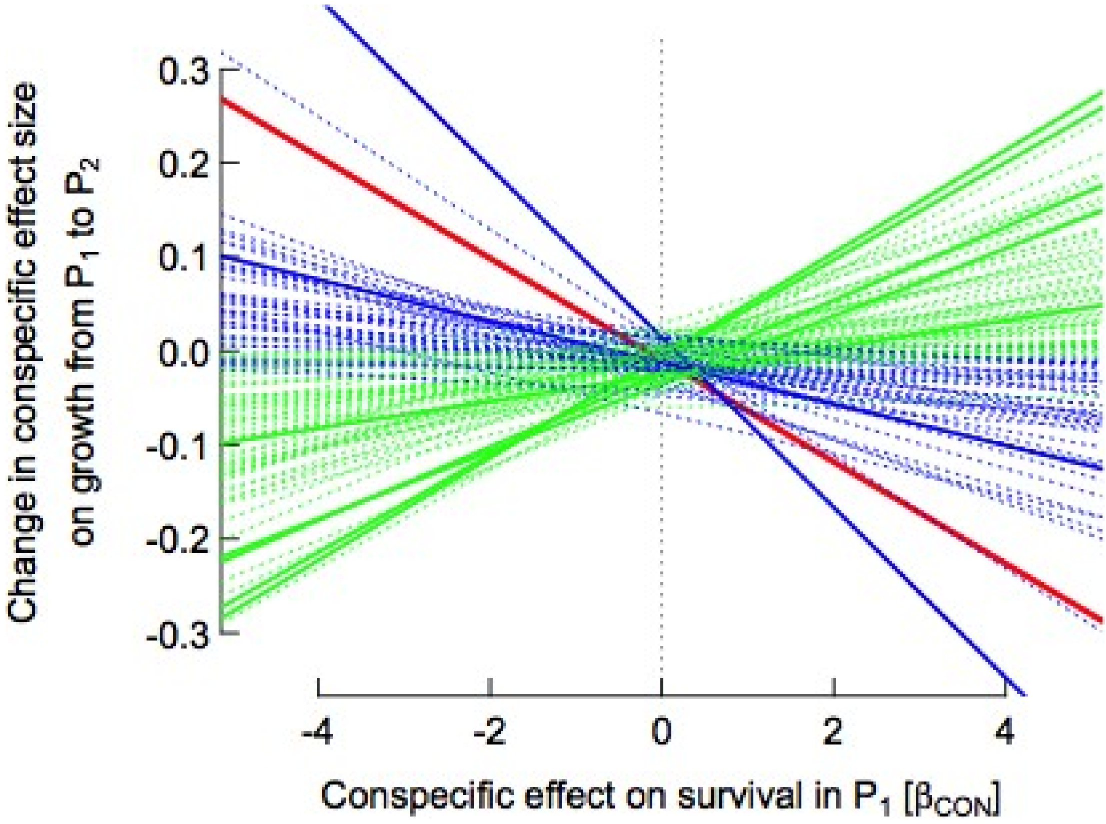

Change in conspecific effect size on growth from $P_1$ to $P_2$
0.3
0.2
0.1
0.0
-0.1
-0.2
-0.3
-4
-2
0
2
4
Conspecific effect on survival in $P_1$ [$\beta_{CON}$]

**(c) HET, 38 spp.**

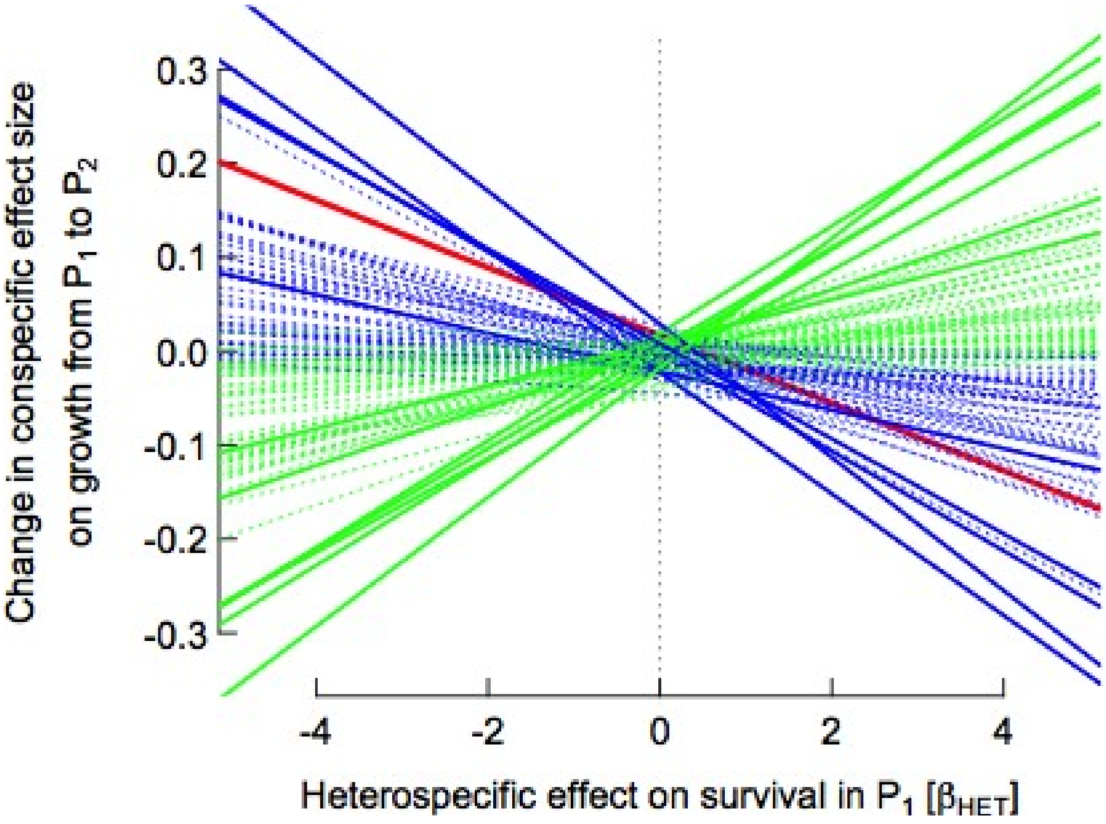

0.3
0.2
0.1
0.0
-0.1
-0.2
-0.3
Change in conspecific effect size on growth from $P_1$ to $P_2$
-4
-2
0
2
4
Heterospecific effect on survival in $P_1$ [$\beta_{HET}$]

(c) **HET, 24 spp.**

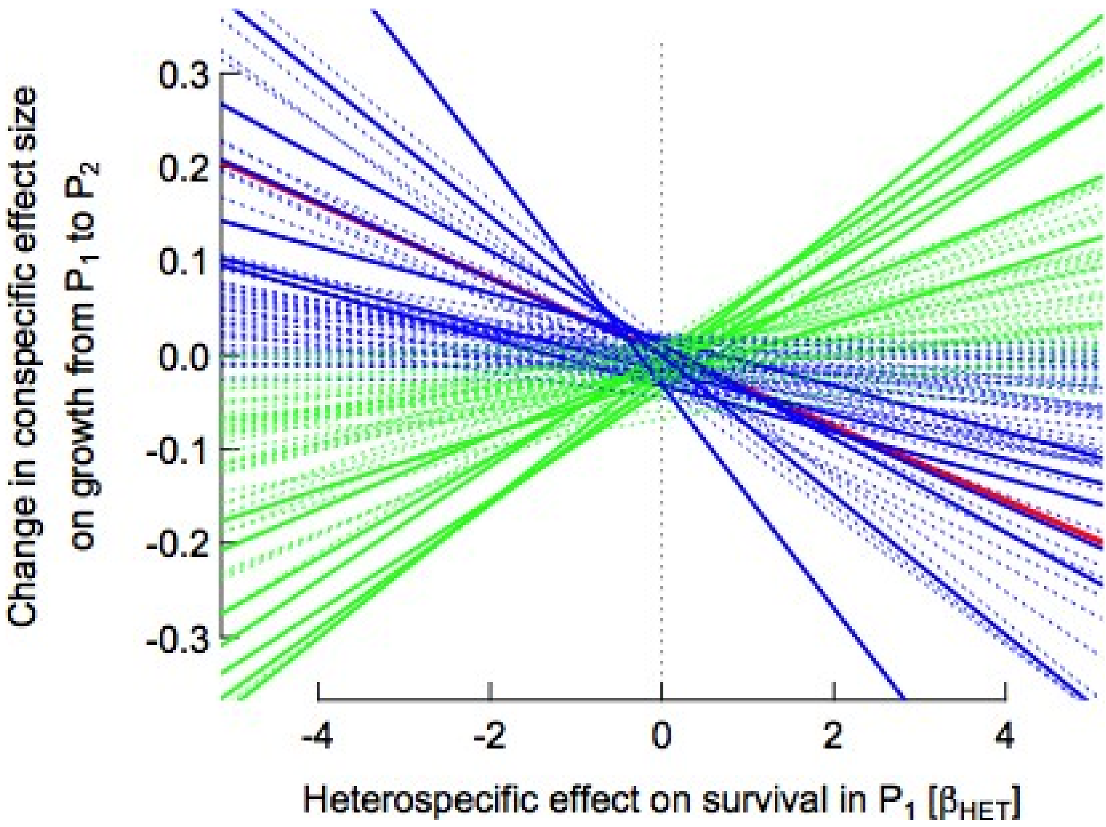

Change in conspecific effect size on growth from P1 to P2
0.3
0.2
0.1
0.0
-0.1
-0.2
-0.3
-4
-2
0
2
4
Heterospecific effect on survival in P1 [βHET]
